# CRISPR-dPCR: A unified multiplex digital PCR framework for quantitative characterization of targeted knock-in

**DOI:** 10.64898/2026.10.25.761665

**Authors:** Michael Launspach, Lena O. Raab, Viktor Glaser, Ana-Maria Nitulescu, Stefanie Beez, Maik Stein, Laura M. Hartmann, Shoaib Afzal, Julia Macos, Cassidy J. Quilalang, Joy J. Chen, David N. Nguyen, Ross C. Wilson, Hedwig E. Deubzer, Dimitrios L. Wagner, Annette Künkele

## Abstract

Accurate characterization of CRISPR knock-in requires distinguishing site-specific integration from total donor abundance and donor-derived background while measuring editing at the target site. Here, we present CRISPR-dPCR, a modular multiplex digital PCR framework combining a locked nucleic acid-enhanced cut-site assay with junction-based detection and two orthogonal short-amplicon strategies: Poisson linkage and homology-arm subtraction. We evaluated assay modules across genomic loci, neuroblastoma models and primary human T cells, with selected assays compared on two dPCR platforms. In genotype-defined clones and mixtures with low residual donor, homology-arm subtraction most closely recovered expected knock-in abundance. Linkage provided highly specific evidence of target-cargo association without junction-spanning amplification, whereas junction assays supplied boundary-specific confirmation but were more sensitive to amplicon length and multiplexing. Five-channel panels integrated cut-site editing, donor abundance, copy-number normalization and site-specific knock-in in a single reaction. Residual donor DNA limited quantitative interpretation, particularly in early post-editing samples. Modular extensions quantified defined unintended outcomes, including chromosomal translocations. CRISPR-dPCR integrates complementary measurements into a scalable analytical framework for genome-editing research, cell and gene therapy product development and applications in longitudinal clinical monitoring of targeted integration and transgene persistence.

**Key findings:**

- An LNA-enhanced probe drop-off assay enables cross-platform quantification of CRISPR-Cas9-induced indel formation with improved positive-negative partition separation.
- Systematic benchmarking of three orthogonal knock-in strategies reveals complementary performance: homology-arm subtraction provides the closest quantitative estimates in samples with low residual donor, Poisson linkage provides highly specific evidence of target-cargo association, and junction assays provide direct boundary-specific confirmation but are more sensitive to amplicon architecture.
- Five-channel CRISPR-dPCR integrates cutting, donor abundance, copy-number normalization and site-specific knock-in in a single reaction; linkage and homology-arm measurements remain compatible with multiplexing, whereas long-range junction assays are more susceptible to multiplex-associated artifacts.
- Residual donor DNA is a critical pre-analytical determinant of knock-in quantification, and modular dPCR assays extend the framework to defined unintended outcomes, including donor-derived integration products and chromosomal translocations.

## Introduction

CRISPR-Cas genome editing has transformed functional genomics, biotechnology, and therapeutic engineering, but its safe and effective application depends critically on accurate quantification of editing outcomes. This urgency is underscored by rapid clinical translation, with the first CRISPR-based therapies approved for sickle cell disease and transfusion-dependent β-thalassemia^1,2^, in vivo CRISPR therapeutics advancing into late-stage clinical development^3^, and CRISPR-engineered T cell therapies progressing through early oncology trials^4,5^ . As the field expands beyond gene disruption toward precise genome engineering, HDR-mediated knock-in is becoming an increasingly important strategy for gene and cell therapies^6–8^. Recent studies have demonstrated efficient in vivo site-specific knock-in following systemic delivery of CRISPR components ^9^, highlighting the translational potential of targeted sequence insertion while creating new analytical challenges for characterization of precise genomic integration of multi-kilobase therapeutic cargos.

Current approaches for CRISPR editing analysis include mismatch-cleavage assays, Sanger sequencing deconvolution ^10,11^, capillary electrophoresis ^12^, and next-generation sequencing (NGS) ^13–15^. NGS-based profiling has been highly effective for characterizing unintended editing outcomes, with approaches such as Repair-seq revealing large deletions, rearrangements, and complex repair signatures after CRISPR editing^16,17^. Genome-wide assays including GUIDE-seq, HTGTS, and long-read sequencing further demonstrate that off-target knock-ins, donor capture, and chromosomal translocations can arise at appreciable frequencies^18–20^. However, these methods are not optimized for rapid quantitative assessment of HDR-mediated knock-in. Short-read amplicon sequencing provides nucleotide-resolution analysis but may incompletely resolve the full structure of large or complex integration events involving multi-kilobase therapeutic cargos, whereas genome-wide approaches prioritize comprehensive discovery over standardized quantitative assessment of predefined editing outcomes. Comprehensive characterization of HDR-mediated knock-in therefore requires complementary orthogonal measurements of editing efficiency, precise site-specific integration, total donor abundance, unintended donor integration, and copy-number normalization.

Digital PCR (dPCR) provides absolute nucleic acid quantification with high sensitivity, precision, and scalability, making it well suited for targeted analysis of genome editing outcomes. Previous studies have demonstrated accurate dPCR-based quantification of editing efficiency and copy-number variation^21^, while multiplex hydrolysis-probe systems such as qEva-CRISPR improved detection of edited alleles^22^. Probe drop-off assays further enabled targeted detection of nuclease activity with minimal background^23^. Despite these advances, no unified dPCR framework simultaneously quantifies nuclease activity, precise HDR-mediated knock-in, donor persistence, unintended donor integration, and copy-number variation across diverse genomic loci and editing platforms. Junction-based dPCR assays can detect HDR and NHEJ^24,25^, but require extensive locus-specific optimization, are constrained by amplicon architecture, and are difficult to generalize across editing strategies or multiplex assays. New multiplex dPCR systems such as MEGA dPCR further illustrate the potential of dPCR to quantify multiple editing classes in unified workflows^26^. As CRISPR therapies advance toward mainstream clinical use -including the first FDA-and EMA-approved treatment (exa-cel) and ongoing trials in oncology and hereditary diseases^1,27,28^ -robust, scalable assays for quantifying editing outcomes are increasingly essential.

Here, we introduce CRISPR-dPCR, a unified multiplex framework for comprehensive quantification of CRISPR knock-in outcomes. The framework integrates an LNA-enhanced probe drop-off assay for Cas9 cutting with junction-based dPCR and two orthogonal knock-in assays -a Poisson linkage assay that infers site-specific integration from co-partitioning of donor and target sequences, and a homology-arm subtraction assay -to independently quantify precise genomic integration and total donor abundance. These components are integrated into a five-channel multiplex architecture that simultaneously reports CRISPR cutting, CNV-normalized knock-in, and total donor copy number within a single reaction. The modular framework is readily extensible to additional editing outcomes, including structural variants such as chromosomal translocations. Together, CRISPR-dPCR provides a platform-independent, scalable analytical framework for quantitative characterization of CRISPR editing outcomes across basic research, process development, manufacturing quality control, and translational studies supporting the development of ex vivo and in vivo genome-editing therapies.

## Methods

### Ethics statement

All experiments involving human material were performed in accordance with the Declaration of Helsinki and institutional guidelines. Peripheral blood collection and use of primary human T cells were approved by the Charité ethics committee (EA4/091/19), and written informed consent was obtained from all donors. No animal experiments were performed in this study. Originally unmodified Neuroblastoma cell lines SK-N-BE(2)c and SK-N-AS cells were obtained from Prof. Michael Claus V Jensen at Seattle Children’s Hospital.

### Data Sources and Sample Information

Neuroblastoma knock-in and cut-only cell lines (SK-N-AS and SK-N-BE2c) used in this study were not newly generated; they were produced in the course of our previous work (Launspach et al., 2025^8^). In that work, all genome editing procedures -including gRNA design, HDR template construction, Cas9 RNP electroporation, bulk enrichment or clonal isolation, and full genotypic validation by junction PCR and Sanger sequencing -were performed and described in detail. Previously validated and authenticated cell lines were used as input material for the digital PCR assays described here. Comprehensive methodological information on the generation and validation of these edited neuroblastoma lines is available in Launspach et al., 2025^8^. For this study, peripheral blood was collected from two healthy adult donors under written informed consent, in accordance with the Declaration of Helsinki and with approval from the Charité ethics committee (EA4/091/19). T cells were isolated from fresh leukocyte cones, activated, and edited using our non-viral CRISPR/Cas workflows to generate *TRAC* knock-out (*TRAC*-KO) and *TRAC*-CAR knock-in products. These primary T-cell experiments, including isolation, activation, editing, and downstream dPCR analyses, were performed specifically for the present study. In addition to newly generated samples, we also re-analysed a previously published digital PCR translocation dataset from Glaser et al. (Genome Biology 2023)^29^. That dataset was generated from primary human T cells obtained from the same donor pool under the same ethical approval (EA4/091/19) and edited with multiplex CRISPR gRNAs as described in the original study. No new genome editing or sample generation was performed for the translocation dataset; only the raw ddPCR outputs were reprocessed and analysed here. All human samples -both newly collected and previously published -were handled in compliance with institutional and ethical guidelines.

### T cell editing

Primary human T cells were edited using the Lonza 4D-Nucleofector system. For each reaction, Cas9 ribonucleoprotein (RNP) complexes were assembled immediately before electroporation by mixing synthetic gRNAs (Synthego; sequences in Supplementary Table 3) with Alt-R S.p. Cas9 Nuclease V3 (IDT) at a 2:1 molar ratio (80 pmol gRNA and 40 pmol Cas9 per 1×10⁶ cells) together with 50 μg poly-L-glutamic acid (PGA), followed by a 15-min incubation at room temperature. HDR repair templates (HDRTs) were added at 1.0 μg per 1×10⁶ T cells, and the final RNP-HDRT mixture was combined with 20 μL of T cells resuspended in buffer P3 with Supplement 1. Electroporation was performed in 16-well Nucleocuvette strips using program EH-115, followed by immediate transfer into antibiotic-free RPMI supplemented with 10% FBS. Where indicated, HDR enhancer treatment consisted of AZD7648 (0.375 μM) and ART558 (0.75 μM), added immediately after nucleofection; enhancer-free controls were processed identically without supplementation. After 24 h, medium was replaced to reduce enhancer exposure. For the positive-control dataset re-analysed in Fig. 4f, primary T cells had been electroporated with Cas9 nuclease and three independent gRNAs in the original work, generating six pairwise balanced translocation products. Negative controls consisted of cells electroporated but unedited T cells, providing baseline measurements of Cas9-independent genomic instability. Genomic DNA was isolated 4-6 days after electroporation.

### Genomic DNA Sample Isolation and quality control

Genomic DNA was extracted from harvested cultured cells using the QIAamp DNA Mini Kit (QIAGEN) according to the manufacturer’s protocol. Briefly, up to 5 × 10⁶ cells were lysed with Proteinase K and Buffer AL at 56 °C, followed by ethanol-mediated binding of DNA to a QIAamp Mini Spin Column. Columns were washed with AW1 and AW2 buffers to remove contaminants, and DNA was eluted in Buffer AE and stored at −20°C until use. DNA concentration and purity were measured using a NanoDrop 2000 spectrophotometer (Thermo Fisher Scientific), and samples were accepted only if the A260/280 ratio was 1.8-2.0, the A260/230 ratio exceeded 1.8, and the concentration was ≥10 ng/µL. Samples showing evidence of degradation, discoloration, or precipitation were excluded. For digital PCR, DNA input was required to fall within the validated dynamic range of the assays (typically 5-75 ng per well). Wells were discarded if they exhibited >99.5% positive partitions (saturation) or <1000 total positive partitions across all channels (insufficient template). Reactions affected by pipetting errors, air bubbles, or abnormal partition counts were repeated. Only genomic DNA preparations meeting all QC criteria were used for downstream dPCR analyses.

### Primer and Probe design and optimization

Primer and probe design for all digital PCR assays was performed using Primer3Plus, IDT PrimerQuest, and, where applicable, manufacturer-supported design services (TIB MolBiol), and the final oligonucleotide sequences are listed in **Supplementary Table 1**. Amplicons were designed to be 60-200 bp in length (except long-range Out/In-In/Out assays) with a GC content of 40-60%, while avoiding regions predicted to form stable secondary structures. Target accessibility was assessed using mFold (UNAfold) under assay-relevant conditions (60 °C; 50 mM Na⁺; 2 mM Mg²⁺). Primers were designed to a length of 17-25 nt, with a GC content of 50-60% and a melting temperature (Tm) of 55-65 °C calculated using SantaLucia (1998) parameters at 50 mM monovalent salt and 300 nM oligonucleotide concentration^32^. Runs of more than three consecutive G or C bases were avoided, G/C residues were placed at the 5′ and 3′ ends when possible, and forward/reverse primers were screened to eliminate 3′ complementarity. Primer3Plus and IDT primer quest tools were configured with a detection task, an adjusted divalent cation concentration (3.8 mM) and dNTP concentration (0.8 mM), organism-specific mispriming libraries, and an internal oligo window of 15-30 nt (Tm 64-70 °C). Primer specificity was verified using NCBI Primer-BLAST. Hydrolysis probes were designed to a length of 17-25 nt (preferably <30 nt), with a GC content of 30-80% and a Tm 1-3 °C higher than the corresponding primers. Probes were positioned adjacent to, but not overlapping with, primer binding sites, were enriched for C rather than G residues, and were designed without a 5′-terminal G to avoid quenching after hydrolysis. Probes were synthesized with fluorophores and a quencher as listed in supplementary table 1. For CNV assays, the target and reference gene (*AFF3, NAGK,* and *RPP30*) amplicons were designed using identical PrimerQuest parameters to ensure comparable amplification efficiencies and matched amplicon lengths where possible. For all *AAVS1*-targeting assays (cut-site, transgene, target-locus, linkage, homology-arm, and Out/In-In/Out) and for the *TRAC*-targeting linkage and homology-arm assays run on the QIAcuity platform, we adopted a minimal primer-probe screening strategy. For each assay, we designed two forward and two reverse primers (four possible combinations) together with one probe and evaluated all combinations in a single experimental run. The primer pair showing the clearest cluster separation and most stable amplification behavior was selected. No further optimization of primer composition, annealing temperature, polymerase cycling parameters, or probe conditions was performed.

### Digital PCR for Copy Number Variation (CNV) Analysis

Digital PCR (dPCR) was performed using both the QIAcuity One system (QIAGEN) and the QX200 and QX600 Droplet Digital PCR Systems (Bio-Rad) to quantify absolute DNA copy numbers and determine CRISPR-mediated knock-in efficiencies in transgenic cell lines. For QIAcuity-based assays, reactions were prepared using the QIAcuity Probe PCR Master Mix (QIAGEN) with final concentrations of 800 nM forward and reverse primers and 400 nM hydrolysis probe for all transgene, Out/In-In/Out, linkage and homology-arm assays, and 400 nM forward and reverse primers and 200 nM hydrolysis probe for the reference gene assays. DNA input per reaction was 40 ng for QIAcuity 26k and QX200/QX600 assays, and 12 ng for QIAcuity 8.5k assays (6.6 ng genomic DNA corresponds to approximately 1,000 diploid human cells). To improve template accessibility, 0.05 U/µL XbaI or HindIII (New England Biolabs) were added to each reaction. Reaction volumes were adjusted according to format: 40 µL for 24-well nanoplates, 12 µL for 96-well nanoplates and 22 µl for droplet generation. For QX200/QX600 assays, reactions were prepared using the ddPCR Supermix for Probes (No dUTP) (Bio-Rad) with equal primer-probe concentrations for reference assays and provider-recommended standard primer-probe concentrations for all other assays (900 nM forward and reverse primers and 250 nM probes). Restriction enzyme digestion with XbaI or HindIII (0.05 U/µL) was performed prior to droplet generation. Droplets were generated using the QX200 Droplet Generator or QX600 AutoDG Droplet generator, and samples were transferred to a 96-well PCR plate for thermal cycling. Negative controls containing unmodified genomic DNA were included on both platforms to ensure assay specificity. Thermal cycling for QIAcuity assays was performed on the QIAcuity One system using either a short or long cycling protocol depending on assay type. Both protocols began with enzyme activation at 95°C for 2 min. Short program: 40 cycles of 95°C for 15 s and 58°C for 30 s, followed by on-plate imaging. Long program used for Out/In-In/Out assays: Identical cycling conditions with an additional 2-min extension at 72°C after each annealing/extension step and up to 60 total cycles. Reactions run on the QX200/QX600 were thermocycled on a C1000 Touch Thermal Cycler (Bio-Rad) using the manufacturer-recommended protocol: 95°C for 10 min, followed by 40 cycles of 94°C for 30 s and 60°C for 1 min (ramp rate 2°C/s), and a final step of 98°C for 10 min for droplet stabilization. For QIAcuity assays, fluorescence signals in each partition were quantified using QIAcuity Suite Software (QIAGEN), which calculates absolute copy numbers based on Poisson statistics. For QX200/600 assays, droplets were read using the QX200/600 Droplet Reader, and data were processed with QuantaSoft Analysis Pro (Bio-Rad). Fluorescence thresholds were manually set for each channel per plate by placing the cutoff immediately above the upper boundary of the negative partition cloud, based on the collective negative droplet distribution across all wells on that plate -with each plate including at least one negative control (unedited genomic DNA or no-template control) to anchor the negative cloud -, and this was performed blinded to sample identity. Wells exhibiting indistinct negative clouds, excessive rain, or abnormal amplitude distributions were excluded and rerun to ensure consistent cluster separation. Final copy-number estimates were calculated from thresholded partition counts using standard Poisson occupancy modelling. Transgene copy number was normalized to endogenous reference genes *AFF3, NAGK* or *RPP30* after empirically determining locus-specific copy-number relationships where required. This enabled estimation of target copies per 100 cells while accounting for variability in genomic DNA input. Partition/droplet plots, scatter plots, and quantitative tables were exported directly from the respective software platforms.

### Homology-arm-based knock-in quantification

Knock-in abundance was quantified from three digital PCR channels-transgene (T), homology-arm(H), and reference gene (R)-after converting positive droplet counts into absolute copies per µL via Poisson inversion (copies_per_uL = −ln(1 − p) / V, where p is the fraction of positive partitions and V is the partition volume). The HA method estimates on-target HDR by subtracting excess homology-arm signal arising from persisting episomal donor or off-target integration. The knock-in copy number (CN) per 100 cells was calculated as:

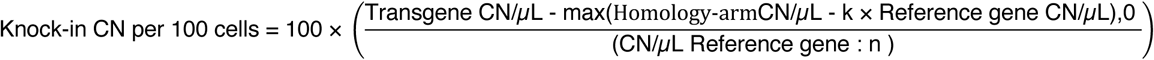

where k is the target-locus-to-reference-gene copy-number ratio in unmodified cells (typically ∼1 for diploid loci); and n is the reference-gene copy number per cell (typically 2 for diploid loci). This calculation assumes equivalent amplification efficiencies for genomic and donor-derived homology-arm sequences and stable reference-gene ploidy. A full derivation and discussion of assumptions, edge cases, and failure modes (e.g., truncated donor fragments) are provided in **Supplementary Note 5**.

### Linkage-based knock-in quantification

The linkage assay uses a three-channel dPCR architecture in which wild-type (WT-only) molecules are detected in the ROX channel, transgene-only molecules in the FAM channel, and true HDR-integrated WT-TG molecules generate simultaneous ROX⁺FAM⁺ signals. Positive droplet counts were converted to absolute copies per µL using Poisson inversion. Knock-in inference proceeded in two steps: (i) hypothesis testing for non-random co-localization and (ii) maximum-likelihood estimation of the fraction of linked molecules. Under the null hypothesis of no linkage (H₀), WT and TG molecules distribute independently across partitions, and the expected double-positive frequency is determined by the marginal Poisson occupancies of the two channels. Statistical significance was assessed using a binomial test comparing the observed double-positive count to the expectation under H₀. When H₀ was rejected, we estimated the mixture proportions of WT-only, TG-only, and linked WT-TG molecules -denoted a, b, and c, respectively -using a maximum-likelihood Poisson mixture model. Here, a represents the fraction of ROX-only molecules, b the fraction of FAM-only molecules, and c the fraction of WT-TG linked molecules arising from true HDR integration. The model jointly fits the observed ROX⁺, FAM⁺, and ROX⁺FAM⁺ frequencies to recover the mixture parameters under a shared Poisson occupancy rate. The fraction of target copies attributable to linked WT–TG molecules was calculated as *c / (a+b+2c)*. Linked copies per µL were then calculated as: *linked_copies_per_uL = linked_target_copy_fraction × (FAM_total_copies + ROX_total_copies)*. Knock-in frequency per 100 cells was then computed as: *KI_per_100_cells = 100 × (linked_copies_per_uL / cell_equivalents_per_uL)*, with cell equivalents per µL derived from reference-gene copies per µL. Full mathematical derivation, likelihood equations, and boundary conditions are provided in **Supplementary Note 6.**

### Determination of Assay performance parameters

Assay performance parameters were determined in accordance with established digital PCR evaluation guidelines, and all reporting follows the digital MIQE (Minimum Information for Publication of Quantitative Digital PCR Experiments) standards^30^. Formulas for the quantitative metrics used in this study are provided in Supplementary Table 2. “Rain” was defined as partitions exhibiting intermediate fluorescence intensities that were not clearly classifiable as positive or negative; manual threshold placement was used to assign these partitions, and copy numbers were calculated both including and excluding rain, with the relative difference reported as percent rain. For duplex Out/In-In/Out assays, rain was quantified only for the transgene channel because the *AFF3* reference assay produced negligible rain, whereas for linkage and homology-arm assays, rain was calculated separately for each component assay (wild-type and transgene, or homology-armand transgene, respectively), and the percentages were combined to obtain cumulative rain. Rain was assessed for each assay as the mean rain percentage across three independent runs. Diagnostic sensitivity and specificity were evaluated by classifying all results as true positive, true negative, false positive, or false negative using the median limit of blank (LoB) of each assay type as the cutoff. Quantitative accuracy was assessed using a genomic DNA dilution series (50 ng, 5 ng, 0.5 ng, 0.05 ng), in which measured copy numbers at each dilution were normalized to the undiluted sample to yield relative accuracy (1.0 = 100% accuracy). Quantitative recovery was additionally assessed by comparing measured knock-in copy numbers with expected values in genotypically confirmed single-cell clones and defined 1:1 mixtures of edited single-cell clones and unedited cells, corresponding to expected abundances of 50 or 100 knock-in copies per 100 cells. The LoB was determined from multiple independent negative controls by taking the 95th percentile of measured copy numbers, corresponding to a theoretical specificity of 95% and providing the threshold for background discrimination as recommended by CLSI EP17-A2^31^. Precision was evaluated using the conventional coefficient of variance (CoV), calculated both within runs (intra-assay CoV, using replicate measurements of the same sample within a single run) and across runs (inter-assay CoV, using repeated measurements of the same sample across multiple runs), with all assay-sample CoVs aggregated per assay to describe overall dispersion. The probability of detection was determined from a low-input dilution series (5 ng, 2.5 ng, 1.25 ng, 0.625 ng, 0.3125 ng DNA). Assay efficiency was evaluated using measurements of samples tested alone and in combination with a synthetic gene block containing target sequences for all assays and an orthogonal reference assay (SARS-CoV Nucleocapsid gene targeting assay^33^, ATT550). For gene block-only measurements, the ratio of target assay to reference assay was expected to be 1:1 and deviations were used to quantify assay efficiency. In combined gene block + genomic DNA measurements, the gene block contribution was subtracted from the total signal and normalized to the sample-only measurement to assess recovery efficiency. For homology-arm assays, in which channel overlap prevented inclusion of the SARS-CoV reference assay, previously measured efficiencies of the corresponding transgene assays in the gene block were used to infer 100% efficiency, and transgene copy numbers from the gene block served as the endogenous control.

### Automated computational simulations of dPCR Assays

Monte Carlo simulations were implemented in R (v4.1) using foreach, doParallel, and dplyr to benchmark the HA and linkage assays across experimentally relevant ranges of cell input, donor background, knock-in fraction, and partition count. Full scripts and a Shiny-based calculator for the linkage assay will be available via Zenodo (**DOI: 10.5281/zenodo.21432198**).

HA simulations: For each parameter combination, reference, homology-arm, and transgene molecule numbers were computed from the specified number of cells, ground-truth knock-in fraction, and episomal donor molecules. Molecule occupancy was simulated using Poisson loading, droplet positivity was converted to copies/µL by Poisson inversion, and knock-in frequency per 100 cells was calculated using the closed-form HA equation (see Methods “*Homology-arm-based knock-in quantification”* and **Supplementary Note 5**). Classification metrics (TP, FP, FN, TN), sensitivity, and specificity were computed relative to the ground-truth KI input. Simulation grids varied cell input (500-32,000 cells), donor noise (0-128,000 molecules), partition number (24,000-48,000) and knock-in frequency (0-100%). For each condition, 100 independent replicates were generated, and both replicate-level outputs and aggregated summaries were exported for downstream analysis.

Linkage simulations. WT-only, TG-only, and linked molecules were distributed into partitions by random assignment. Positive counts were converted into single-and double-positive fractions, used for null-hypothesis testing and mixture-fraction estimation as described in Methods “*Linkage-based knock-in quantification.”* and **Supplementary Note 6**. Partition counts, knock-in input, and FAM copy number were varied across wide ranges (8,000-60,000 partitions; 1,000-10,000 cells; low to very high TG loads; 0-100% knock-in). For each condition, 10-100 replicates were performed, and we recorded mean knock-in estimates, 95% CI widths, expected vs observed double positives, and empirical H0 rejection rates. Additional simulations examined limit-of-blank behaviour and statistical power under rare-event scenarios.

### Multiplex assay design and fluorophore selection

Multiplex panel composition and dye assignment were guided by the optical layout of the QIAcuity and QX200/600 systems as well as manufacturer recommendations. Fluorophores were distributed across channels to minimize spectral overlap, with FAM, HEX, ATTO550, ROX, and Cy5 reserved for the highest-priority assays (cut-site, transgene, reference, and target-locus/linkage/HA assays), whereas ATTO590 was used only when adequate separation from adjacent channels had been confirmed in simplex runs. Assays expected to produce high-amplitude signals (such as reference or target-locus assays) were preferentially placed in channels with minimal bleed-through into lower-amplitude channels. Fluorophore combinations were chosen to avoid assigning probes with overlapping emission spectra to neighbouring channels within the same reaction, and all candidate dye sets were validated in simplex mode prior to multiplexing. Primer and probe concentrations in multiplex reactions were identical to those used in the corresponding simplex assays, and no further concentration optimization was performed. Template DNA input (ng per well) and thermocycling conditions were identical between simplex and multiplex reactions. All assays were run using the same QIAcuity cycling program as described above, i.e. either the short protocol or the long protocol when an Out/In-In/Out assay was included. This ensured direct comparability of assay performance across simplex and multiplex formats.

### Statistics and Reproducibility

All statistical analyses were performed in accordance with the Digital MIQE guidelines and the CLSI EP17-A2 framework for analytical detection capability. Formulas for quantitative metrics-rain, accuracy, diagnostic sensitivity/specificity, LoB, CoV, assay efficiency, and probability of detection-are provided in **Supplementary Table 2** and were not redefined in the Statistics section to avoid redundancy. Unless stated otherwise, data are shown as mean ± SD. No formal sample-size calculation was performed, as the study focused on analytical benchmarking of assay architectures; replicate numbers followed established standards for dPCR assay development and plate-format constraints. The experiments were not randomized. Biological replicates were defined as independent DNA preparations or independently edited cell batches. Technical replicates were averaged within each biological replicate and were not used as independent observations for hypothesis testing. Unless otherwise specified in the figure legends, comparisons across classes (Out/In-In/Out, linkage, homology-arm) were performed using Kruskal-Wallis tests with Dunn’s post-hoc correction. Pairwise comparisons (e.g., unsorted versus sorted samples, internal versus external reference controls) were analysed using two-sided Mann-Whitney tests. Two-way ANOVA with Tukey’s correction was used for comparisons involving multiple factors such as donor × assay type, simplex versus multiplex dPCR, and dilution-or cycle-number-dependent effects. Correlation analyses-including comparisons between dPCR-derived KI frequencies and flow-cytometry-derived reporter positivity-used Spearman’s rank correlation. Linear regression was applied for concordance analyses (e.g., clone-level true vs measured copy numbers). For linkage-assay inference, binomial tests were used as a mandatory pre-test: only samples rejecting the null hypothesis of random WT-TG co-localization (p < 0.05) proceeded to maximum-likelihood estimation of linked fractions. Statistical analysis. To compare the performance of the four methods (Out/In, LA, HA, FC) across multiple target samples in Figure 2d, we applied a linear mixed-effects model. In this model, Method was treated as a fixed effect, while Sample was included as a random effect to account for repeated measurements within the same biological sample. This approach corresponds to a repeated-measures design but is more flexible, as it allows for unbalanced data with missing values. The model was fitted in R (version 4.3.1) using the lmerTest package (function lmer). Significance of the main effect of Method was assessed via ANOVA, and post-hoc pairwise comparisons between methods were performed using the emmeans package with Tukey’s adjustment for multiple testing. Monte-Carlo simulations used the same statistical decision rules as described above (LoB thresholds, binomial pre-tests, diagnostic specificity criteria) and report aggregate sensitivity, specificity, and accuracy metrics across 10-100 simulation replicates per condition. All analyses were two-sided, performed in GraphPad Prism v10 or R v4.x, and exact statistical tests and p-values are provided in the figure legends and the Source Data file.

Additional methods are described in the supplementary methods section. Equipment, consumables, antibodies and software used are listed in **Supplementary tables 4-6**.

## Results

### Quantification of CRISPR Cutting Using a Probe Drop-Off dPCR Assay

To establish a high-precision cutting module for CRISPR-dPCR, we benchmarked a probe drop-off digital PCR assay in which a hydrolysis probe spanning the CRISPR cut site no longer binds when editing and indel formation disrupts the target sequence, resulting in a measurable loss of fluorescence relative to a reference copy number assay (**Fig. 1a**). Cutting frequency was calculated from the proportional decrease in unedited target copies, normalized to an internal reference assay and the corresponding untreated control, enabling direct estimation of the fraction of edited genomes (**Fig. 1a**). To improve signal discrimination, we compared a conventional DNA hydrolysis probe with a locked nucleic acid (LNA)-modified probe containing a methylene bridge that constrains the ribose in the C3′-endo conformation, rather than the predominantly C2′-endo conformation of unmodified DNA (**Fig. 1b**). Across two neuroblastoma models (SK-N-AS and SK-N-BE2c), electroporated with *AAVS1* targeted gRNA/Cas9 ribonucleoprotein (RNP), the LNA probe consistently yielded significantly higher estimated cutting frequencies than the standard probe (mean difference SK-N-AS: Δ = 17 percentage points, 95% CI [2.2-31.5]; SK-N-BE2c: Δ = 41 percentage points, 95% CI [7.2-75.9]; two-way ANOVA with Tukey’s correction, both p < 0.005) and produced values closely matching Synthego ICE analysis in both cut-only and HDR co-delivery conditions (**Fig. 1c**, **Supplementary Fig. 2a).** Amplitude plots showed substantially less fluorescence rain with the LNA probe, yielding sharper positive-negative partition separation **(Supplementary Fig. 1a).** To characterize analytical performance, we constructed a synthetic gene block containing the cut-site assay target, multiple knock-in assay targets, and a unique internal control sequence derived from SARS-CoV-2 (**Fig. 1d**; **Supplementary Fig. 2b).** Using this defined template, we observed near-quantitative amplification efficiency (∼100%) across all assay configurations (**Fig. 1e**). To determine whether unedited target sequences could also be quantified in a predominantly edited genomic background, the unedited target sequence-containing gene block was spiked into edited tumor-cell genomic DNA. The LNA probe recovered a mean of 90% of the expected unedited target copies across the genomic DNA backgrounds (**Fig. 1f**), supporting its ability to quantify residual unedited alleles within a complex edited sample. We next validated performance in primary human T cells undergoing *TRAC* editing. Flow cytometry confirmed efficient *TRAC* knockout (>95% CD3 loss) across two donors following RNP electroporation, irrespective of donor template co-delivery or HDR enhancer treatment (**Fig. 1g**; **Supplementary Fig. 1b).** The cut-site assay detected near-complete *TRAC* cutting on both the QIAcuity and QX200 platforms, demonstrating robust performance across two dPCR systems (**Fig. 1h**). Together, these data establish the LNA-enhanced probe drop-off assay as a precise, efficient, and platform-agnostic module for quantitative measurement of CRISPR-Cas9 cutting within the CRISPR-dPCR framework.

**Fig. 1.**
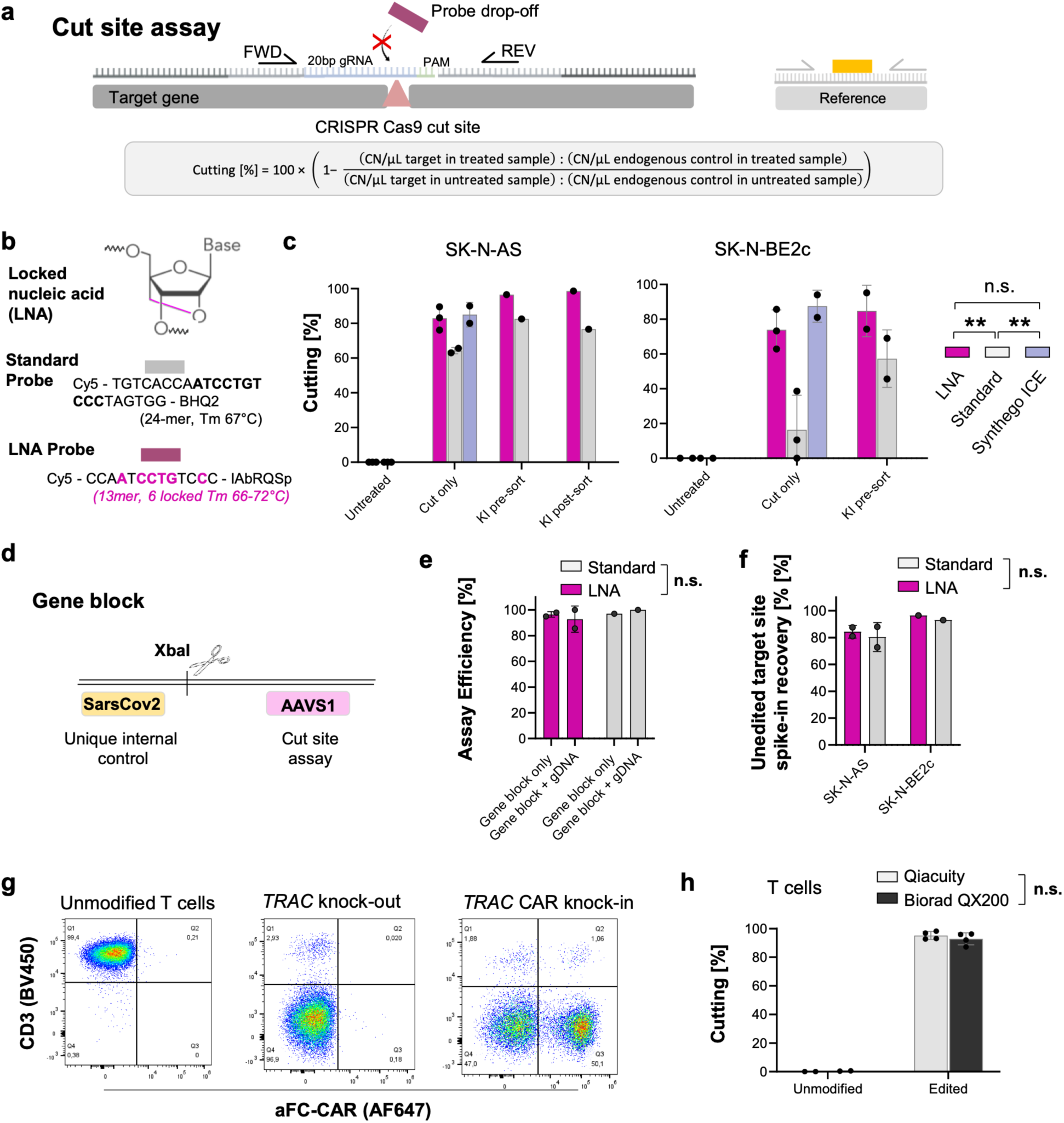
A probe drop-off digital PCR assay enables high-precision quantification of CRISPR-Cas9 cutting across cell types and platforms. (**a**) Schematic of the cut-site assay in which a hydrolysis probe spanning the Cas9 cleavage site is lost upon double-strand break formation, enabling quantification of cutting by comparing target copy numbers to an internal reference assay. (**b**) Standard DNA probe versus locked nucleic acid (LNA)-modified probe design for AAVS1-targeting, with locked nucleotides indicated. (**c**) Cutting efficiencies measured in SK-N-AS and SK-N-BE2c cells edited either with Cas9 RNP alone (“cut only”) or together with HDR donor templates (CXCL10 knock-in, pre-and post-FACS enrichment (pre-/ post-sort)). Cutting values obtained with the standard or LNA probe are compared with Synthego ICE estimates. (**d**) Schematic of the synthetic gene block used for analytical characterization, containing the cut-site target region, donor-derived assay sites, and an orthogonal SARS-CoV-2 control sequence separated by an XbaI restriction site. (**e**) Analytical efficiency of standard and LNA probes measured on gene block DNA alone and in unedited genomic DNA spike-ins. (**f**) A gene block containing the unedited target sequence was spiked into the edited genomic DNA samples analysed in panel c. The expected number of unedited target copies was calculated from the residual unedited genomic target copies and the added gene-block copies. Spike-in recovery was calculated as the observed relative to the expected number of unedited target copies in the mixture. **(g)** Representative flow cytometry plots of primary human T cells edited at TRAC, showing CD3 loss (TRAC-KO) and CAR expression (TRAC-CAR). **(h)** Quantification of TRAC cutting in edited primary T cells shown in (g), using the LNA probe on the QIAcuity and QX200/600 platforms, demonstrating cross-platform concordance. **Data presentation:** (c, e-h) Mean ± SD; dots represent biological replicates. **Statistical tests:** (c) Two-way ANOVA with Tukey’s post-hoc correction, (e) Two-way ANOVA with Sidak’s multiple comparisons test, (f) Two-way ANOVA, (h) Two-sided Mann-Whitney test; exact p values are reported in the Source Data file. p values: *<0.05, **<0.01, ***<0.001, ****<0.0001; n.s., not significant. Source data are provided as a Source Data file.

### Site-Specific Knock-In Quantification

Because accurate quantification of HDR-mediated knock-in requires correction for locus-specific copy number variation, we first established CNV baselines for each target locus using HiC and dPCR, revealing substantial discrepancies in *AFF3* and *AAVS1* copy numbers in SK-N-AS and SK-N-BE2c cells that necessitated locus-specific normalization for all downstream CRISPR-dPCR analyses **(Supplementary Fig. 3a-c and Supplementary Note 1).** We next established dPCR assays for non-site-specific transgene quantification. As expected, total transgene measurements included both correctly integrated donor molecules and non-integrated donor DNA, including extrachromosomal donor DNA and off-target integrations **(Supplementary Fig. 4a-d).** To characterize HDR-mediated knock-in, we compared three orthogonal dPCR strategies (**Fig. 2a-c**). Directional Out/In or In/Out assays amplified the genome-donor junction but showed increasing rain and reduced performance with longer amplicons **(Supplementary Figs. 5, 6a,b)**^25^. The triplex linkage assay measured transgene, target locus and endogenous reference simultaneously and inferred integration from double-positive partitions exceeding random Poisson co-partitioning; samples rejecting the null hypothesis of random co-localization were quantified by maximum-likelihood estimation (**Fig. 2b**). Two independent linkage designs gave concordant results with minimal rain **(Supplementary Figs. 7, 8)**. The HA subtraction assay estimated site-specific knock-in by comparing transgene, homology-arm and reference copy numbers: correctly integrated alleles contribute matched transgene and homology-arm copies, whereas residual donor DNA or unintended donor-derived integration products can increase homology-arm counts relative to transgene counts (**Fig. 2c**; **Supplementary Fig. 9).** Across loci and donors, knock-in estimates ranked HA > linkage > Out/In–In/Out (median 68%, 41.8% and 34.4%, respectively, versus 54% by flow cytometry; **Fig. 2d**). In negative controls containing residual extrachromosomal donor DNA or off-target donor integration, HA produced donor-dependent apparent knock-in signal, linkage showed no detectable false-positive signal using binary gating, and rare Out/In false positives increased with donor abundance, consistent with PCR-mediated template switching (**Fig. 2e**; **Supplementary Fig. 6c-e).**

We next systematically compared the analytical performance of the three assay classes. Out/In-In/Out assays generated the highest rain fraction, defined as the proportion of partitions with intermediate fluorescence that could not be clearly classified as positive or negative, whereas linkage and HA assays produced markedly cleaner partitioning (**Fig. 3a**; **Supplementary Figs. 5, 7, 9**). All methods showed high diagnostic sensitivity -the proportion of known knock-in-positive samples correctly identified -across knock-in ranges of approximately 3-6 copies per 100 cells in SK-N-AS and 3-5 copies per 100 cells in SK-N-BE2c. Linkage sensitivity improved with increasing partition number in both the 26k and 8.5k formats (**Fig. 3b**). Because promoter-driven transgene expression does not uniquely establish on-target integration, we benchmarked quantitative performance using genotypically confirmed monoallelic single-cell clones and defined 1:1 mixtures of edited single cell clones and unedited cells, corresponding to expected abundances of 50 or 100 knock-in copies per 100 cells. HA measurements most closely matched these expected values, linkage assays showed moderate underestimation, and Out/In-In/Out assays showed the largest negative bias. Accordingly, quantitative recovery, calculated as measured relative to expected knock-in abundance, was highest for HA (mean 1.07), intermediate for linkage (0.77), and lowest for Out/In-In/Out (0.51) (**Fig. 3c,d**). Limit-of-blank (LoB) analysis, defined as the upper boundary of apparent signal observed in clean negative controls, showed the lowest background for HA assays, followed by Out/In-In/Out assays, whereas linkage assays exhibited a higher apparent LoB because of stochastic WT–transgene co-partitioning inherent to Poisson loading (**Fig. 3e**; **Supplementary Fig. 17a).** Increasing partition counts improved discrimination of true linkage from random co-partitioning, as reflected by the per-well p-value distributions **(Supplementary Fig. 17b).** The linkage statistical pre-test correctly classified 38 of 39 negative controls as negative, corresponding to a specificity of 97.4% (95% CI, 86.6-99.9%) (**Fig. 3f**), while diagnostic specificity across all assay classes remained high (90–100%; **Fig. 3g**). Out/In-In/Out assays also had the highest design-failure rate, reflecting their dependence on long-range PCR and complex primer-probe configurations, whereas linkage and HA assays generally required minimal redesign (**Fig. 3h**). Dilution-series testing showed that linkage remained accurate at 1:10 dilution but declined at 1:100-1:1,000, with performance retained longer at higher target abundance; Out/In-In/Out assays performed similarly or marginally better at the most extreme dilutions (**Fig. 3i**; **Supplementary Fig. 16a)**. Probability-of-detection analysis confirmed these trends (**Fig. 3j**). Cycle-number testing showed robust performance of linkage, transgene and HA assays at ≥40 cycles, whereas junction assays required assay-specific optimization **(Supplementary Fig. 10).** Simulations further showed that linkage sensitivity increased with partition count and that combining at least three wells substantially improved detection while maintaining high specificity **(Supplementary Figs. 11, 12, 14a; Supplementary Note 2).** HA accuracy remained stable across most occupancy levels, with modest degradation near saturation and instability only at extreme saturation (>99.9% positive partitions, **Supplementary Figs. 13, 14b; Supplementary Note 3)**. All assays showed high amplification efficiency using gene-block standards alone and after spike-in into genomic DNA, indicating that the observed quantitative differences were not attributable to basic assay inefficiency **(Supplementary Fig. 16b).** Linkage assays also tended to show lower intra-and inter-assay coefficients of variation than junction assays **(Supplementary Fig. 18a,b).** Although the individual WT, transgene and HA components amplified robustly by qPCR, linkage inference required digital partitioning, whereas HA subtraction was in principle compatible with relative qPCR quantification **(Supplementary Fig. 15a,b).** Overall, HA provided the most accurate estimates in samples with minimal residual donor DNA, linkage provided high analytical specificity, and optimized junction assays supplied independent boundary-specific evidence of integration.

**Fig. 2.**
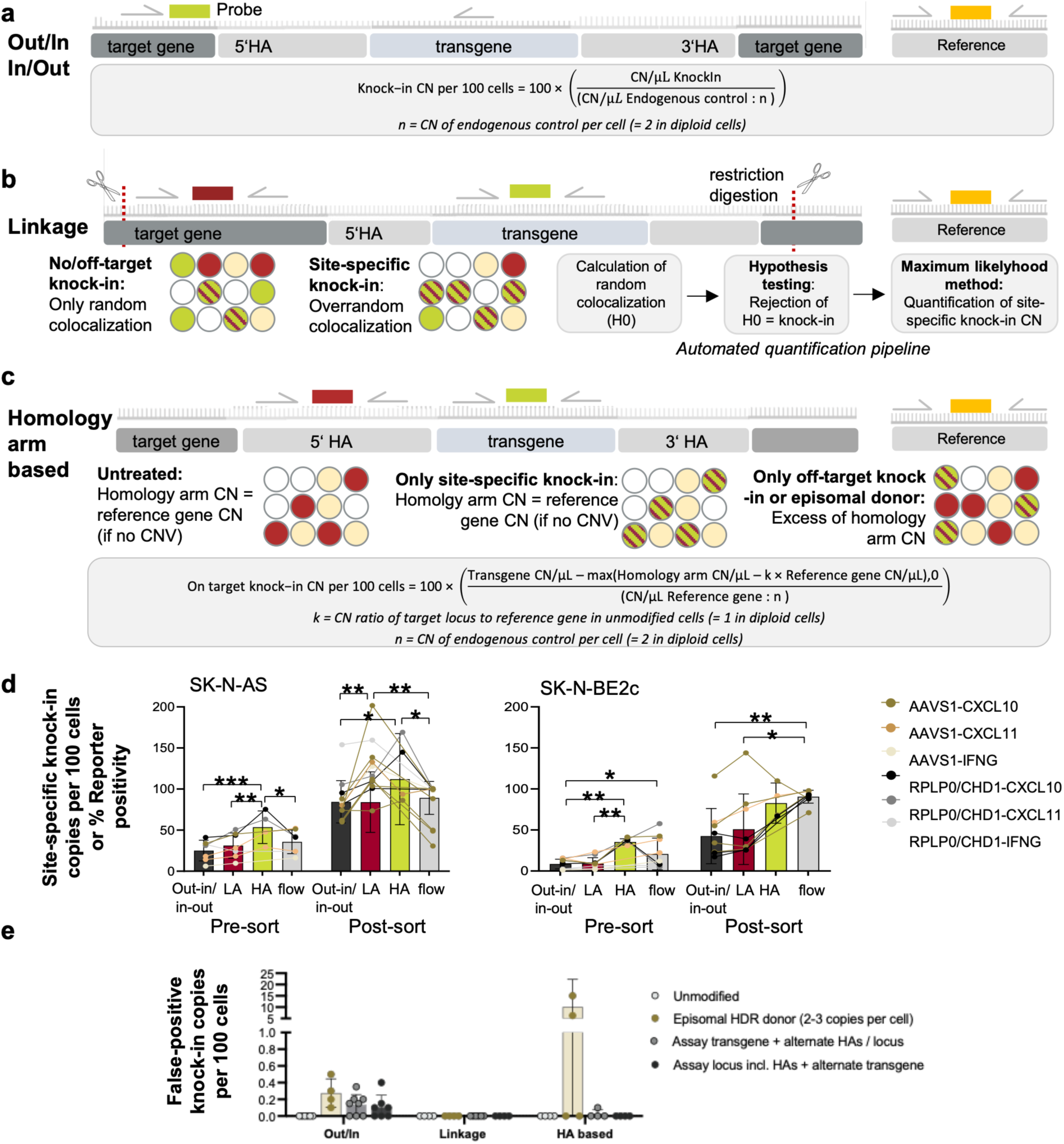
Digital PCR strategies for site-specific knock-in quantification and comparison of assay performance across loci and payloads. (a) Schematic of the long-range Out/In-In/Out junction assay architecture, in which primers span the endogenous locus and HDR payload to generate directional amplicons reporting on-target integration; knock-in copies per 100 cells are calculated by normalizing transgene copies to an endogenous reference gene. (b) Overview of the linkage assay workflow: wild-type (WT) and transgene (TG) molecules are independently labeled and partitioned, true HDR events producing WT⁺TG⁺ co-positive droplets above Poisson-predicted random co-localization; after restriction digestion, observed double-positive rates are tested against the null hypothesis of random co-localization (H₀), and when H₀ is rejected, maximum-likelihood estimation quantifies the fraction of linked WT-TG molecules. (c) Principle of the homology-arm (HA) subtraction assay: untreated or perfect knock-in cells display equal HA and reference-gene copy numbers (after CNV correction), whereas off-target integrations or episomal donor increase HA copies relative to transgene copies; site-specific knock-in is estimated by subtracting excess HA signal from transgene copies and normalizing to the reference gene. **(d)** Comparison of site-specific knock-in values across Out/In-In/Out, linkage, and HA assays in SK-N-AS and SK-N-BE2c cells edited with CXCL10, CXCL11, or IFNG donors; assays were evaluated pre-and post-sort alongside flow-cytometry-derived reporter positivity, with HA assays yielding values closest to reporter-positive fractions and Out/In-In/Out assays showing greater variability. **(e)** Evaluation of assay specificity using unmodified cells, cells containing episomal HDR donor DNA, and cells harboring assay-relevant off-target integrations. Apparent site-specific knock-in copies per 100 cells were measured by the Out/In-In/Out, linkage, and homology-arm (HA) subtraction assays. **Data presentation:** (d, e) mean ± SD; dots represent biological replicates. **Statistical tests:**(d) Mixed-effects model with Tukey post-hoc test; exact p values provided in the Source Data file. p values: *<0.05, **<0.01, ***<0.001; n.s., not significant. Source data are provided as a Source Data file.

**Fig. 3.**
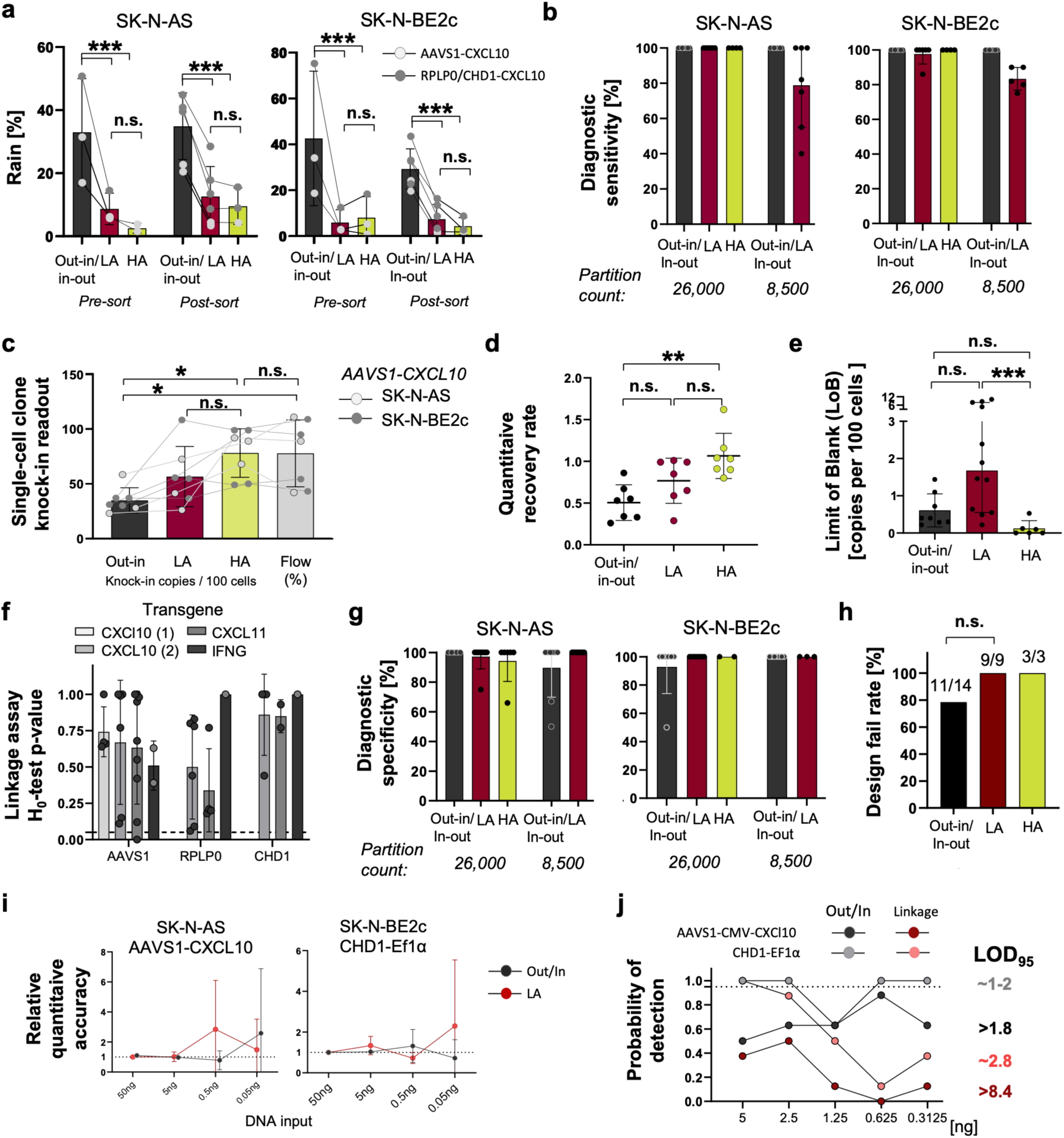
Comparative analytical performance of Out/In-In/Out, linkage, and homology-arm (HA) assays for site-specific knock-in quantification. (a) Rain percentages for Out/In-In/Out, linkage, and HA assays in SK-N-AS and SK-N-BE2c cells across pre-sort and post-sort samples for AAVS1-CXCL10 and RPLP0/CHD1-CXCL10 donors. (b) Diagnostic sensitivity of all assay types in SK-N-AS and SK-N-BE2c at high (26k) and low (8.5k) partition numbers. (c) Site-specific knock-in estimates obtained with Out/In-In/Out, linkage, and HA assays in genotypically confirmed monoallelic single-cell clones and defined 1:1 mixtures of edited and unedited cells, corresponding to expected abundances of 50 or 100 knock-in copies per 100 cells; flow-cytometric reporter positivity is shown for comparison. **(d)** Quantitative recovery of each assay relative to the genotype-defined expected knock-in abundance for the samples shown in (c). **(e)** Limit-of-blank (LoB) analysis for HA, Out/In-In/Out, and linkage assays; median LoB values indicated, with the highest LoB observed for the linkage assay reflecting stochastic WT-TG co-localization in negative controls and mandating preliminary binomial pre-testing before knock-in quantification. **(f)** Linkage assay H₀-test p-values (binomial pre-test) across AAVS1, RPLP0, and CHD1 loci for four donor types, measured using the 26,000-partition format, showing consistent rejection of random co-localization in true knock-in samples. **(g)** Diagnostic specificity of all assay types at high and low partition counts in SK-N-AS and SK-N-BE2c. **(h)** Design-failure rate for each assay type (i.e. primer-probe combination did not produce meaningful results or excessive false-positives in negative samples). **(i)** Quantitative accuracy across a genomic DNA dilution series. **(j)** Probability-of-detection curves across DNA inputs for representative donors and loci with n = 8 replicates measured for each DNA input dilution step per sample. **Data presentation:** (a-d, f-j) mean ± SD, (e) median ± 95% CI: dots represent biological replicates. **Statistical tests:** (a,e,g) Kruskal-Wallis with Dunn’s post-hoc correction; (d) one-way ANOVA with Tukey’s post-hoc test; (h) Fisher’s exact test. (i) two-way ANOVA; exact p values are provided in the Source Data file. p values: n.s., not significant; *<0.05; **<0.01; ***<0.001. Source data are provided as a Source Data file.

### Extension of CRISPR-dPCR to unintended genome-editing outcomes

CRISPR-mediated genome engineering can generate unintended genetic alterations in addition to the desired on-target edits. To quantify these events, we designed digital PCR assays capable of detecting both off-target knock-ins and chromosomal translocations with high analytical depth (**Fig. 4a,b,e).** The homology-arm (HA) assay quantifies both site-specific HDR and the total burden of donor-derived sequences by comparing total transgene, site-specific knock-in and homology-arm copy numbers (**Fig. 4b** **and Supplementary Note 4).** Excess homology-arm signal reflects donor molecules that cannot be explained by correctly integrated HDR alleles, including residual extrachromosomal donor DNA and unintended donor-derived integration products. Accurate interpretation requires locus-specific CNV correction and consideration that extensive donor fragmentation may alter measured HA/transgene ratios. Consequently, HA measurements are most informative when interpreted together with orthogonal assays such as linkage measurements, which assess physical association of donor and target loci, and junction assays, which confirm the expected genome-insert boundaries (**Fig. 4a,b**). In neuroblastoma cells, the HA assay distinguished on-target knock-in from excess donor-derived signal attributable to off-target integration and/or persisting episomal donor in both unsorted and sorted populations (**Fig. 4c**). Bulk sorted cells -enriched for high transgene expression -displayed an increased ratio of off-target/episomal donor signal relative to on-target HDR, consistent with selective enrichment of donor-containing cells irrespective of the genomic context of integration (**Fig. 4d**). Because CRISPR can generate chromosomal translocations through illegitimate joining of independent DSBs, we incorporated dPCR assays and datasets from a published use case to quantify these events (**Fig. 4e**)^29^. In these data, primary T cells had been electroporated with three gRNAs, generating six possible pairwise translocation products. Re-analysis of the data showed that five of the six assays detected statistically significant junction formation relative to controls (adjusted p values 0.0025 to <0.0001), while TRAC-CIITA did not show significant enrichment (p=0.54, **Fig. 4f**). Analytical characterization confirmed high depth and specificity of the translocation assays. Limit-of-blank analysis yielded a mean LoB of 0.05 copies -corresponding to an analytical detection depth of approximately 1:2,000 genomes -with specificity exceeding 80% in all but one assay (**Fig. 4g**). Sensitivity was 100% for all samples, reflecting the combination of low LoB and the substantial translocation load generated by multiplex gRNA editing (**Fig. 4f, g**). Together, these results demonstrate that the CRISPR-dPCR framework can be extended beyond on-target knock-in quantification to defined unintended editing outcomes, including donor-derived integration products and CRISPR-induced chromosomal rearrangements.

### Multiplex integration of core CRISPR-dPCR modules

To increase analytical throughput while minimizing sample consumption, we implemented five-channel CRISPR-dPCR multiplex panels that combine multiple assay modules within a single reaction. We designed multiplex assays for *AAVS1* editing in neuroblastoma cell lines and *TRAC* editing in primary human T cells for direct comparison with flow cytometry measurements (**Fig. 5a**, **Supplementary Fig. 19)**. For *AAVS*1, complementary Out/In-and HA-based multiplex panels simultaneously quantified CRISPR cutting, total transgene copy number, copy-number normalization, and site-specific knock-in. Equivalent panel architectures were developed for *TRAC* editing, incorporating either an internal *TRAC* reference or conventional external reference genes for CNV normalization (**Fig. 5a**). Across *AAVS1*-edited samples, multiplexed linkage and homology-arm assays produced knock-in estimates that closely matched their corresponding duplex assays, whereas multiplexed long-range Out/In assays exhibited non-specific amplification and false-positive signals in both edited samples and unmodified genomic DNA spiked with a synthetic knock-in gene block representing the expected HDR junction (**Fig. 5b,e**). CRISPR cutting measurements remained highly consistent across all multiplex configurations, and application of the QIAcuity cross-talk correction matrix improved fluorescence cluster separation but did not eliminate junction assay-specific artifacts (**Fig. 5b**; **Supplementary Figs. 20-22**).

**Fig. 4.**
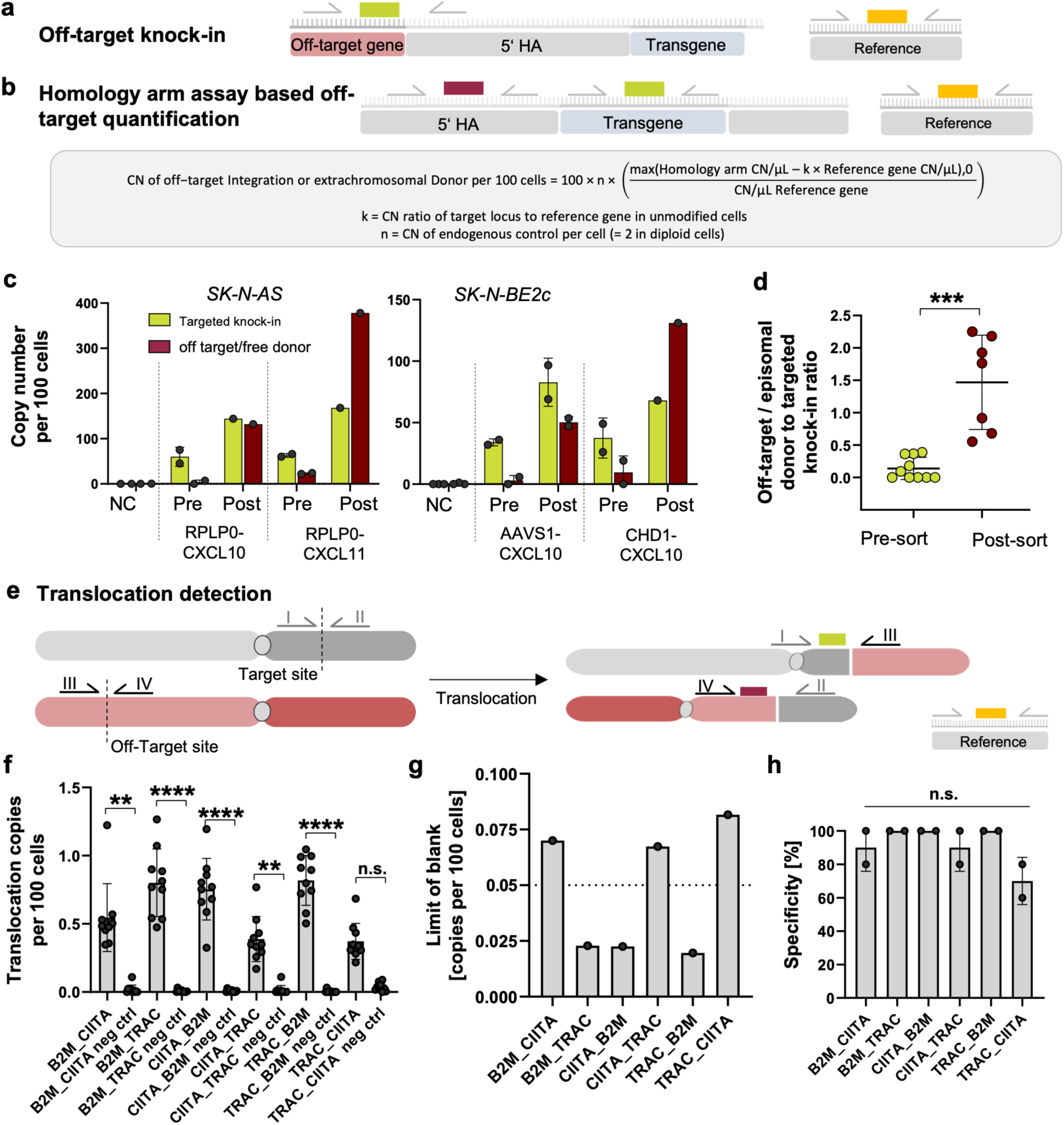
Digital PCR-based quantification of off-target donor integration and CRISPR-induced chromosomal translocations. **(a)** Schematic of a locus-specific off-target knock-in assay in which a forward primer at a known off-target genomic site and a reverse primer in the donor homology-arm amplify unintended HDR integrations, enabling direct detection and quantification of site-specific off-target knock-ins. **(b)** Principle of the homology-arm (HA)-based framework for global off-target donor quantification: the HA channel measures total homology-arm copies (endogenous alleles plus donor-derived homology arms), whereas the transgene (TG) channel measures donor transgene copies. In correctly targeted HDR, the HA:TG relationship is determined by locus and reference ploidy, whereas off-target integrations or extrachromosomal donor increase HA signal without a proportional increase in TG. After Poisson inversion and normalization to the reference gene (Ref), total donor copies per 100 cells are calculated from TG/Ref, and off-target or extrachromosomal donor copies are inferred from the HA–TG difference as summarized in the equation (grey box). **(c)** Quantification of targeted knock-in and off-target donor copies in SK-N-AS and SK-N-BE2c cells edited with different HDR donors (RPLP0-CXCL10, RPLP0-CXCL11, AAVS1-CXCL10, CHD1-CXCL10) before and after sorting using the homology-arm method. **(d)** Off-target donor-to-targeted knock-in ratios in unsorted and sorted neuroblastoma populations, showing increased off-target donor load after enrichment of high expressors. **(e)** Schematic of the translocation assay: independent gRNA-induced DSBs generate reciprocal translocation junctions detected by dPCR using primers flanking newly joined genomic segments. **(f)** Quantification of six expected translocation products in multiplex-edited (3 gRNAs) primary human T cells; five junctions are significantly enriched relative to controls, whereas one shows a non-significant trend. **(g)** Analytical characterization of translocation assays: limit of blank (LoB) and **(h)** diagnostic specificity across translocation targets in edited and control T cells. **Data presentation:** Bar graphs show mean ± SD; dots represent biological replicates (neuroblastoma cell lines or donor-derived primary T-cell samples, as indicated). **Statistical analysis:** (d) two-sided Mann-Whitney test; (f,h) Kruskal-Wallis test with Dunn’s post-hoc correction; (h) specificity calculated relative to median LoB thresholds (CLSI EP17-A2). Exact *p* values are provided in the Source Data file. *p* values: *<0.05, **<0.01, ***<0.001; n.s., not significant. Source data are provided as a Source Data file.

**Fig. 5.**
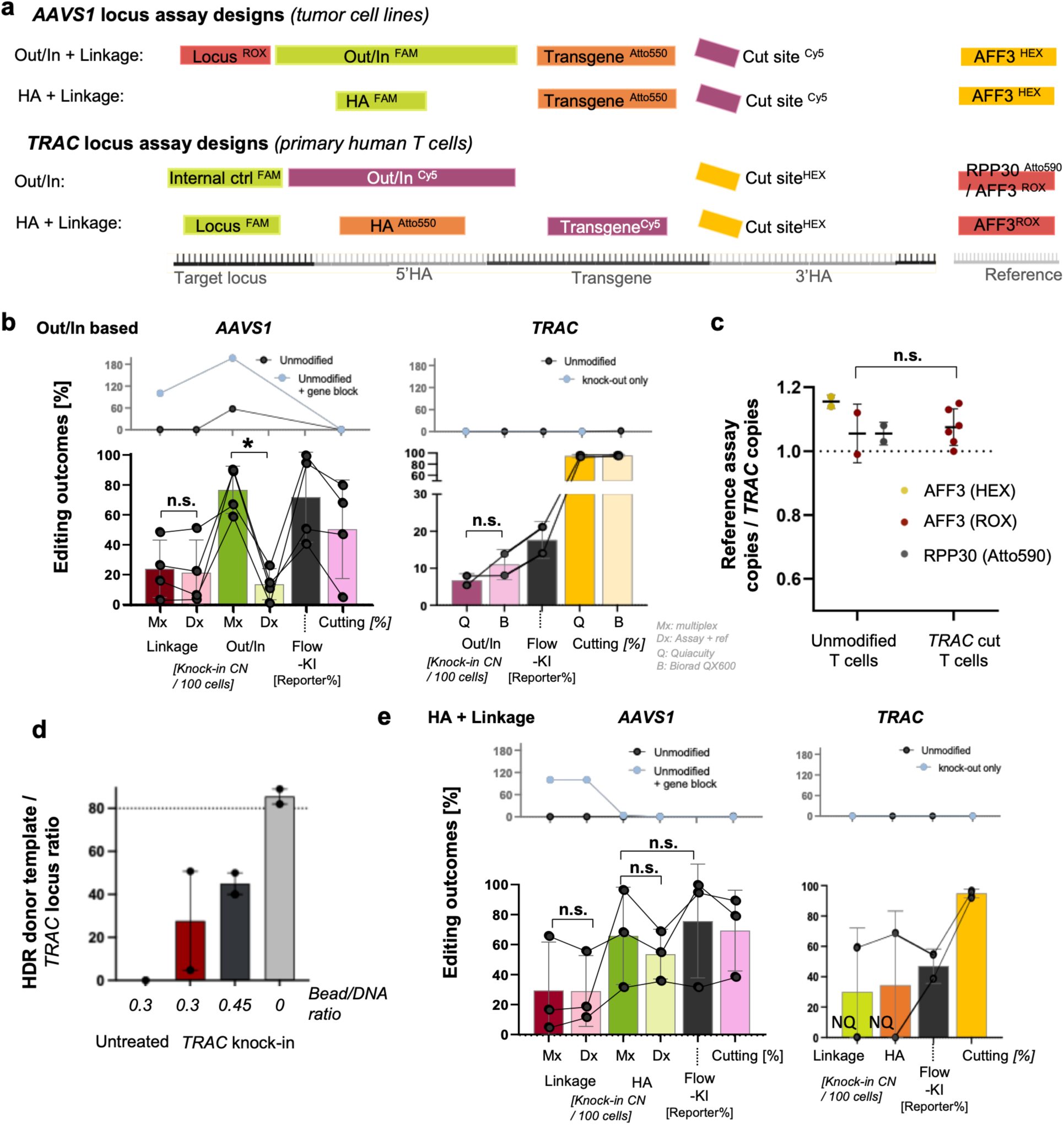
Five-channel multiplex digital PCR enables simultaneous quantification of cutting, CNV, and site-specific knock-in across loci and platforms. (**a**) Schematic of five-channel multiplex assay architectures for *AAVS1* editing in neuroblastoma cell lines and *TRAC* editing in primary human T cells. Two complementary multiplex configurations were implemented for each locus: an Out/In-based panel combining long-range junction, transgene, cut-site and reference assays, and an HA/linkage-based panel replacing the junction assay with homology-arm and linkage measurements. Both architectures enable simultaneous quantification of CRISPR cutting, total transgene copy number, copy-number normalization and site-specific knock-in within a single reaction. (**b**) Evaluation of Out/In-based assay panels for *AAVS1* and *TRAC* editing. For *AAVS1*, lower panels compare five-channel multiplex measurements (Mx) with the corresponding duplex measurements (Dx; assay of interest plus endogenous reference assay). For *TRAC*, lower panels compare Out/In and cut-site measurements across the QIAcuity and Bio-Rad QX600 platforms, together with flow-cytometric readouts. Upper panels show apparent editing outcomes in unedited genomic DNA with or without spike-in of a synthetic knock-in gene block for *AAVS1*, and in unedited or knock-out-only T-cell DNA for *TRAC*. **(c)** Comparison of internal *TRAC* and external reference assays (*AFF3* and *RPP30*) for copy-number normalization in unmodified and *TRAC*-edited primary human T cells. **(d)** Residual HDR donor-template abundance relative to the *TRAC* locus following SPRI bead enrichment at the indicated bead-to-DNA ratios in edited primary T-cell samples. **(e)** Evaluation of linkage-and HA-based assay panels for *AAVS1* and *TRAC* editing. For *AAVS1*, lower panels compare Mx with Dx measurements. For *TRAC*, lower panels show linkage-and HA-based knock-in measurements together with flow-cytometric and cut-site readouts in edited primary T cells. Upper panels show apparent editing outcomes in the corresponding negative-control samples. NQ indicates measurements that could not be reliably interpreted because of insufficient depletion of residual donor template. **Data presentation:** Mean ± SD; dots represent biological replicates. **Statistical analysis:** Dunn‘s multiple comparisons test (d), Mann-Whitney test (c); exact p values are provided in the Source Data. n.s., not significant. Source data are provided as a Source Data file.

To determine whether internal genomic loci could replace conventional external reference genes for copy-number normalization, we compared an internal *TRAC* reference with *AFF3* and *RPP30* in unmodified and edited primary human T cells. Internal and external reference assays showed an approximately 1:1 relationship across all conditions, demonstrating that appropriately selected endogenous loci can provide reliable relative copy-number normalization (**Fig. 5c**). Application of the multiplex workflow to electroporated primary T cells revealed a distinct analytical challenge. Shortly after electroporation, these samples contained large amounts of residual extrachromosomal HDR donor DNA (**Fig. 5d**). As predicted from the analytical characterization (**Fig. 2e**; **Supplementary Figs. 6c,d and 11-13),** these donor levels interfered with quantitative interpretation of linkage and homology-arm assays, while increasing false-positive signals in long-range junction assays. To reduce donor-derived analytical interference, we evaluated SPRI bead-based enrichment of genomic DNA by exploiting the preferential recovery of high-molecular-weight genomic DNA over shorter donor molecules. Spike-in titration experiments identified bead-to-DNA ratios that preferentially depleted donor DNA while retaining genomic DNA **(Supplementary Fig. 23).** Application of the optimized protocol to edited primary T-cell samples produced variable results: one sample showed sufficient depletion of residual donor template to permit linkage-and HA-based quantification, whereas the second sample retained too much donor DNA for reliable quantification by either assay (**Fig. 5d,e**). Although insufficient to fully restore quantitative performance in all highly donor-loaded samples, these findings identify residual donor DNA as an important pre-analytical variable for CRISPR-dPCR and demonstrate that selective genomic DNA enrichment can partially mitigate donor-derived analytical interference.

Taken together, these findings establish CRISPR-dPCR as a flexible multiplex framework capable of integrating quantification of CRISPR cutting, total transgene copy number, and site-specific knock-in within a single reaction. They further identify residual donor DNA as a critical pre-analytical determinant of assay performance in early post-editing samples and demonstrate that sample preparation constitutes an integral component of robust HDR quantification.

## Discussion

Our study establishes a cohesive digital PCR framework that addresses several persistent analytical challenges in CRISPR editing workflows. By evaluating multiple assay classes side-by-side across diverse loci and cell types, we provide a systematic comparison of assay behavior that has been largely absent from earlier work, which typically focused on isolated modalities such as junction PCR^25^, indel profiling^10,11^, or single-target dPCR assays^12^. This integrated analysis reveals how probe chemistry, amplicon architecture, partition statistics, and multiplexing constraints jointly shape performance. The LNA-enhanced cut-site assay demonstrates how modest changes in probe affinity markedly improve cluster separation and quantitative precision. These improvements echo prior observations that high-affinity probes enhance detection of rare variants in dPCR^22^, and our cross-platform consistency highlights the robustness of probe drop-off assays for high-sensitivity cutting quantification. For site-specific knock-in quantification, rather than identifying a single universally optimal strategy, our study demonstrates that each assay exhibits complementary analytical strengths and limitations. Out/In-In/Out assays behaved as expected for long-amplicon designs: highly specific when successful but increasingly prone to rain, and reduced efficiency at longer product lengths, with an additional risk of false-positive signal caused by template switching^25^. To overcome these inherent limitations, we developed two alternative dPCR strategies: the linkage assay and the homology-arm subtraction assay. Linkage assays, grounded in Poisson co-localization statistics, provided reliable estimates when template input and partition occupancy were within an appropriate range. The linkage assay bypasses long-range PCR and provides highly specific orthogonal evidence of site-specific integration, although its sensitivity depends on DNA integrity, partition number, and target abundance. Homology-arm assays achieved highly accurate quantitative measurements in clonal cell populations with minimal residual donor DNA, but were susceptible to false-positive estimates when substantial residual or unintended donor-derived DNA was present.

These findings support a tiered analytical workflow. For routine quantitative analysis after sufficient clearance or depletion of residual donor DNA, we recommend the rapid short-amplicon multiplex approach combining cut-site, transgene, reference, linkage, and homology-arm assays. In this configuration, linkage analysis provides evidence of physical association between the target locus and cargo, whereas homology-arm subtraction provides an independent quantitative estimate and identifies donor-derived signal not explained by correctly integrated alleles. When direct confirmation of the expected genome-insert boundary is required, or when linkage sensitivity is limited by very low target abundance, an adequately optimized and validated Out/In or In/Out assay can be added as a separate orthogonal measurement. Assay performance also depended strongly on sample composition. Residual extrachromosomal donor DNA, which is abundant shortly after ex vivo electroporation, interfered with quantitative interpretation of homology-arm and linkage assays while increasing false-positive signals in long-range junction assays. This limitation was largely absent in clonally expanded populations, where residual donor molecules had been reduced through cell division. Accordingly, samples with substantial residual donor DNA should undergo delayed analysis or selective genomic DNA enrichment before site-specific knock-in measurements are interpreted quantitatively.

Because most assays were evaluated under standardized, minimally optimized conditions, their performance provides realistic expectations for transfer to new loci, although locus-specific optimization may improve individual assays. Linkage and HA measurements remained stable in five-channel panels, whereas Out/In-In/Out assays showed greater multiplex-associated non-specific signal, potentially reflecting oligonucleotide interactions or competition for reaction components ^26^. Analytical evaluation and reporting followed digital MIQE guidelines and the CLSI EP17-A2 framework for detection capability ^30,31^.

Genome-wide approaches such as GUIDE-seq and HTGTS remain indispensable for discovery-oriented identification of previously unknown off-target cleavage sites and structural rearrangements^18–20^. Once relevant loci and editing outcomes have been defined, amplicon-based NGS enables sequence-resolved characterization and quantification but can be affected by amplification bias, variable read depth, and dropout of larger or complex events^16,17^. Compared with amplicon sequencing, dPCR provides faster absolute quantification with high analytical sensitivity, low input requirements, and minimal computational overhead. These characteristics make dPCR particularly attractive for longitudinal monitoring, process development, and, following appropriate assay validation, time-sensitive product-release testing. Our dPCR assays deliberately trade nucleotide-level resolution for quantitative robustness, positioning them as complementary tools rather than replacements for NGS. CRISPR-dPCR fills the emerging need for standardized quantitative assays that accompany targeted knock-in therapies throughout development, manufacturing, and clinical translation. Additionally, while validated here in CRISPR-Cas9 HDR workflows, several components of the framework are not intrinsically dependent on CRISPR or HDR. In particular, junction and linkage assays quantify defined genome-cargo associations and could, in principle, be adapted to targeted integrations generated using other programmable nucleases or site-specific recombinases. By contrast, the homology-arm subtraction assay depends on donor architectures containing homologous sequences and would require modification for non-HDR integration systems.

Recent advances in in vivo targeted genome editing demonstrate that precise integration is becoming feasible beyond ex vivo cell engineering ^9^. Unlike electroporation, which transiently exposes individual cells to a large excess of donor template, systemic in vivo delivery is expected to generate lower intracellular donor copy numbers and faster clearance of residual donor molecules, thereby reducing the donor-derived analytical interference observed shortly after ex vivo editing. Orthogonal CRISPR-dPCR measurements may therefore be particularly well suited to quantitative assessment of in vivo targeted knock-in therapies. More broadly, the CRISPR-dPCR framework integrates measurements of CRISPR cutting, donor abundance and site-specific knock-in, with modular extensions for defined unintended editing outcomes to support applications across therapeutic development, including donor biodistribution, tissue-specific integration and transgene persistence in IND-enabling studies, as well as rapid, low-input longitudinal monitoring after clinical translation. By combining complementary analytical principles whose performance depends on assay architecture and sample composition, CRISPR-dPCR provides a flexible and scalable foundation for genome-editing research, manufacturing and clinical monitoring of therapies based on targeted knock-in.

## Author contributions

ML, LOR, VG, AN, SB, MS, SA, JM, CJQ and JJC performed the experiments. ML, LOR, VG, and AN analysed the data. ML, LOR, VG, AN, SB, LMH, MS, and DLW developed the methodology. ML and LOR wrote the manuscript with input from all authors. ML, DNN, RCW, HED, DLW and AK supervised the study. ML developed the linkage-and homology-arm-based site-specific knock-in quantification assays and methods and conceived and designed the study.

## Data and code availability

All source data underlying the figures and statistical analyses in this study are provided with the manuscript. All R scripts used for simulation and analysis, as well as an optional Shiny interface for the linkage assay, are available at Zenodo (DOI: **10.5281/zenodo.21432198**).

Reviewer access: Zenodo repository access link

## Acknowledgments

The work was supported by Berliner Krebsgesellschaft (grant number LAFF202008 to M.L.) and KINDerLEBEN e.V. Berlin (to M.L.). M.L. was supported by the German Research Foundation (DFG, Deutsche Forschungsgemeinschaft) through a Walter Benjamin Fellowship (Project No. 571675766). Additional support was provided by Charité -Universitätsmedizin Berlin and the Berlin Institute of Health (BIH) at Charité -Universitätsmedizin Berlin (AdHoc Booster Grant to D.L.W. and M.L.). The project was partly supported by European Union under Grant Agreement Nr. 101057438 (https://www.genetiga-horizon.eu/) to D.L.W. Views and opinions expressed are however those of the author(s) only and do not necessarily reflect those of the European Union or the European Health and Digital Executive Agency (HADEA). Neither the European Union nor the granting authority can be held responsible for them. M.L. participates in the BIH Charité Clinician Scientist Program, funded by Charité -Universitätsmedizin Berlin and the BIH. A.K. participates in the BIH Charité Advanced Clinician Scientist Pilot Program, also funded by Charité -Universitätsmedizin Berlin and the BIH. The authors thank Nikolaus Rajewsky for his support as a scientific mentor within the framework of the BIH Charité Clinician Scientist Program.

