## Supplement for "CRISPR-dPCR: A unified multiplex digital PCR framework for quantitative characterization of targeted knock-in"

*(First submission, August 2026)*

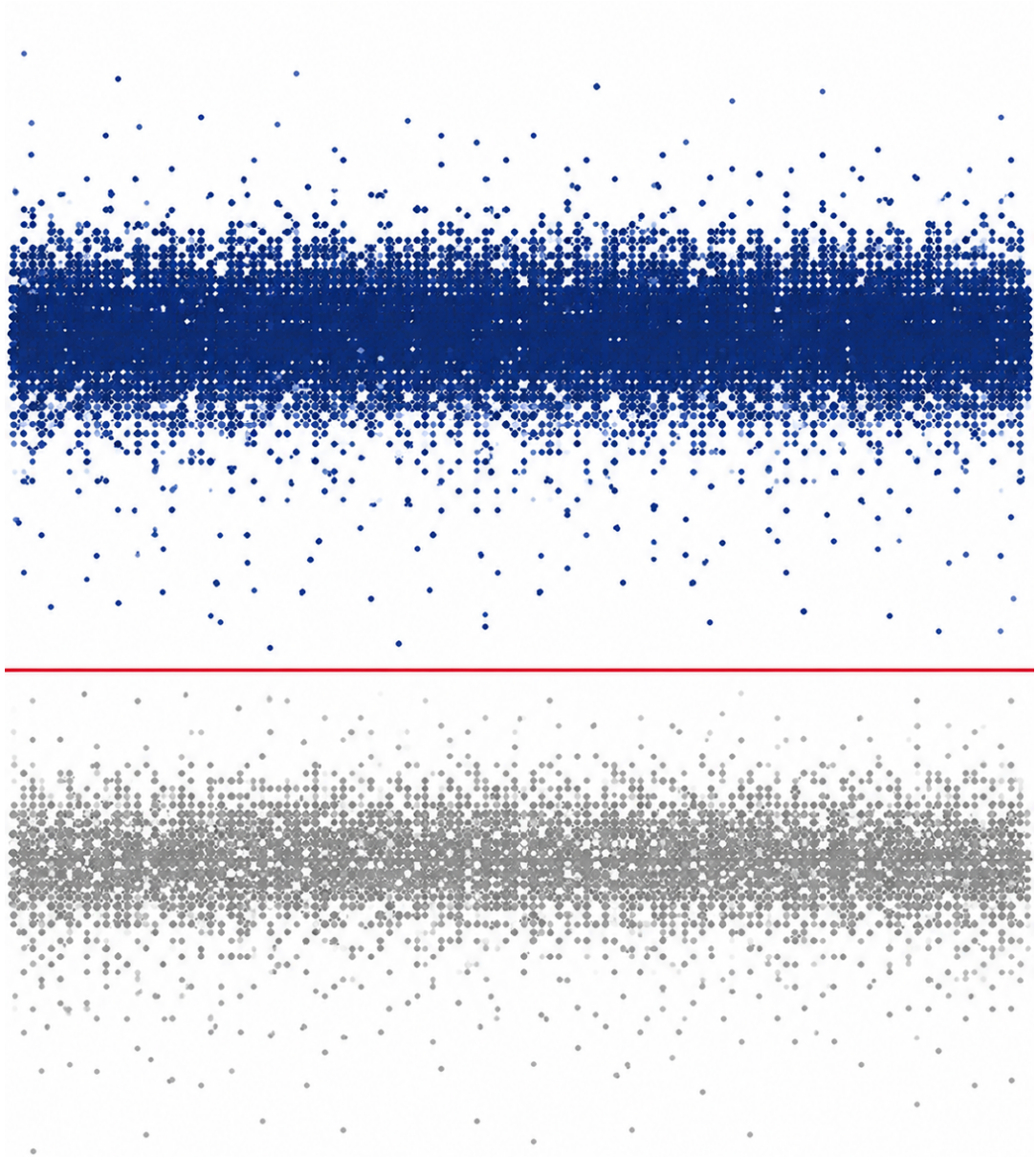

Supplementary Fig. 24

**Cover artwork.**

Rendering based on representative digital PCR droplet distribution generated in this study.
The artwork preserves the characteristic distribution of the experimental data
while applying only minor graphical adjustments for publication purposes.

#### List of Abbreviations

1D / 2D - one- / two-dimensional fluorescence amplitude plots

AAV - adeno-associated virus

AAVS1 - adeno-associated virus integration site 1

AFF3 - AF4/FMR2 family member 3 (reference gene)

BHQ1 / BHQ2 - Black Hole Quencher 1 / 2

CAR - chimeric antigen receptor

Cas9 - CRISPR-associated protein 9

CD - cluster of differentiation

CHD1 - chromodomain-helicase-DNA-binding protein 1

CN - copy number

CNV - copy-number variation

CRISPR - clustered regularly interspaced short palindromic repeats

CXCL10 - C-X-C motif chemokine ligand 10

CXCL11 - C-X-C motif chemokine ligand 11

CXTM - cross-talk matrix

Cy5 / FAM / HEX / ATTO550 / ATTO590 - fluorophores used for dPCR probes

dPCR - digital polymerase chain reaction

DSB - double-strand break

EF1α - elongation factor 1 alpha promoter

FCS - fetal calf serum

FWD / REV - forward / reverse primer

gDNA - genomic DNA

gRNA - guide RNA

HA - homology arm / homology-arm assay

HDR - homology-directed repair

HDRT - homology-directed repair template

HUVEC - human umbilical vein endothelial cell

ICE - Inference of CRISPR Edits

IDAA - indel detection by amplicon analysis

IFNG - interferon gamma

KI - knock-in

LHA / RHA - left / right homology arm

LoB - limit of blank

LoD - limit of detection

LNA - locked nucleic acid

Linkage assay - WT-TG co-localization knock-in method

MIQE - Minimum Information for Publication of Quantitative Digital PCR Experiments

MLE - maximum-likelihood estimation

Mx / Dx - multiplex / dualplex assay configuration

NAGK - N-acetylglucosamine kinase (reference gene)

NC - negative control

NGS - next-generation sequencing

NHEJ - non-homologous end joining

NTC - no-template control

Out/In / In/Out - long-range junction assay orientations

PBMC - peripheral blood mononuclear cell

P2A - self-cleaving 2A peptide

PGA - poly-L-glutamic acid

PoD - probability of detection

qPCR - quantitative polymerase chain reaction

RPP30 - ribonuclease P/MRP subunit p30 (reference gene)

ROX - carboxy-X-rhodamine fluorophore

RNP - ribonucleoprotein

RPLP0 - ribosomal protein lateral stalk subunit P0

SD - standard deviation

SSKI - site-specific knock-in

T - transgene (dPCR channel)

T7E1 - T7 endonuclease I

TIDE - Tracking of Indels by Decomposition

TG - transgene assay

TRAC - T cell receptor $\alpha$ constant locus

WT - wild type

#### Supplementary Notes

##### Supplementary Note 1 - Reference gene copy number variation

Accurate quantification of HDR-mediated knock-in (KI) of large inserts (>400 bp) requires correction for locus-level copy number variation (CNV), which can be substantial in tumor-derived cell models such as the neuroblastoma lines used here. To ensure correct site-specific knock-in (SSKI) quantification, we systematically determined target and reference CNVs using HiC profiling and dPCR (Supplementary Fig. 3a). In SK-N-AS cells, HiC data showed two copies per cell for all evaluated loci (AFF3, NAGK, AAVS1, CHD1) except RPLP0, which exhibited three copies (Supplementary Fig. 3b). In contrast, SK-N-BE2c cells displayed only 1-1.5 copies per cell for the internal controls AFF3 and NAGK on chromosome 2, while maintaining two copies for AAVS1, RPLP0, and CHD1. Direct comparison of HiC and dPCR measurements revealed that the AFF3:AAVS1 ratio deviated from the expected 1:1. Specifically, the AFF3:AAVS1 ratio of 2-3:1 observed by dPCR indicated that the AAVS1 assay captured only one of the two alleles. This matched the raw HiC evidence and would have caused systematic underestimation of AAVS1-based knock-in values; thus, locus-specific correction was applied for subsequent SSKI quantification (Supplementary Fig. 3c).

##### Supplementary Note 2 - Simulation-Based Evaluation of Linkage Assay Sensitivity and Specificity

We performed extensive Monte Carlo simulations to evaluate linkage assay performance across a wide range of input conditions, including varying target copy number, knock-in fraction, partition occupancy, and off-target donor levels (Supplementary Figs. 11, 12, 14a). The simulations demonstrated that linkage assay sensitivity increases monotonically with partition count due to improved stabilization of double-positive (WT+TG) estimates. Merging data from three or more wells significantly enhanced detection of low-frequency knock-ins while preserving high specificity, even at elevated occupancy. False-positive rates remained low across all tested conditions, provided that occupancy remained below saturation thresholds. These results establish partition count as a key determinant of linkage assay performance and support the use of multi-well merging in applications requiring maximal analytical depth.

##### Supplementary Note 3 - Simulation-Based Evaluation of Homology-Arm Assay Performance

We simulated homology-arm (HA) assay behavior across varying occupancy, copy-number inputs, knock-in fractions, and free-donor backgrounds (Supplementary Figs. 13, 14b). Unlike assays reliant on absolute droplet counts, the HA method uses relative arithmetic between the transgene and homology-arm channels combined with Poisson correction, enabling stable accuracy even at high occupancy. Performance degradation occurred only when occupancy exceeded ~99.9%, at which point partition saturation prevented reliable estimation of relative target ratios. These results highlight the intrinsic robustness of the HA approach across a broad dynamic range and support its application in workflows involving high template loads or variable DNA integrity.

##### Supplementary Note 4 - Theoretical and Practical Considerations for HA-Based Quantification of Off-Target Knock-Ins

CRISPR-induced HDR using double-stranded donor templates generates both intended on-target knock-ins and unintended donor-derived products, including off-target integrations and residual extrachromosomal donor DNA. The homology-arm (HA) assay detects all molecules carrying the donor-derived homology arm, irrespective of genomic context, whereas the transgene (TG) assay detects only molecules containing the inserted transgene cassette. After converting droplet positivity to absolute copies per $\mu\text{L}$ via Poisson inversion, the total number of donor-derived molecules is expressed per 100 cells by normalizing TG copies to the reference gene (Ref):

$$\text{Total donor CN per 100 cells} = 100 \times n \times (\text{TG} / \text{Ref})$$

where TG and Ref denote copies per $\mu\text{L}$ for the transgene and reference assays, and $n$ is the reference-gene copy number per cell ($n = 2$ for diploid loci). The on-target knock-in copy number per 100 cells is obtained separately using the HA knock-in equation described in the Methods, which subtracts excess HA signal to isolate true HDR events. Off-target or extrachromosomal donor copies are then calculated as:

Off-target integration/extrachromosomal donor CN per 100 cells = Total donor CN – Knock-in CN

This framework enables simultaneous quantification of HDR precision and unintended donor integration within a single dPCR assay.

###### *Necessary assumptions*

- Equivalent amplification efficiencies: The HA assay must amplify genomic homology-arm sequences and donor-derived homology-arm copies with similar efficiency; otherwise HA-TG differences will not accurately reflect unintended integrations.
- Stable normalization to the reference gene: DNA extraction and input must not differentially bias HA or Ref signals. If the reference gene exhibits variable ploidy—common in neuroblastoma and other cancer lines—$n$ must be measured once in the parental line and used consistently.
- Impact of truncated donor fragments: Donor fragments lacking the TG assay site reduce TG counts and may lead to underestimation of total donor load. In such cases, quantifying both LHA and RHA or including a donor-backbone assay improves accuracy.
- Distinguishing HDR from non-HDR donor capture: Because the HA assay cannot differentiate correct HDR junctions from illegitimate donor capture, junction assays targeting genome-insert boundaries are required for definitive confirmation of on-target HDR.

##### **Supplementary Note 5 - Homology Arms-based site-specific knock-in quantification: Full Derivation, Formula, and Assumptions**

###### *Overview:*

The homology-arm (HA) assay quantifies HDR-mediated knock-in by comparing three independently measured digital PCR channels:

T = Transgene CN/μL

H = Homology-arm CN/μL

R = Reference gene CN/μL

The HA channel reports both true HDR integrations and unintended donor-derived signals (free donor or off-target donor capture). In contrast, the transgene channel reflects only insert-containing DNA molecules. The reference channel provides a stable denominator for estimating cell equivalents and normalizing copy numbers.

###### *1. Poisson inversion for absolute dPCR quantification*

For each channel, the absolute concentration is computed from the observed fraction of positive partitions ($p$) and the partition volume ($V$):

$$\text{copies\_per\_}\mu\text{L} = -\ln(1 - p) / V$$

This relationship is applied independently to transgene (T), homology-arm (H), and reference (R) assays.

###### *2. Expected signal structure*

In unedited diploid cells:

$R = n$ copies per cell, with $n = 2$ for a diploid reference locus.

$H = k \times R$, where $k$ is the expected HA:reference ratio in wild-type cells. For most diploid loci, $k \approx 1$.

T = 0.

After HDR knock-in:

T = HDR\_copies + donor\_noise

H = HDR\_copies + donor\_noise

Thus, any excess HA signal beyond its expected wild-type baseline:

excess\_H = H – k × R represents unintended donor-derived molecules (off-target knock-ins or residual donor DNA).

##### 3. Core derivation - final KI formula used in the manuscript

True HDR estimate

true\_HDR = T – excess\_H

Cell-equivalent denominator: Cells per $\mu\text{L}$ are estimated from the reference locus as

cells\_per\_$\mu\text{L}$ = R / n, with n = 2 in diploid samples.

Final knock-in formula (as shown in the manuscript and in Figure 2c):

$$\text{Knock-in CN per 100 cells} = 100 \times \left( \frac{\text{Transgene CN}/\mu\text{L} - \max(\text{Homology arm CN}/\mu\text{L} - k \times \text{Reference gene CN}/\mu\text{L}, 0)}{(\text{CN}/\mu\text{L Reference gene} : n)} \right)$$

This expression is the operational formula applied throughout the manuscript and simulations.

##### 4. Assumptions

- Amplification parity: HA and reference assays must have comparable amplification efficiencies under dPCR conditions.
- Stable reference locus: The reference gene is assumed to be unaffected by CRISPR genome editing.
- Donor truncation effects: Truncated donor fragments may yield HA<sup>+</sup> / T<sup>-</sup> species. This increases excess\_H, reducing the inferred KI. This bias is conservative, preventing overestimation.
- Ploidy dependence: For aneuploid lines (e.g., neuroblastoma), k and n were empirically determined once in parental cells and used in all downstream calculations.
- High-occupancy robustness: HA quantification remains accurate even at high partition occupancy because the KI estimate relies on relative channel arithmetic, not droplet-by-droplet counting. Failure occurs only at extreme saturation (>99.9% positivity).

##### 5. Failure or edge-case handling

If H < kR:

Excess donor is interpreted as zero; the algorithm clips using

max(H – kR, 0)

to avoid negative off-target estimation.

High donor contamination: When donor-noise ≫ HDR, KI estimates approach zero, reflecting biological reality (no detectable integration at the target locus).

Partition saturation:

If any channel exceeds ~99.9% positivity, Poisson inversion becomes unstable.

These rare cases were excluded or flagged during analysis.

#### Supplementary Note 6 - Linkage method for site-specific knock-in quantification- Mathematical Framework and Maximum-Likelihood Derivation

##### Overview:

The linkage assay quantifies HDR-mediated knock-in by detecting an excess of FAM<sup>+</sup>ROX<sup>+</sup> double-positive partitions beyond that expected from independent Poisson loading of wild-type (WT) and transgene (TG) molecules. True HDR integration generates linked WT-TG molecules that always co-occupy a partition, whereas unlinked WT-only and TG-only molecules distribute independently.

The linkage analysis proceeds in two stages:

(1) Hypothesis testing (binomial test) to determine whether observed double positives exceed the expectation under independence.
(2) Maximum-likelihood estimation (MLE) of the mixture fractions of WT-only, TG-only, and linked WT-TG molecules, denoted $a$, $b$, and $c$, respectively.

All quantities and formulas below correspond exactly to the implementation in the R simulation code.

##### 1. Poisson occupancy model

Let:

$P$ = total number of accepted partitions

$p_F$ = FAM-positive fraction = $FAM\_pos / P$

$p_R$ = ROX-positive fraction = $ROX\_pos / P$

$p_{FR}$ = double-positive fraction = $double\_pos / P$

Partition-level Poisson inversion gives:

$$\lambda_F = -\ln(1 - p_F)$$

$$\lambda_R = -\ln(1 - p_R)$$

These $\lambda$ values represent the mean number of TG-only or WT-only molecules per partition if no linkage existed. They are used for the null model ($H_0$).

##### 2. Null hypothesis: independent WT and TG molecules

Under $H_0$ (no linkage: $c = 0$), TG-only and WT-only molecules load independently.

The expected double-positive fraction is:

$$expected\_double\_fraction = 1 - \exp(-\lambda_F) - \exp(-\lambda_R) + \exp(-(\lambda_F + \lambda_R))$$

Expected count:

$$expected\_double\_pos = P * expected\_double\_fraction$$

Statistical significance is evaluated using a binomial test:

$$p\_value = 1 - \text{BinomialCDF}(double\_pos - 1; n = P, p = expected\_double\_fraction)$$

If $p\_value < 0.05$, $H_0$ is rejected and HDR linkage is inferred.

##### 3. Maximum-likelihood estimation of mixture fractions (a, b, c)

If $H_0$ is rejected, the observed partition fractions arise from a three-component Poisson mixture:

a = fraction of WT-only molecules

b = fraction of TG-only molecules

c = fraction of true HDR-linked WT-TG molecules

The R code solves the following MLE equations obtained by inverting the single- and double-positive probabilities:

Define:

$$a\_plus\_c = -\ln(1 - pF) / \lambda\_total$$

$$b\_plus\_c = -\ln(1 - pR) / \lambda\_total$$

Here $\lambda\_total = \lambda\_hat$ is obtained from the negative-partition fraction:

$$p\_none = negative\_partitions / P$$

$$\lambda\_total = -\ln(p\_none)$$

Then the mixture parameters are:

$$c = (a\_plus\_c + b\_plus\_c - 1)$$

$$a = a\_plus\_c - c$$

$$b = b\_plus\_c - c$$

This solution corresponds to the maximum-likelihood estimate of a, b, c under a shared Poisson occupancy parameter $\lambda\_total$.

##### 4. Conversion to absolute copy numbers and knock-in frequency

Using Poisson inversion, compute per-μL concentrations:

$$FAM\_total\_copies = -\ln(1 - pF) / V$$

$$ROX\_total\_copies = -\ln(1 - pR) / V$$

$$HEX\_total\_copies = -\ln(1 - pHEX) / V$$

where V is the platform-specific partition volume.

Because each linked WT–TG molecule contributes one copy to both the FAM and ROX measurements, linked molecules are counted twice in the summed FAM and ROX copy concentrations. The conversion factor applied to the summed target-copy concentration is therefore:

$$linked\_target\_copy\_fraction = c / (a + b + 2c)$$

The absolute concentration of linked molecules is:

$$linked\_copies\_per\_uL = linked\_target\_copy\_fraction * (FAM\_total\_copies + ROX\_total\_copies)$$

For a diploid reference present at two copies per cell, cell equivalents per μL are:

$$cell\_equivalents\_per\_uL = HEX\_total\_copies / 2$$

Final knock-in frequency per 100 cells:

$$KI\_per\_100\_cells = 100 * (linked\_copies\_per\_uL / cell\_equivalents\_per\_uL)$$

##### 5. Approximate 95% confidence interval

The R code uses a binomial/normal approximation based on the variance of c:

$$CI\_95 \approx 1.96 * \sqrt{c * (1 - c) / P} * scaling\_factor$$

where the scaling factor rescales the variance of $c$ into units of knock-in per 100 cells.

###### 6. Assumptions and boundary conditions

- Molecules follow Poisson loading across partitions.
- Reference gene is diploid.
- Channels have similar amplification efficiency and no cross-talk after correction.
- Partitions are not saturated ($p_F$, $p_R$, $p_{FR} < \sim 0.95$).
- At low molecule counts, uncertainty in $c$ increases; simulations capture this.
- $H_0$ rejection is required before KI estimation (prevents false positives).

###### 7. Failure/edge cases

- Very low FAM copy numbers ($< \text{linked\_molecules}$) → insufficient signal (the R code returns NULL).
- High occupancy $\rightarrow \lambda_{\text{total}}$ estimation unstable; detection unreliable above $\sim 99\%$ positivity.
- No rejection of $H_0 \rightarrow$ KI estimate not reported (no evidence of HDR linkage).

#### Supplementary Methods

##### Quantitative PCR (qPCR)

Quantitative PCR was performed to evaluate primer-probe performance and confirm assay specificity under conventional real-time amplification conditions. Reactions were assembled using the same primer-probe sets employed in digital PCR, together with a probe-based qPCR master mix, in a final volume of 20 $\mu\text{L}$. Genomic DNA or cDNA (3 $\mu\text{L}$ per reaction) was added as template, and all samples were run in technical duplicates. Thermal cycling conditions were matched to those used for digital PCR: assays with short amplicons were amplified using a two-step protocol (95 $^{\circ}\text{C}$ for 2 min followed by 40 cycles of 95 $^{\circ}\text{C}$ for 15 s and 58 $^{\circ}\text{C}$ for 30 s), whereas assays requiring extended amplification followed an equivalent program incorporating an additional 72 $^{\circ}\text{C}$ extension step. Fluorescence was recorded during the annealing/extension phase, and amplification curves and $C_t$ values were inspected to confirm specific amplification, the absence of non-specific signal, and consistent replicate performance. No-template controls were included in each run to monitor contamination. qPCR was used solely to assess assay behavior; all quantitative analyses of genome editing outcomes were performed using digital PCR.

##### Cell Lines and Cell Culture

SK-N-BE(2)c and SK-N-AS neuroblastoma cells were propagated in high-glucose DMEM (Gibco) supplemented with 10% heat-inactivated fetal calf serum and 1% penicillin-streptomycin. Human umbilical vein endothelial cells (HUVECs; ATCC CRL-1730) were maintained in Endothelial Cell Growth Medium 2 (PromoCell) supplemented with EGF, IGF, VEGF, FGF2, 2% fetal bovine serum, and penicillin-streptomycin. All adherent cell cultures were kept at 37 $^{\circ}\text{C}$ in a humidified atmosphere containing 5% $\text{CO}_2$. Peripheral blood lymphocytes were obtained from fresh leukocyte cones from healthy adult donors, and $\text{CD3}^+$, $\text{CD4}^+$, or $\text{CD8}^+$ T cells were enriched using EasyStep magnetic isolation kits (StemCell Technologies). Purified T cells were seeded at $1 \times 10^6$ cells/mL in X-VIVO 15 medium (Lonza) supplemented with 5% human serum, penicillin-streptomycin, and cytokines IL-7 and IL-15 (5 ng/mL each; Miltenyi). Cells were activated for 48 hours using anti-CD3/CD28 Dynabeads (Thermo Fisher Scientific) at a 1:1 bead-to-cell ratio prior to downstream applications.

##### HiC-based detection of copy number variations (CNVs)

HiC libraries were generated using the EpiTect HiC Kit (QIAGEN) according to the manufacturer's instructions, with minor adaptations for adherent cell lines. Briefly, $5 \times 10^5$ cells per reaction were harvested, trypsinized, and crosslinked in suspension using formaldehyde to preserve spatial chromatin interactions. Crosslinking was quenched, cells were washed in PBS, and nuclei were isolated and immobilized on magnetic beads. Chromatin was digested using a restriction endonuclease recognizing GATC sites, followed by fill-in of 5' overhangs with biotinylated nucleotides. Proximal DNA ends were ligated under dilute conditions to favor intranuclear ligation events. Crosslinks were reversed by proteinase K digestion, and DNA was purified. Purified HiC DNA was sheared by focused ultrasonication (Covaris) to a median fragment size of 400-600 bp. Biotin-labeled ligation junctions were captured using streptavidin beads, followed by end repair, A-tailing, and ligation of indexed Illumina sequencing adapters. Libraries were amplified by PCR using cycling conditions recommended by the manufacturer, purified, and quality-controlled using an Agilent 2100 Bioanalyzer. Library concentration was additionally determined by qPCR. Sequencing was performed on an Illumina NextSeq 500/550 platform to a depth of approximately 35-45 million paired-end reads per sample, which was determined empirically to provide sufficient resolution for structural variant and CNV detection while avoiding saturation effects.

HiC data processing and CNV inference: Raw sequencing data were processed using the QIAGEN EpiTect HiC analysis pipeline to generate genome-wide contact matrices. Copy number variation (CNV) analysis was performed using HiNT (HiC for copy number variation and translocation detection)<sup>38</sup>, which infers CNVs directly from HiC read coverage profiles. Briefly, HiNT normalizes HiC contact counts to account for genomic distance, restriction fragment density, and mappability, and derives copy number profiles based on deviations in normalized read depth across genomic bins. CNVs were calculated as $\log_2$ ratios of observed versus expected copy number, with the expected copy number centered at zero for each chromosome. CNV profiles were compared between edited samples and their corresponding wild-type controls to identify gains and losses associated with genome editing. CNV calls were interpreted in conjunction with HiC-based structural variant detection (HiC Breakfinder), allowing correlation of copy number changes with detected intra- and interchromosomal rearrangements. All CNV analyses were performed at consistent resolution and sequencing depth across samples to ensure comparability.

##### Flow Cytometry

Flow cytometry was used to quantify transgene expression, lineage markers and cell viability. Measurements were acquired on a BD LSRFortessa X-20 equipped with 405, 488, 561 and 640 nm lasers, and data were analyzed in FlowJo v10. Cells were washed in PBS and stained in 100 μL of antibody solution; CD34 expression was detected using PE-conjugated anti-CD34 (QBEND/10, Invitrogen, 1:10), and CAR staining was performed following pre-incubation with anti-mouse F(ab')<sub>2</sub> reagent and serum block to minimize non-specific binding. After 30 min incubation at 4 °C, cells were washed and analyzed immediately. Fluorescence compensation was performed using single-stained primary CD8<sup>+</sup> T cells. All analyses followed a consistent gating sequence: exclusion of debris based on FSC/SSC, doublet discrimination, live/dead gating, lineage gating (e.g., CD3<sup>+</sup>), and finally detection of CD34 or CAR expression. Representative gating workflows are provided in **Supplementary Fig. 19**.

##### Custom Cross-Talk Matrix for Multiplex dPCR

To ensure accurate fluorescence discrimination in five-color multiplex dPCR, we generated a custom cross-talk matrix (CXTM) according to the manufacturer's guidelines for the QIAcuity system. Optical cross-talk was quantified by running all assays used in the multiplex panel as simplex reactions under identical cycling and imaging conditions. Simplex wells contained a single fluorophore-labeled probe at template concentrations yielding approximately 25-75% positive partitions, enabling reliable estimation of dye bleed-through into neighboring channels. Additional wells containing master mix only were included to define background fluorescence for all optical channels. Using these data, the software computed channel-specific correction factors that describe the contribution of each fluorophore to all other channels. The resulting cross-talk matrix was saved and applied to the corresponding multiplex reactions, thereby recalculating fluorescence amplitudes on a per-partition basis and reducing false-positive signal populations caused by spectral overlap. Cross-talk matrices were generated separately for each multiplex assay composition, as recommended, and were reapplied only when assay combinations, probe concentrations, or imaging conditions remained unchanged. All procedures followed the manufacturer's instructions. Multiplex panel composition and dye assignment were guided by the optical layout of the QIAcuity system and manufacturer recommendations. Fluorophores were distributed across channels to minimize spectral overlap, with FAM, HEX/ATTO550, ROX and Cy5 reserved for the highest-priority assays (cut-site, transgene, reference, and target-locus/linkage/HA assays) and ATTO590 used only in configurations where sufficient separation from neighboring channels had been verified in simplex runs. Assays expected to generate high-amplitude signals (e.g. reference and target-locus assays) were preferentially placed in channels with minimal bleed-through into lower-amplitude channels. Fluorophore combinations were selected such that no two probes with strongly overlapping emission spectra were assigned to adjacent channels in the same reaction, and candidate dye sets were validated in simplex mode prior to multiplexing. Unless otherwise stated, probe and primer concentrations in multiplex reactions were identical to their simplex counterparts (typically 800 nM primers and 400 nM probe for primary targets, 400 nM primers and 200 nM probe for reference assays). Where necessary to improve cluster separation or reduce cross-talk, probe concentrations of the brightest assays were reduced by up to 50% while maintaining robust amplification, and the final panel configuration was locked once clear positive-negative separation and stable amplitude distributions were achieved across all channels.

#### Supplementary Figures

**a**

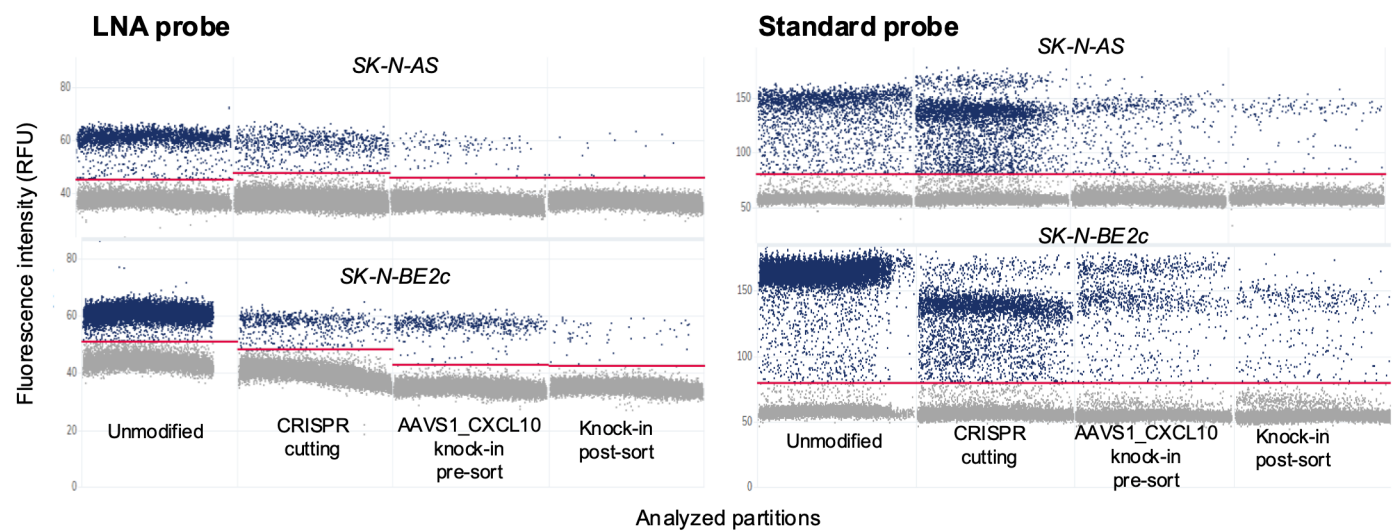

**b**

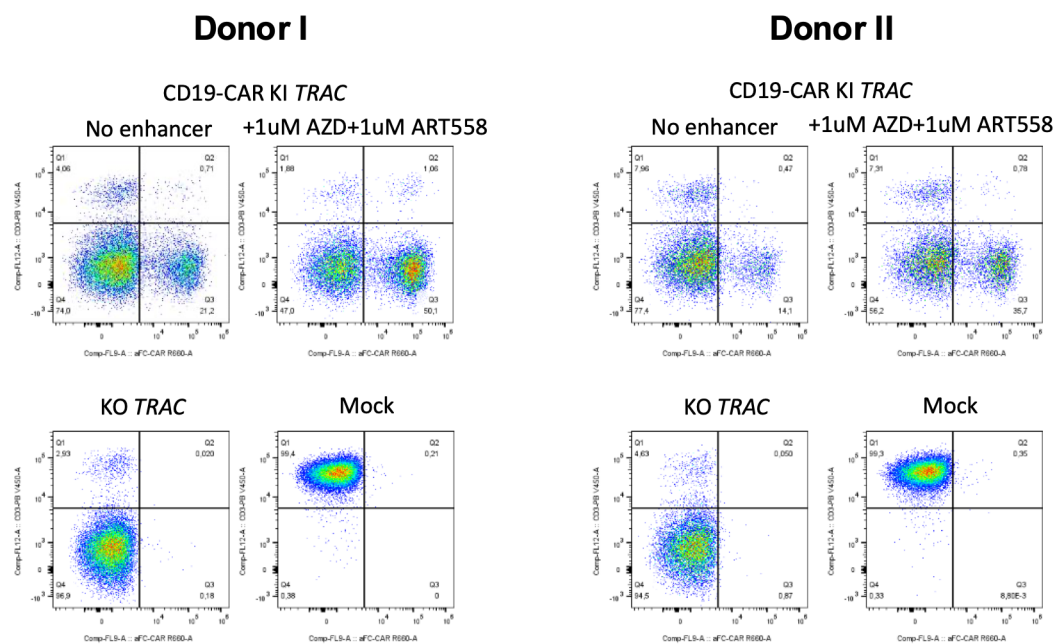

**Supplementary Fig. 1. Representative amplitude plots and flow-cytometry validation for probe drop-off cutting assays.**

**(a)** One-dimensional fluorescence amplitude plots comparing the LNA-modified probe (left) with the standard DNA probe (right) for detection of AAVS1 cutting in SK-N-AS and SK-N-BE2c cells. Plots show unmodified controls, CRISPR cutting-only samples, AAVS1\_CXCL10 knock-in (pre-sort), and knock-in (post-sort). The LNA probe displays tighter positive-negative cluster separation and reduced rain relative to the standard probe across both cell lines. Red dashed lines indicate thresholds. **(b)** Flow cytometry analysis of primary human T cells edited at the TRAC locus. Representative plots from two independent donors show CD19-CAR knock-in (upper panels) with or without HDR enhancers (1 μM AZD + 1 μM ART558), and TRAC knockout (lower panels). Mock-treated cells are shown as negative controls. Enhancer treatment increases the frequency of CAR<sup>+</sup> cells in both donors. **Data presentation:** representative plots from biological replicates. Source data are provided as a Source Data file.

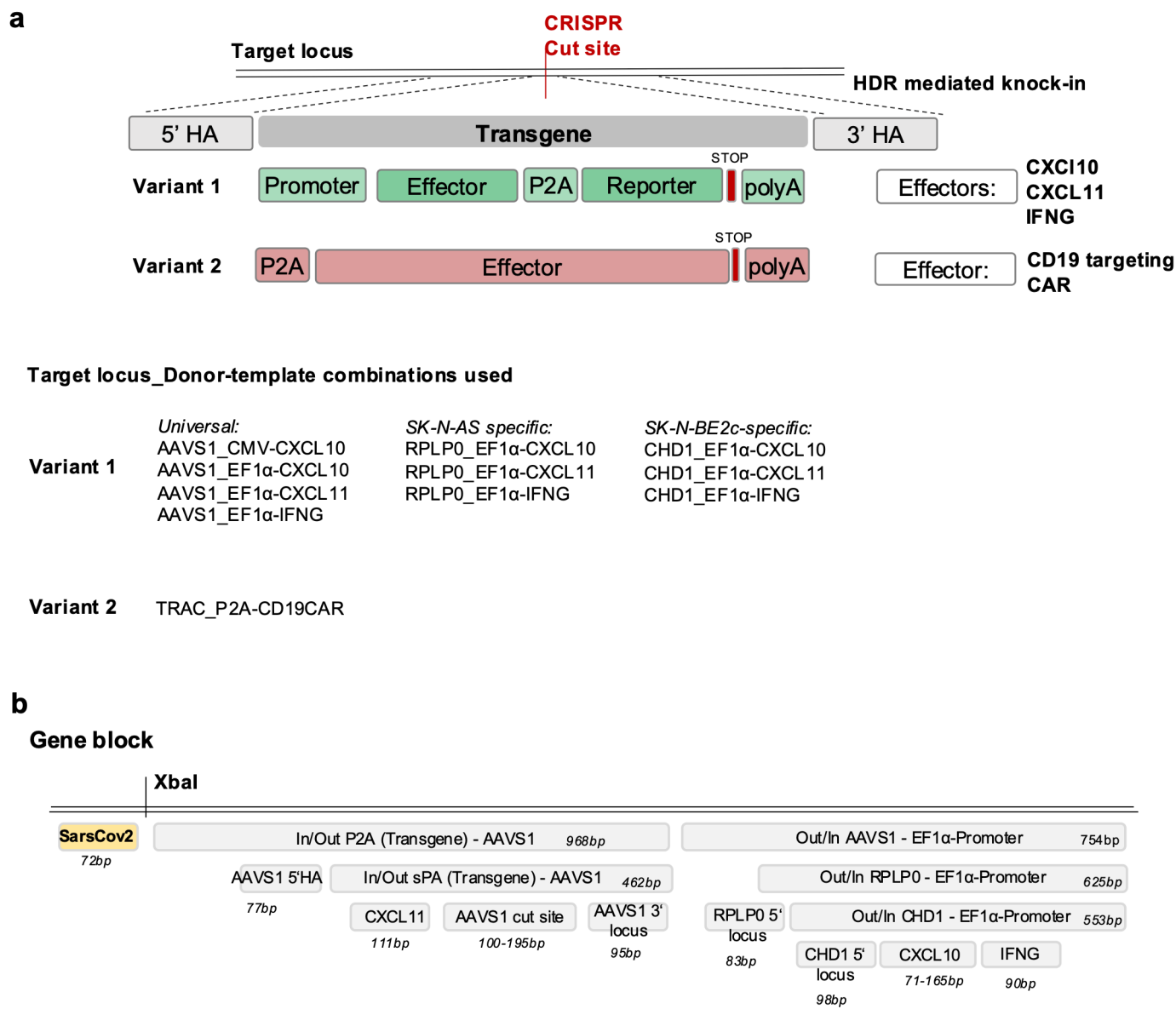

**Supplementary Fig. 2. Donor-template architectures and synthetic gene block used for assay development and analytical benchmarking.**

**(a)** Schematic overview of HDR donor constructs used across assays. Two donor-template variants are shown. Variant 1 contains a promoter-effector-P2A-reporter cassette followed by a polyadenylation signal, while Variant 2 comprises a P2A-linked effector terminating in a polyA sequence. Examples of effector payloads include CXCL10, CXCL11, IFNG, and a CD19-targeting CAR. Target-locus-donor combinations used in SK-N-AS, SK-N-BE2c, and universal AAVS1 contexts are listed. **(b)** Map of the synthetic gene block used for analytical characterization of assay efficiency, specificity, and multiplex behaviour. The construct contains the SARS-CoV-2 N-gene control sequence, an XbaI restriction site, and binding sites for In/Out, Out/In, transgene, homology-arm, and target-locus assays across multiple effector payloads. Amplicon sizes for all assay components are indicated. Source data are provided as a Source Data file.

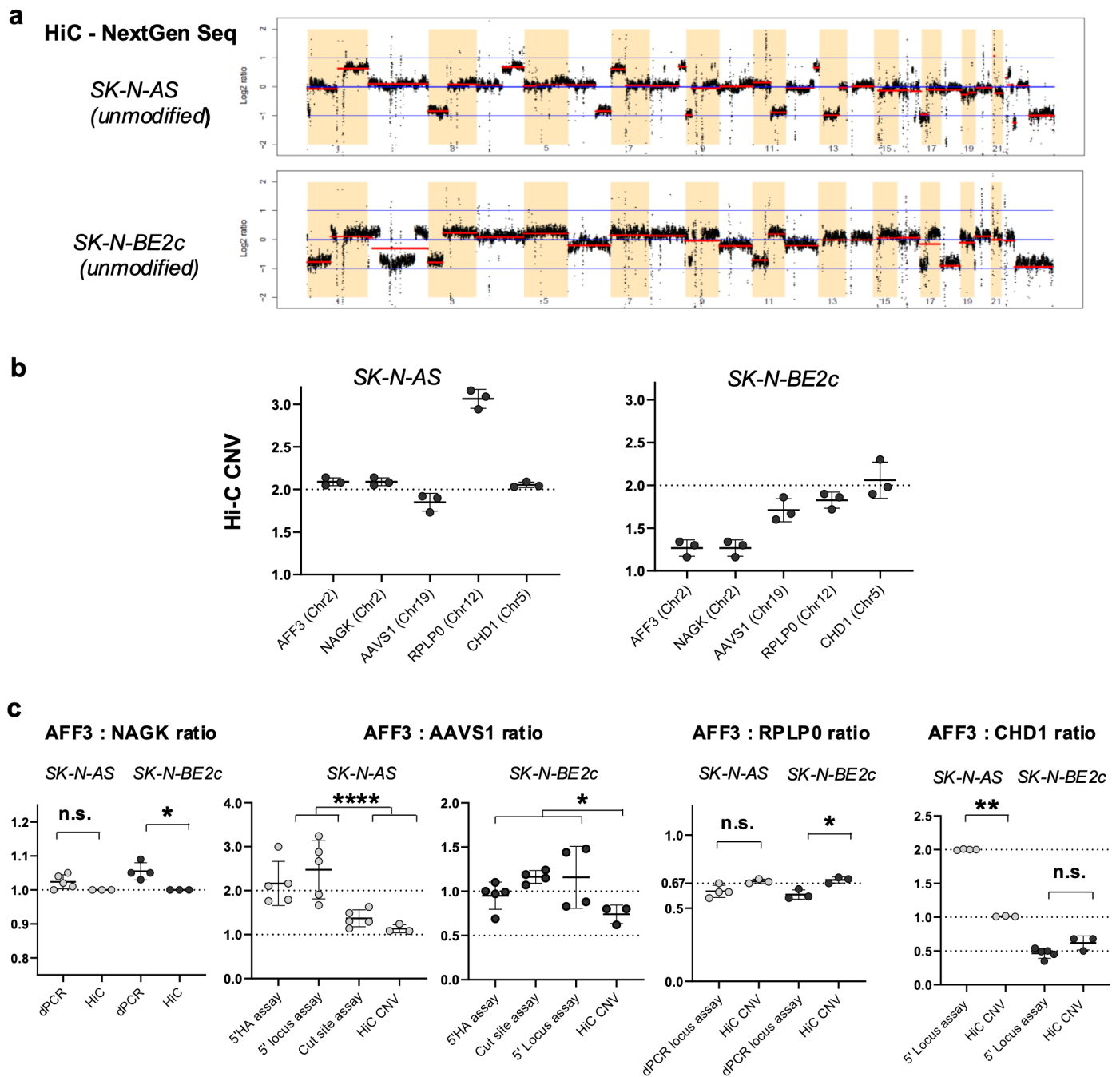

Supplementary Fig. 3. Copy-number variation (CNV) analysis of neuroblastoma cell lines and validation of reference-target ratios for site-specific knock-in quantification.

(a) HiC-derived genome-wide CNV profiles of unmodified SK-N-AS and SK-N-BE2c cells, highlighting broad chromosomal gains and losses that differ between the two neuroblastoma models. (b) HiC-derived absolute CN estimates for key loci used in dPCR assays (AFF3, NAGK, AAVS1, RPLP0, CHD1) showing locus-specific deviations from diploidy in both cell lines, with elevated AAVS1 and RPLP0 copy numbers in SK-N-BE2c and increased CHD1 CN in SK-N-AS. (c) Comparison of AFF3-to-target gene ratios (AFF3:NAGK, AFF3:AAVS1, AFF3:RPLP0, AFF3:CHD1) measured by dPCR (different assays) and HiC SK-N-AS and SK-N-BE2c cell lines. Ratios reveal consistent deviations from the expected diploid reference across loci, confirming the need for cell-line-specific CNV correction when quantifying HDR-mediated knock-in. **Data presentation:** (b,c) mean ± SD; dots represent individual biological or technical replicates. **Statistical tests:** (c) Kruskal-Wallis with Dunn's post-hoc correction; exact p values are reported in the Source Data file. p values: n.s., not significant; \*$<0.05$; \*\*$<0.01$; \*\*\*\*$<0.0001$. Source data are provided as a Source Data file.

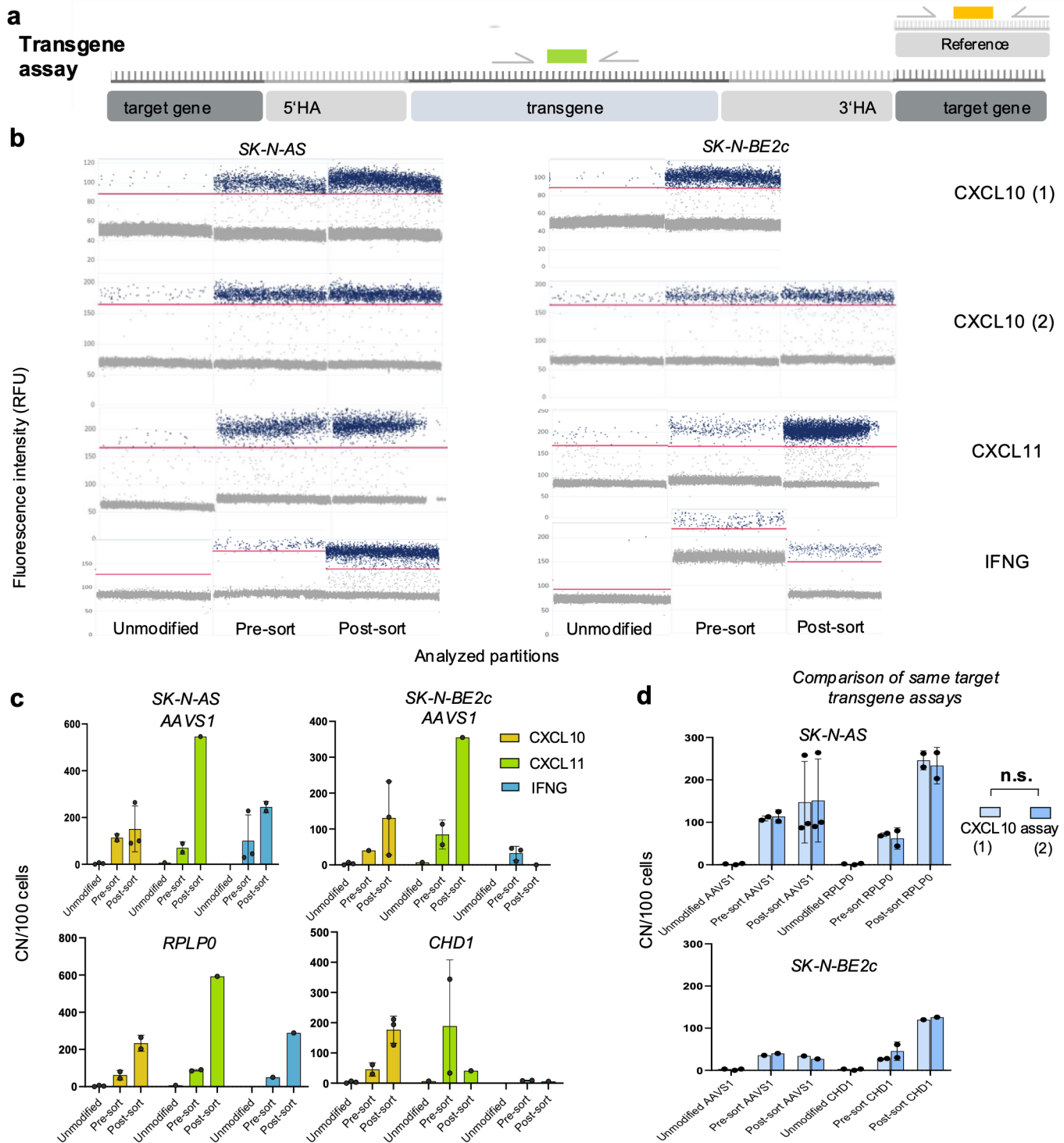

**Supplementary Fig. 4. Performance and concordance of transgene assays across loci, donors, and assay designs.**

**(a)** Schematic of the transgene assay architecture within the integrated transgene, with normalization to an endogenous reference gene. **(b)** Representative amplitude plots for four donor payloads (CXCL10 (assay 1), CXCL10 (assay 2), CXCL11, IFNG) in SK-N-AS and SK-N-BE2c cells across unmodified, pre-sort, and post-sort samples, illustrating robust positive-negative cluster separation and minimal rain across transgene assay designs. **(c)** Comparison of transgene-derived copy numbers per 100 cells across AAVS1-, RPLP0-, and CHD1-targeting knock-ins in SK-N-AS and SK-N-BE2c cells, showing consistent assay performance and expected increases upon sorting, with donor-specific variation in HDR efficiency across payloads. **(d)** Direct comparison of two independent CXCL10 transgene assays (CXCL10 (1) vs. CXCL10 (2)) targeting identical payloads in SK-N-AS and SK-N-BE2c cells, demonstrating near-identical quantitative output and confirming assay interchangeability. **Data presentation:** (c, d) mean ± SD; dots represent biological replicates. **Statistical tests:** (d) Two-way Anova; exact p values are reported in the Source Data file. p values: n.s., not significant; $* < 0.05$. Source data are provided as a Source Data file.

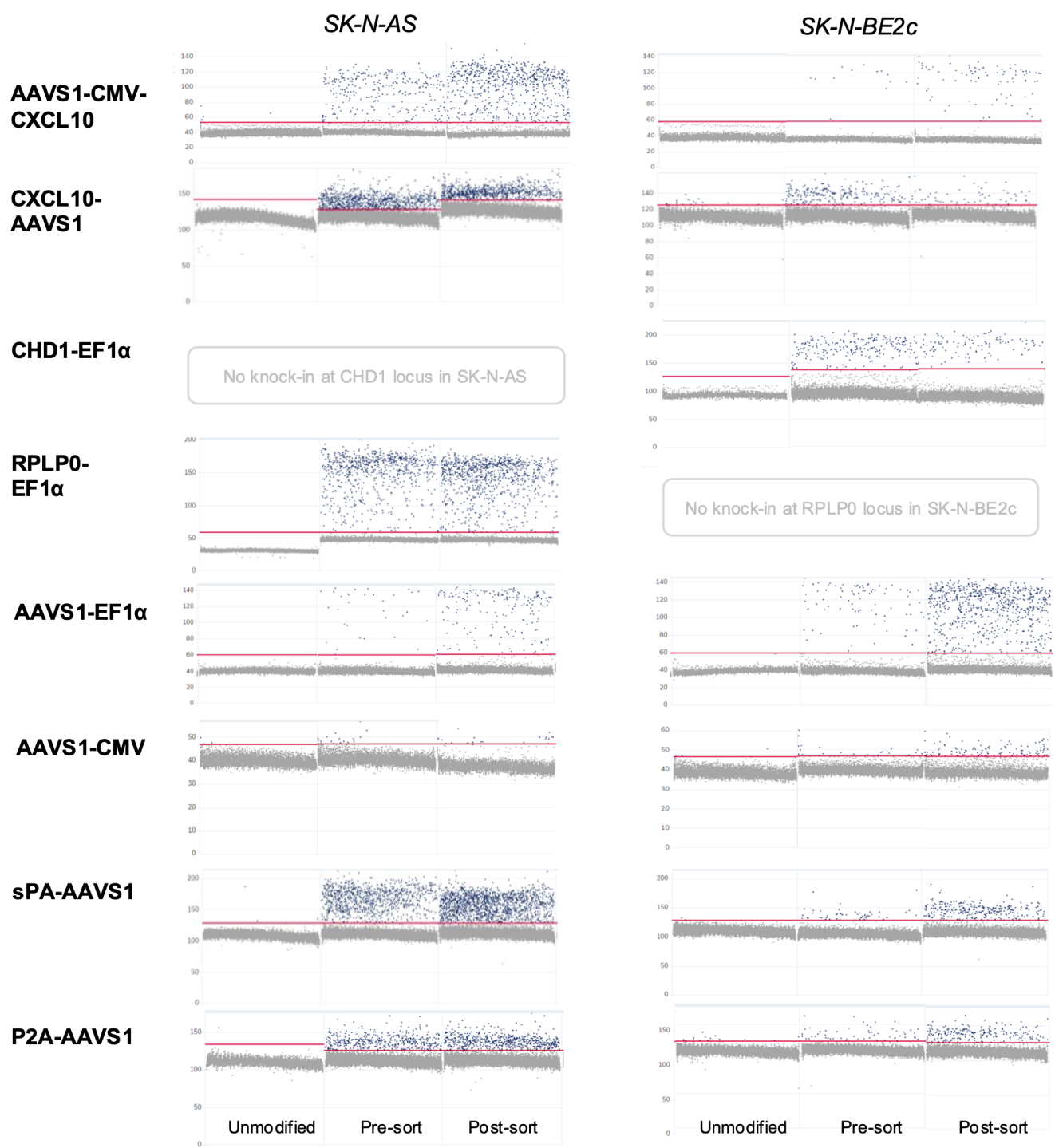

**Supplementary Fig. 5. Transgene assay performance across diverse donor-locus combinations in SK-N-AS and SK-N-BE2c cells.**

**(a)** Representative one-dimensional fluorescence amplitude plots for eight transgene assays spanning multiple donor-locus combinations in SK-N-AS and SK-N-BE2c cells (AAVS1-CMV-CXCL10, CXCL10-AAVS1, CHD1-EF1α, RPLP0-EF1α, AAVS1-EF1α, AAVS1-CMV, sPA-AAVS1, P2A-AAVS1), shown across unmodified, pre-sort, and post-sort populations. Assays targeting loci without confirmed knock-ins (CHD1 in SK-N-AS and RPLP0 in SK-BE2c) display only background-level fluorescence. In all productive edits, amplitude plots demonstrate clear positive-negative cluster separation with minimal rain, confirming robust transgene assay behaviour across donors, promoters, and genomic contexts.

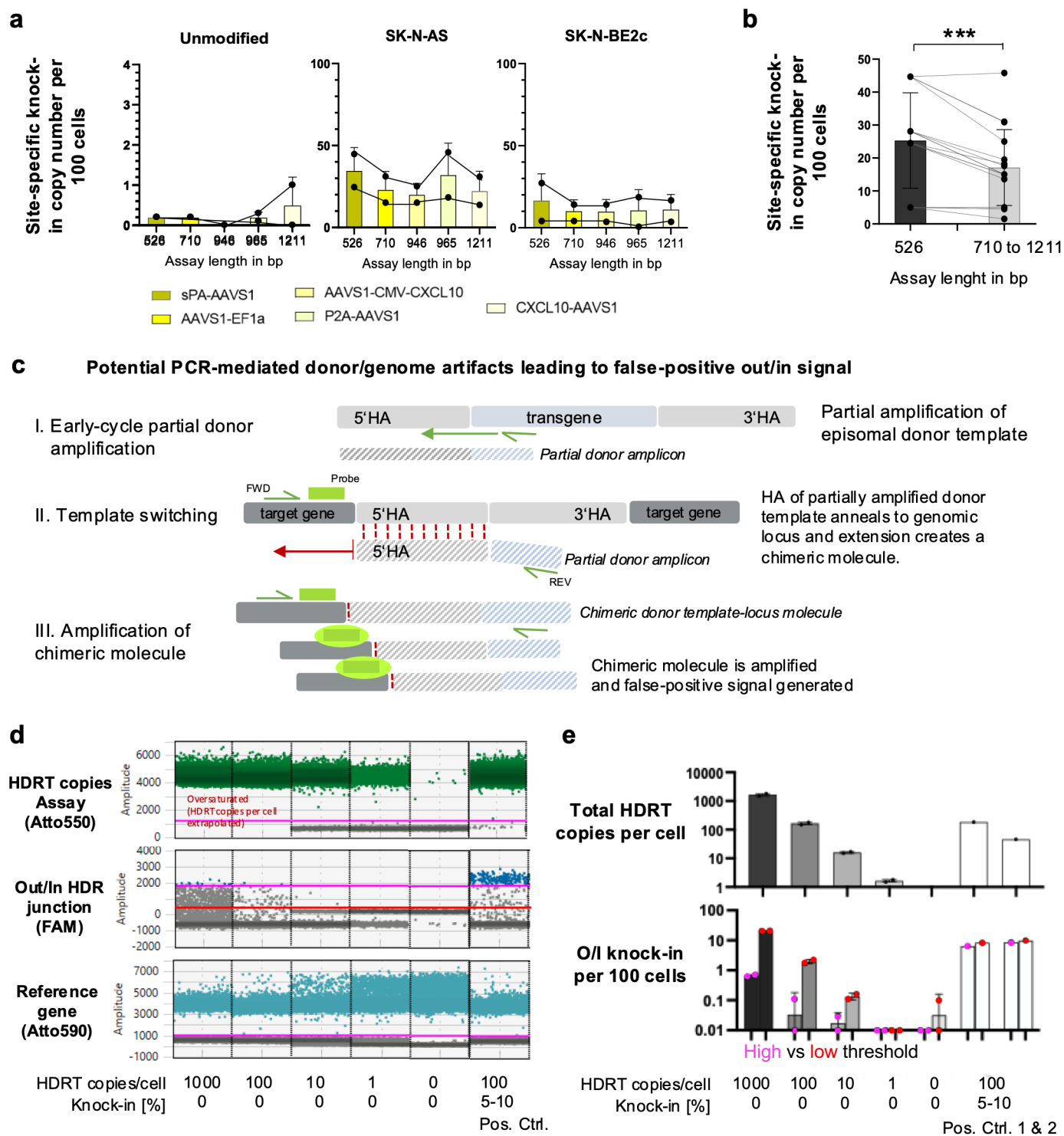

Supplementary Fig. 6. Out/In-In/Out junction assay performance and false-positive risk.

(a) Quantification of site-specific knock-in across Out/In-In/Out assays of increasing amplicon length (526-1211 bp) in unmodified cells and in edited SK-N-AS and SK-N-BE2c cells for multiple donor-locus combinations (sPA-AAVS1, AAVS1-EF1a, AAVS1-CMV-CXCL10, P2A-AAVS1, CXCL10-AAVS1). Longer amplicons consistently reduce apparent knock-in abundance within the same sample and increase variability, indicating decreased amplification efficiency and reduced assay robustness with increasing product length. (b) Paired analysis of knock-in values obtained with the shortest amplicon (526 bp) versus longer assays (710-1211 bp), showing a systematic underestimation of knock-in at longer lengths; statistical comparison demonstrates a significant reduction in measured knock-in for longer amplicons. (c) Schematic illustrating a potential donor/genome template-switching event during PCR. Annealing of the 5' homology arm (HA) of a partially amplified donor during early PCR cycles to the homologous genomic target locus generates a chimeric donor-genome molecule that can subsequently be amplified by the out/in assay and lead to false-positive results. (d) Representative ddPCR plots for the transgene assay (Atto550), out/in assay (FAM), and genomic reference assay (Atto590) across T cell samples with spiked in CD19 CAR HDRT or after CD19 CAR knock-in at the TRAC locus

(positive controls). Mixing increasing amounts of free HDR donor template into untreated cellular gDNA generated a low-amplitude out/in-positive population, representing potential false-positive events, which was either included or excluded from quantification depending on the thresholding strategy applied (inclusive, red; exclusive, pink). **(e)** Quantification of total HDR template copies per cell (top) and out/in-positive events per 100 cells (bottom) for the indicated conditions. Comparison of inclusive and exclusive thresholding demonstrates a low-amplitude out/in-positive population that increases with the amount of free HDR donor template, consistent with potential false-positive signal generation. **Data presentation:** (a,b, e) mean ± SD; dots represent biological replicates. **Statistical tests:** (b) two-tailed paired Wilcoxon test; exact p values are provided in the Source Data file. p values: n.s., not significant; \*\*\*<0.001. Source data are provided as a Source Data file.

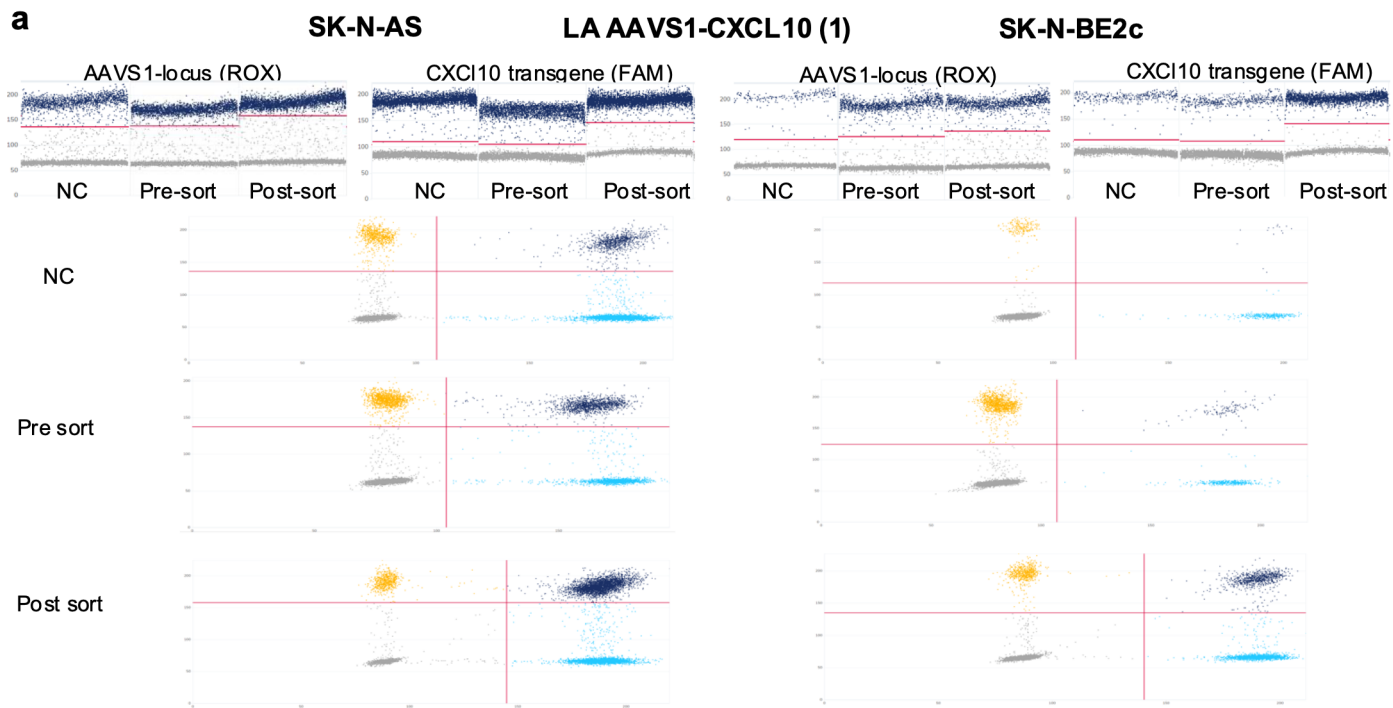

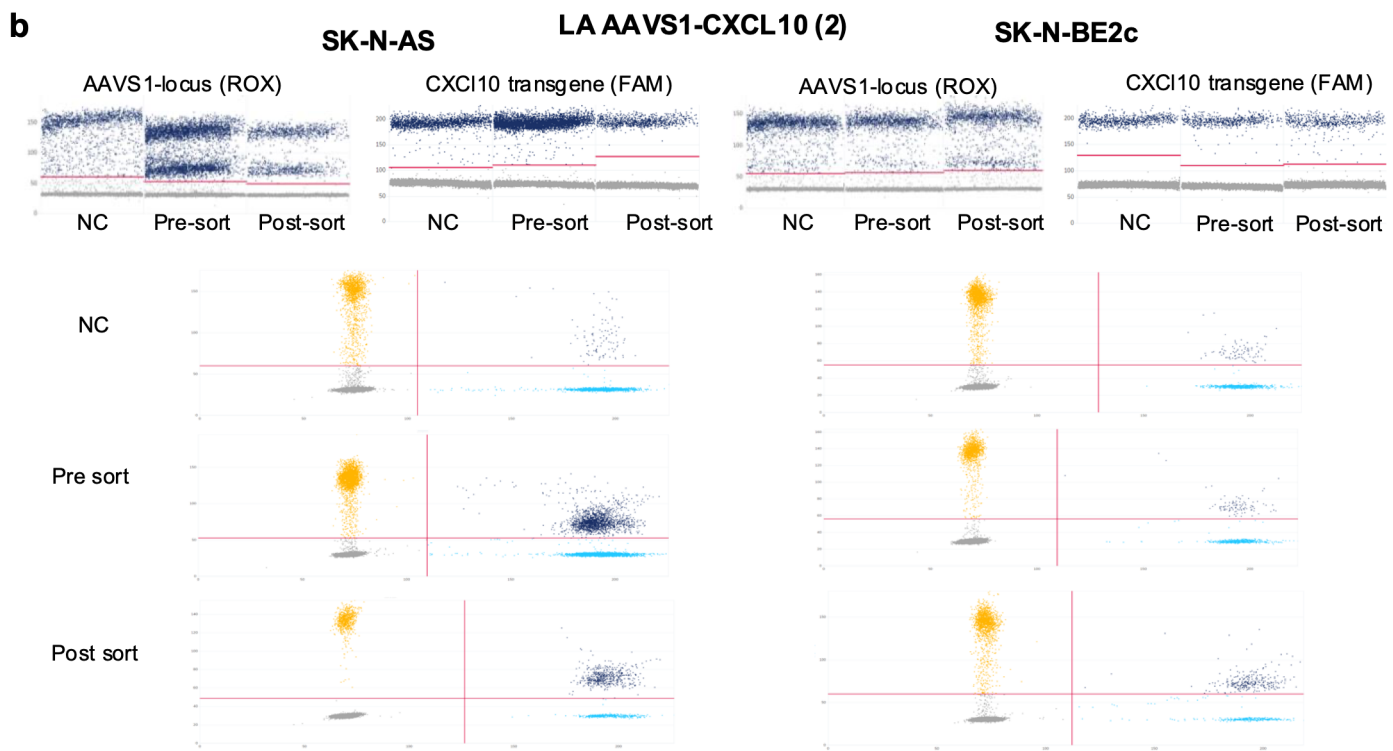

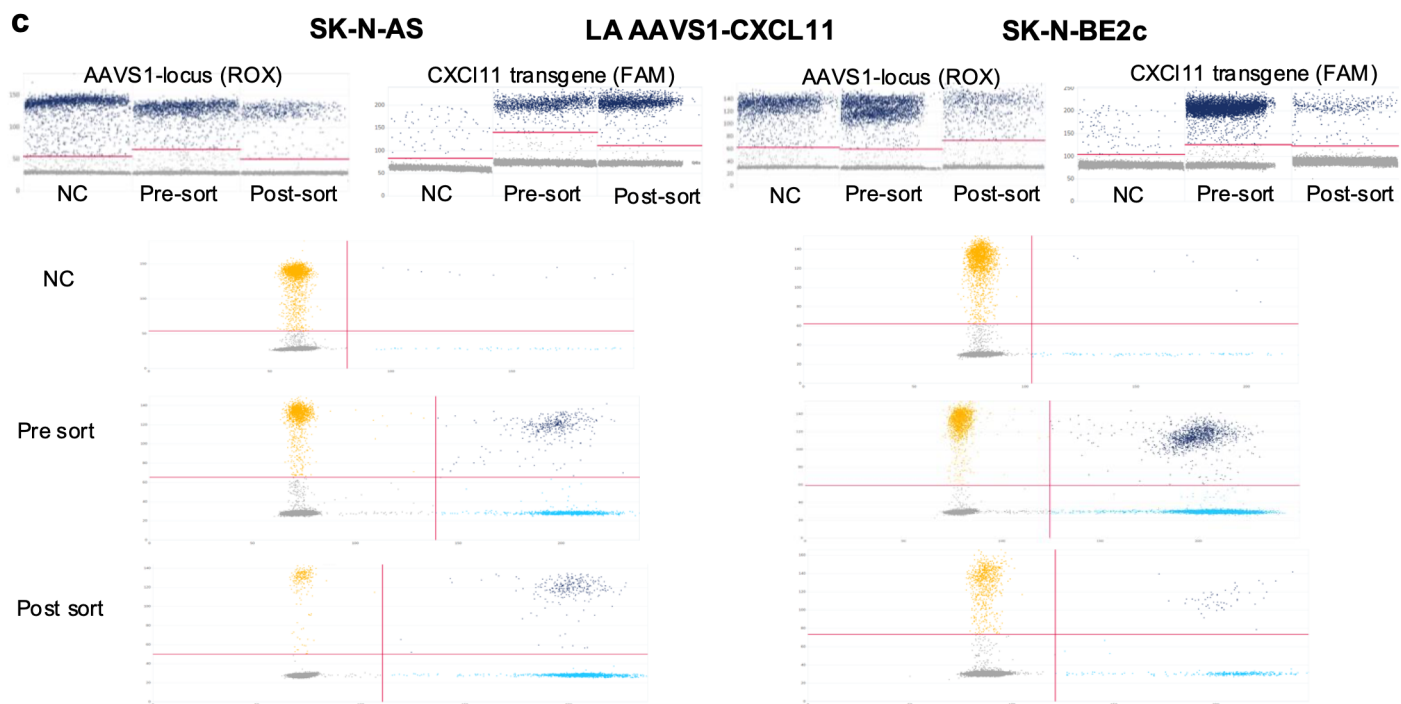

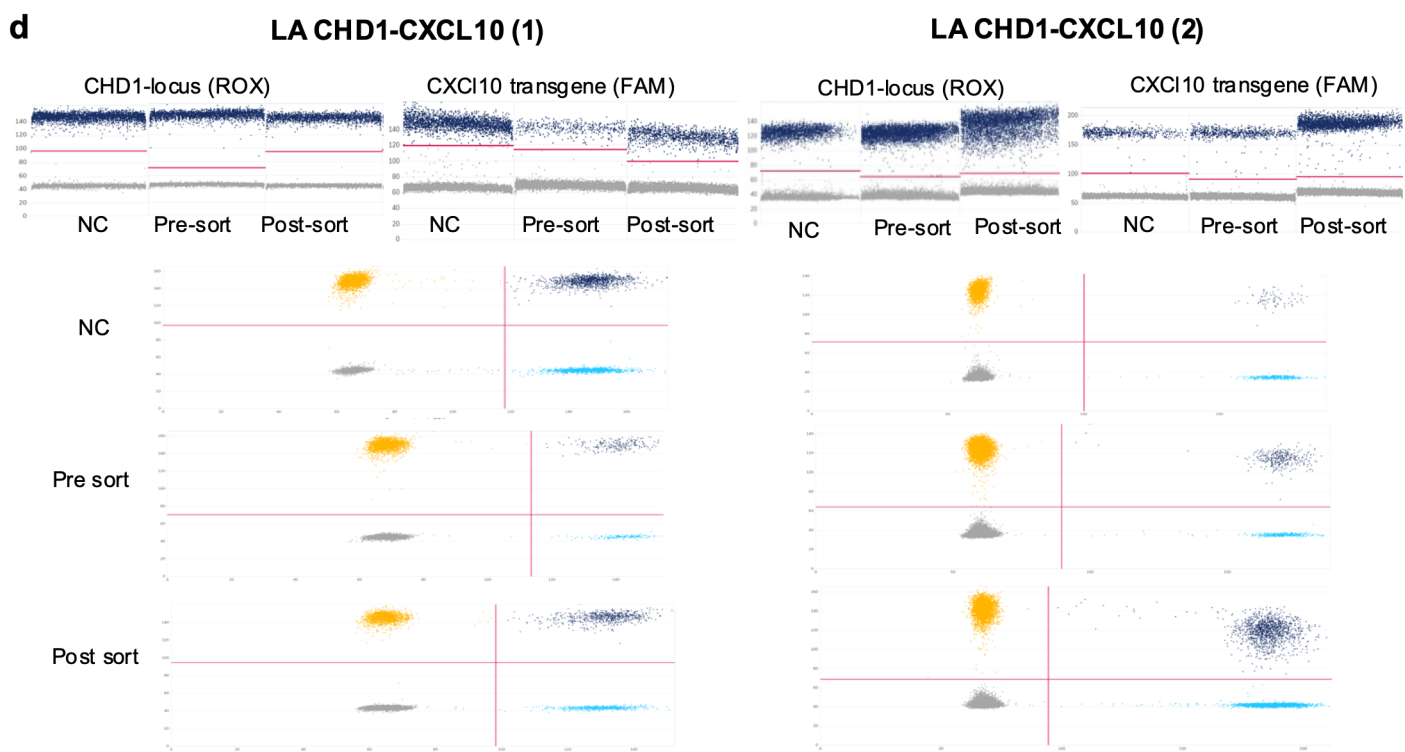

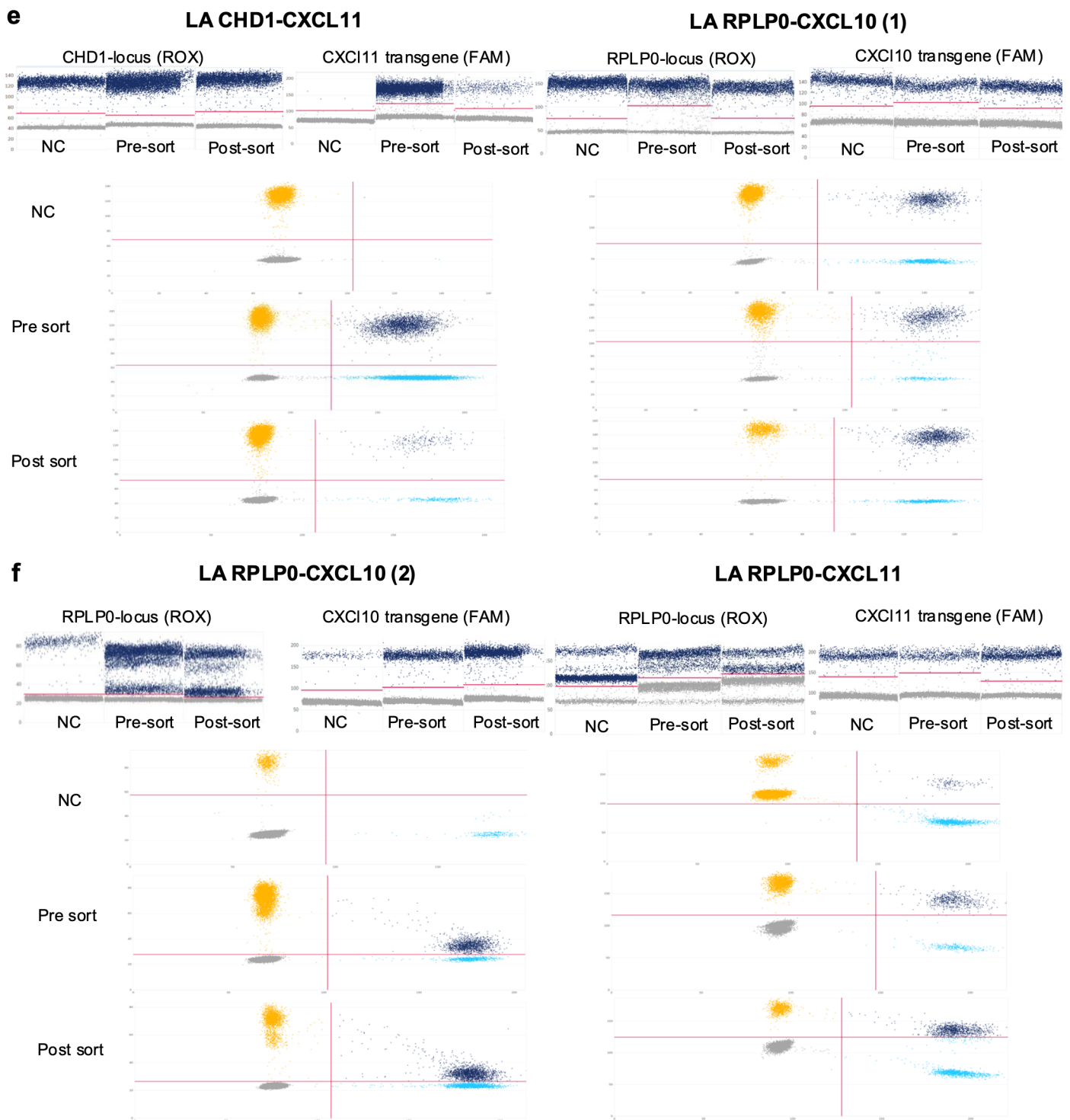

**Supplementary Fig. 7. Linkage assay amplitude plots demonstrate clean cluster separation and robust WT-TG co-positivity for site-specific knock-in.**

**(a)** Representative 1D and 2D amplitude plots for linkage assay variant 1 (AAVS1-CXCL10 (1)) in SK-N-AS and SK-N-BE2c cells. ROX detects the wild-type AAVS1 locus and FAM detects the CXCL10 transgene. Negative controls (NC), pre-sort, and post-sort samples show increasing frequencies of ROX<sup>+</sup>FAM<sup>+</sup> double-positive droplets in edited populations, demonstrating clear on-target HDR-driven linkage. Both cell lines exhibit minimal rain and well-defined cluster boundaries. **(b)** Plots for linkage assay variant 2 (AAVS1-CXCL10 (2)), confirming highly reproducible WT-TG co-positivity patterns across independent assay designs and cell lines. Here, colocalization of the two assay designs within the same partition caused signal interference, generating two distinct positive clouds in the Rox channel (i.e. reduced signal intensity); however, separation from the negative droplet population remained fully preserved. **(c)** Linkage assay performance for AAVS1-CXCL11 donors in SK-N-AS and SK-N-BE2c, demonstrating robust WT-TG co-positivity patterns with minimal rain. **(d)** CHD1-targeting linkage assays with CXCL10 donors (two independent designs) showing expected emergence of linked droplets in edited samples and absence of linkage in NCs. **(e)** Linkage assays for CHD1-CXCL11 and

RPLP0-CXCL10 (variant 1) donors, confirming strong HDR-driven linkage at both loci and clear discrimination between WT-only, transgene-only, and linked droplets. **(f)** Additional RPLP0-targeting linkage assays (CXCL10 (2) and CXCL11) showing robust transgene linkage in edited samples and stable cluster separation across assay designs.

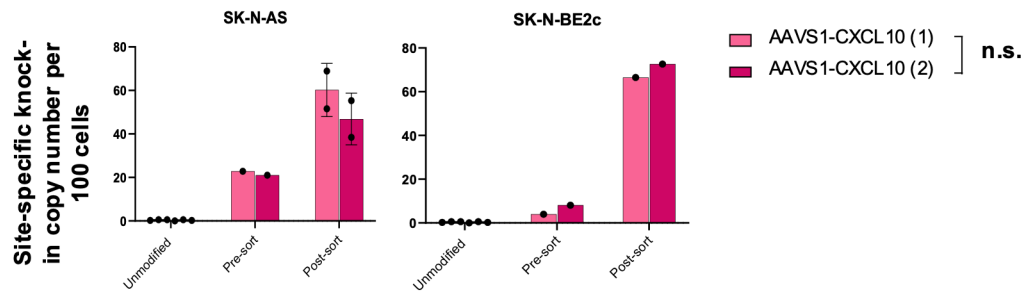

**Supplementary Fig. 8. Concordance of independent linkage assay designs for quantifying AAVS1-CXCL10 knock-in.**

**(a)** Comparison of site-specific knock-in values obtained using two independent linkage assay designs targeting the same AAVS1-CXCL10 donor (AAVS1-CXCL10 (1) and AAVS1-CXCL10 (2)) in SK-N-AS and SK-N-BE2c cells. Both assays yield highly similar knock-in estimates across unmodified, pre-sort, and post-sort samples, demonstrating assay interchangeability and confirming that independent primer-probe configurations do not systematically bias linkage-derived HDR quantification. **Data presentation:** (a) mean ± SD; dots represent biological replicates. **Statistical tests:** (a) two-way ANOVA; exact p values are provided in the Source Data file. n.s., not significant. Source data are provided as a Source Data file.

### SK-N-AS

HA AAVS1-
CXCL10 (1)

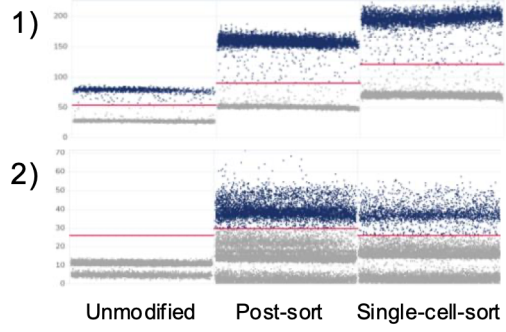

HA AAVS1-
CXCL11

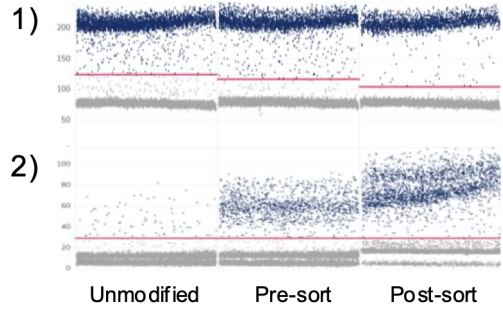

HA RPLP0-
CXCL10 (1)

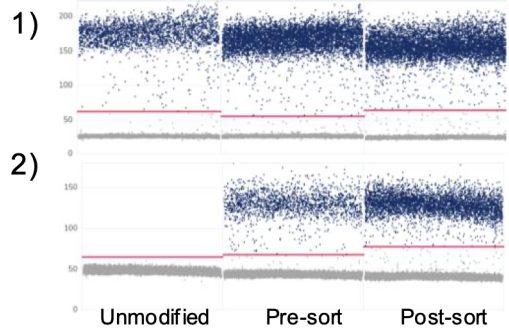

HA RPLP0-
CXCL11

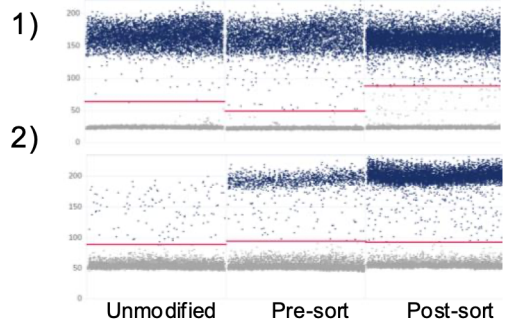

### SK-N-BE2c

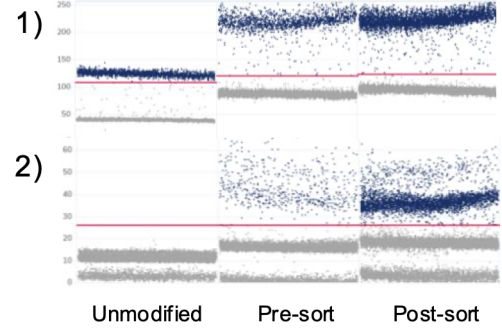

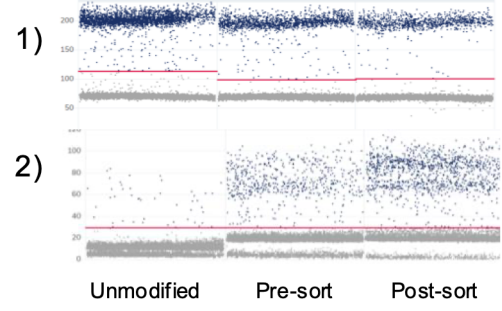

No knock-in at RPLP0 locus in SK-N-BE2c

No knock-in at RPLP0 locus in SK-N-BE2c

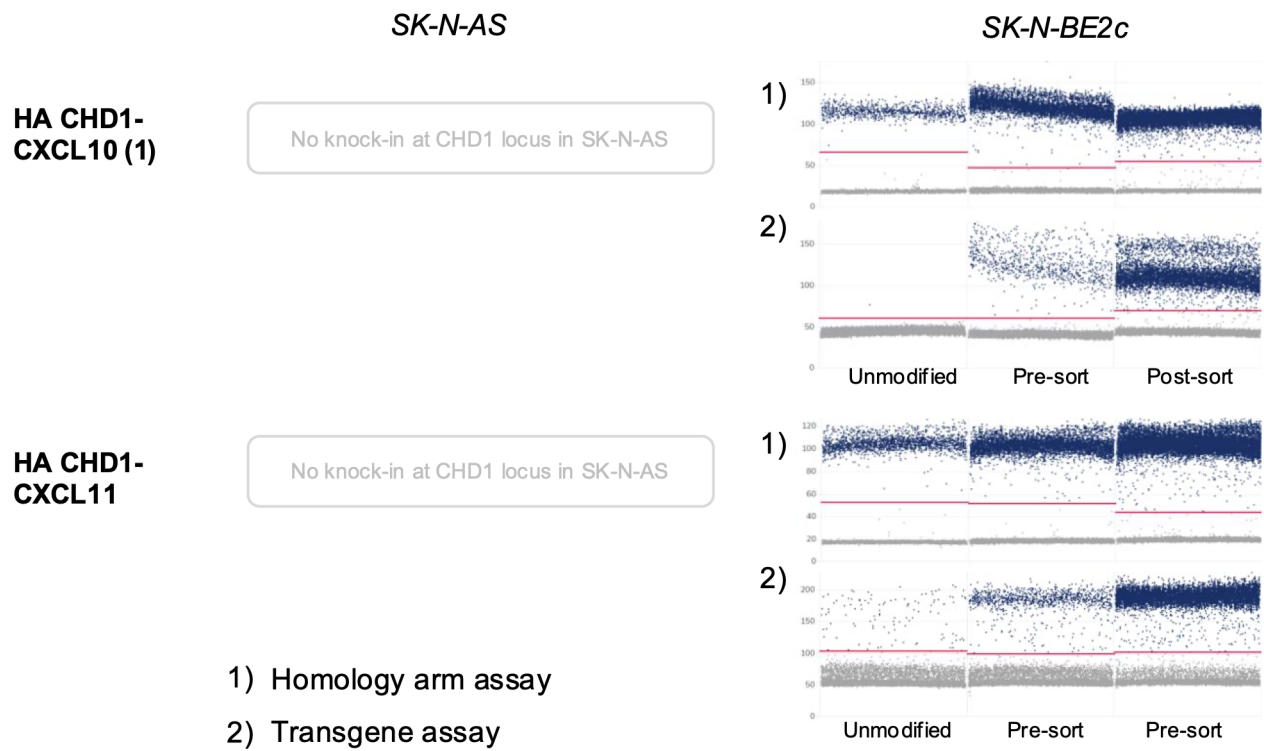

**Supplementary Fig. 9. Homology-arm (HA) and transgene assay amplitude plots demonstrate correct behavior across donors, loci, and edited cell populations.**

Representative 1D amplitude plots for paired homology-arm assays (1) and transgene assays (2) across AAVS1-, RPLP0- and CHD1-targeted CXCL10 and CXCL11 knock-ins in SK-N-AS and SK-N-BE2c cells. HA assays show WT-like distributions in unmodified samples, with clear emergence of HA-positive clusters in edited cells and further enrichment after sorting. Matched transgene assays display concordant patterns, suggesting robust HDR-mediated integration in SK-N-AS and in SK-N-BE2c. RPLP0-targeted edits were analysed only in SK-N-AS, and CHD1-targeted edits only in SK-N-BE2c, where strong HA- and transgene-positive clusters appear in edited and sorted populations. Across all loci and donors, both HA and transgene assays exhibit minimal rain and stable separation from negative droplets.

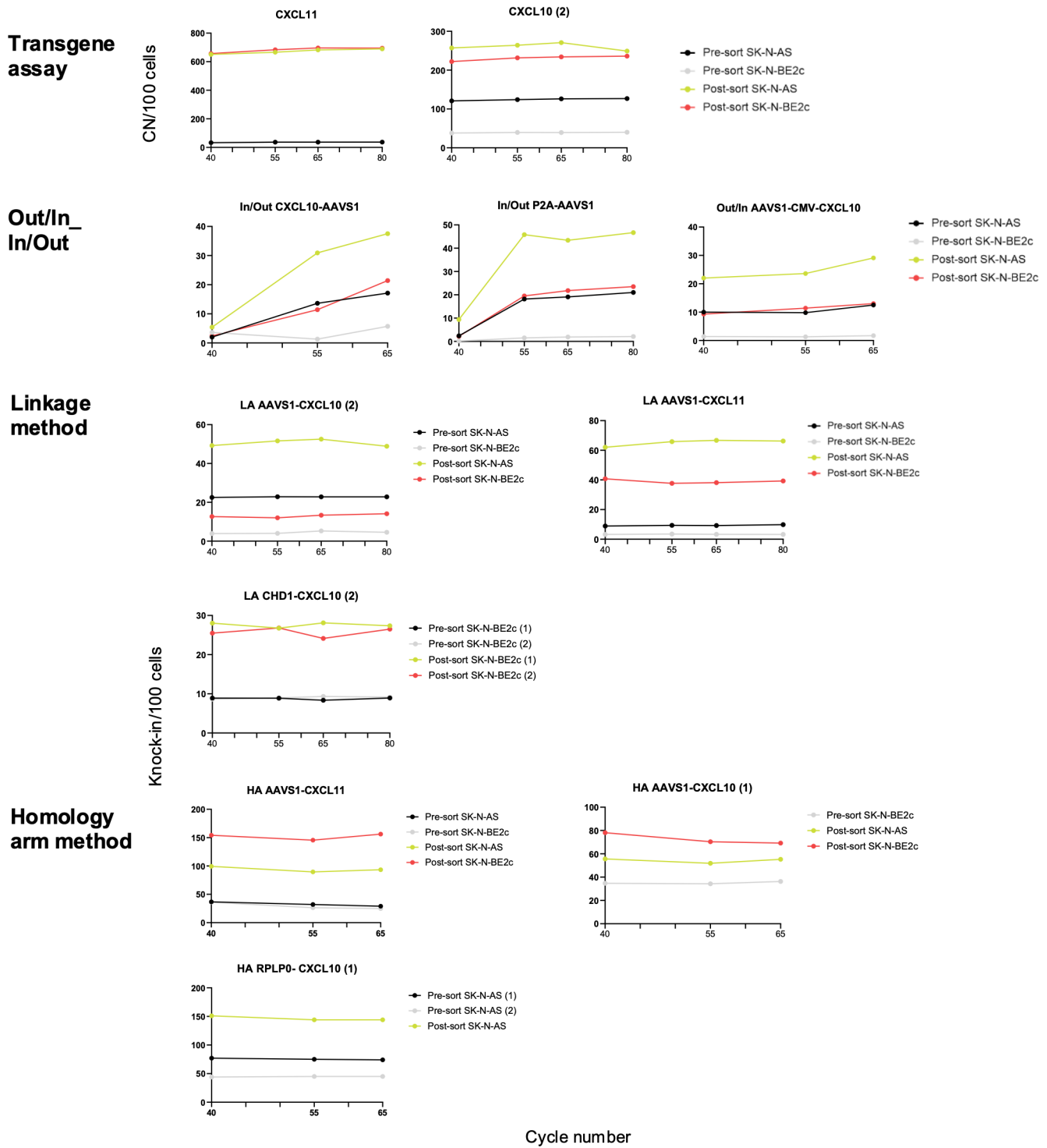

**Supplementary Fig. 10. Effect of cycle number on quantitative stability of transgene, Out/In-In/Out, linkage, and homology-arm assays.**

Knock-in estimates across increasing PCR cycle numbers for transgene assays, Out/In-In/Out assays, linkage assays, and homology-arm assays in SK-N-AS and SK-N-BE2c cells. Transgene, linkage, and HA assays display stable quantitative performance from 40 to 55 cycles with minimal drift across pre-sort and post-sort samples, whereas Out/In-In/Out assays show marked variability and upward bias at higher cycle numbers, consistent with decreased robustness of long-amplicon designs under insufficient cycling counts. Across donors, loci, and assay types, short-amplicon dPCR designs (transgene, linkage, HA) maintain consistent knock-in estimates, while long-range assays require locus-specific optimization to avoid amplification-induced inflation or underestimation. **Data presentation:** mean ± SD; dots represent biological replicates.

#### Linkage assay

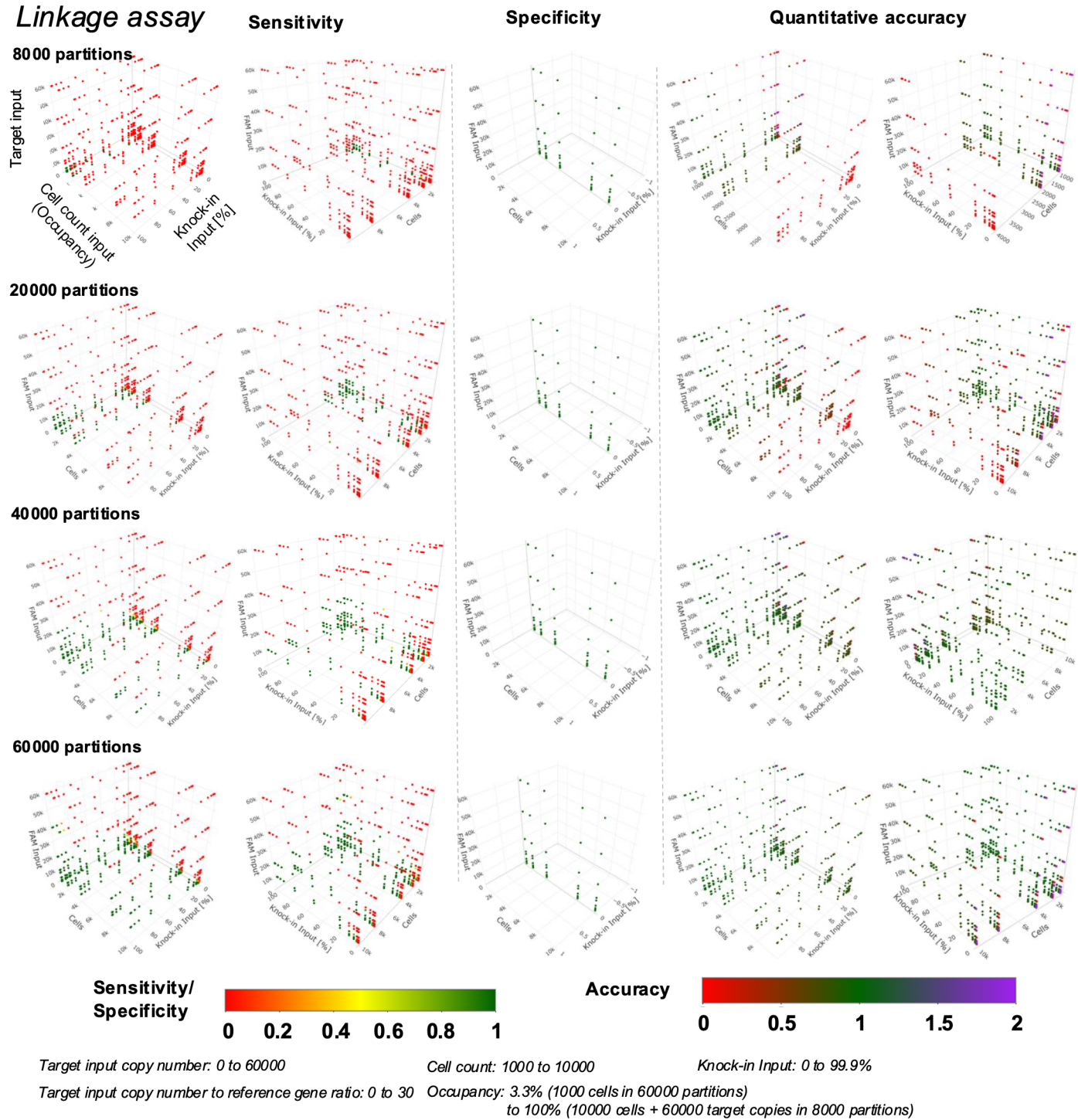

**Supplementary Fig. 11. Monte Carlo simulations reveal the performance landscape of the linkage assay across partition counts, cell inputs, and knock-in frequencies.**

Three-dimensional simulation grids depicting sensitivity, specificity, and quantitative accuracy of the linkage assay across 8000, 20 000, 40 000, and 60 000 partitions. Each simulation varies target-input copy number, knock-in fraction, and cell-equivalent input, spanning occupancy levels from 3.3% (1000 cells in 60 000 partitions) to saturation (>100% occupancy). Sensitivity increases markedly with partition count, especially at low knock-in fractions; specificity remains high across most conditions but decreases at extreme occupancy or very high target-to-reference ratios. Quantitative accuracy improves progressively with partition number and is optimal in intermediate occupancy regimes, with highest accuracy observed at $\geq 40\,000$ partitions. Heatmaps indicate sensitivity/specificity (red-green scale) and accuracy (red-purple scale). **Data presentation:** simulated datasets; 100 independent replicates per parameter combination.

#### Linkage assay

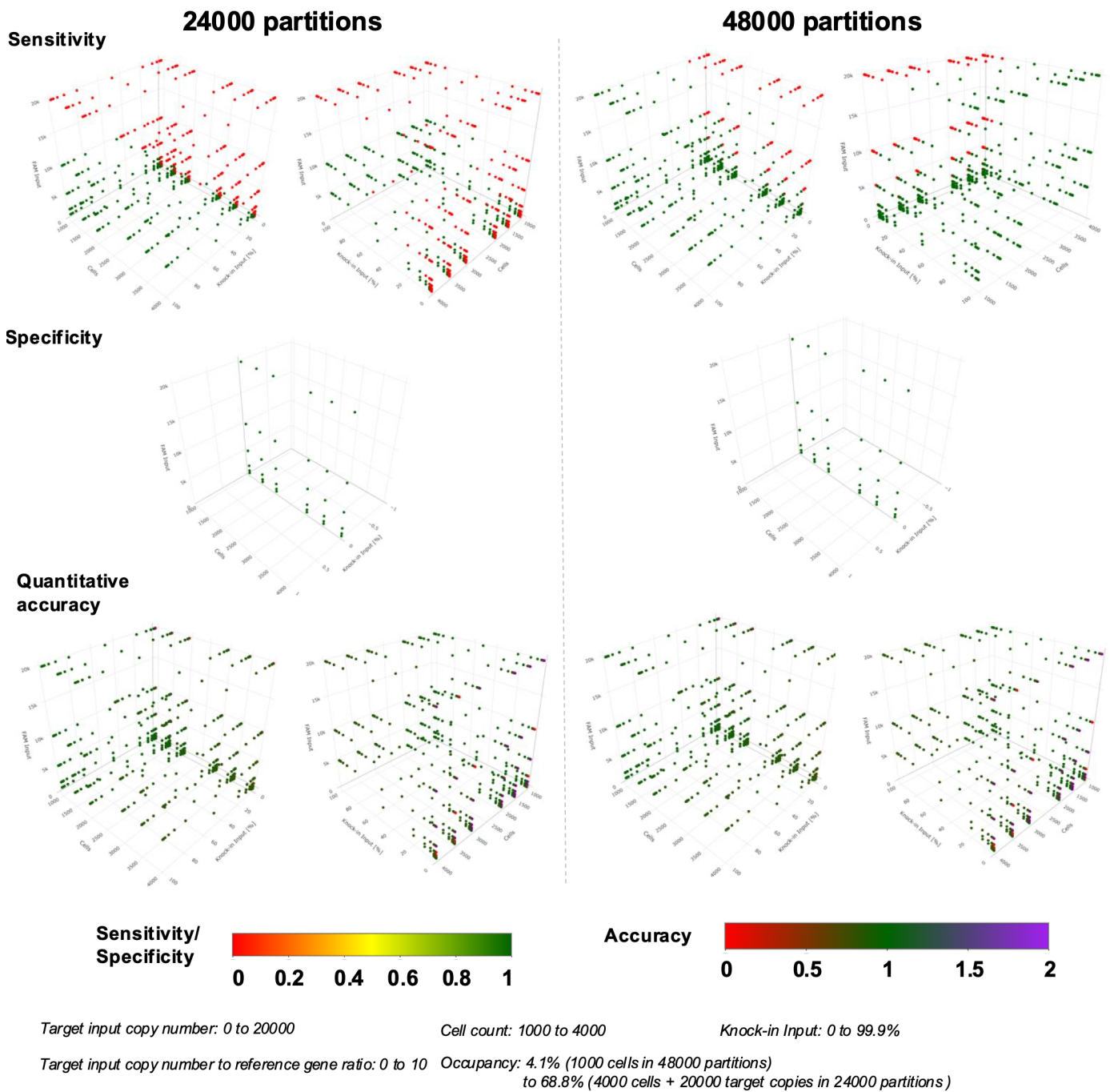

**Supplementary Fig. 12. Impact of intermediate versus high partition numbers on linkage assay performance across simulated editing conditions.**

Three-dimensional simulation grids illustrating sensitivity, specificity, and quantitative accuracy of the linkage assay at 24 000 and 48 000 partitions with realistic input parameters for statistical analysis. Across a parameter space spanning varying target-input copy numbers, knock-in fractions, and cell inputs, sensitivity improves markedly when increasing partitions from 24 000 to 48 000, especially at low knock-in abundance, where false-negative rates decline. Specificity remains uniformly high across both partition counts except at extreme occupancy levels. Quantitative accuracy increases with partition number, with 48 000 partitions producing tighter error distributions and more reliable knock-in estimates across diverse input scenarios. Heatmaps depict sensitivity/specificity (red-green) and accuracy (red-purple) scales. **Data presentation:** simulated datasets; 100 independent replicates per parameter combination.

#### Homology arm based

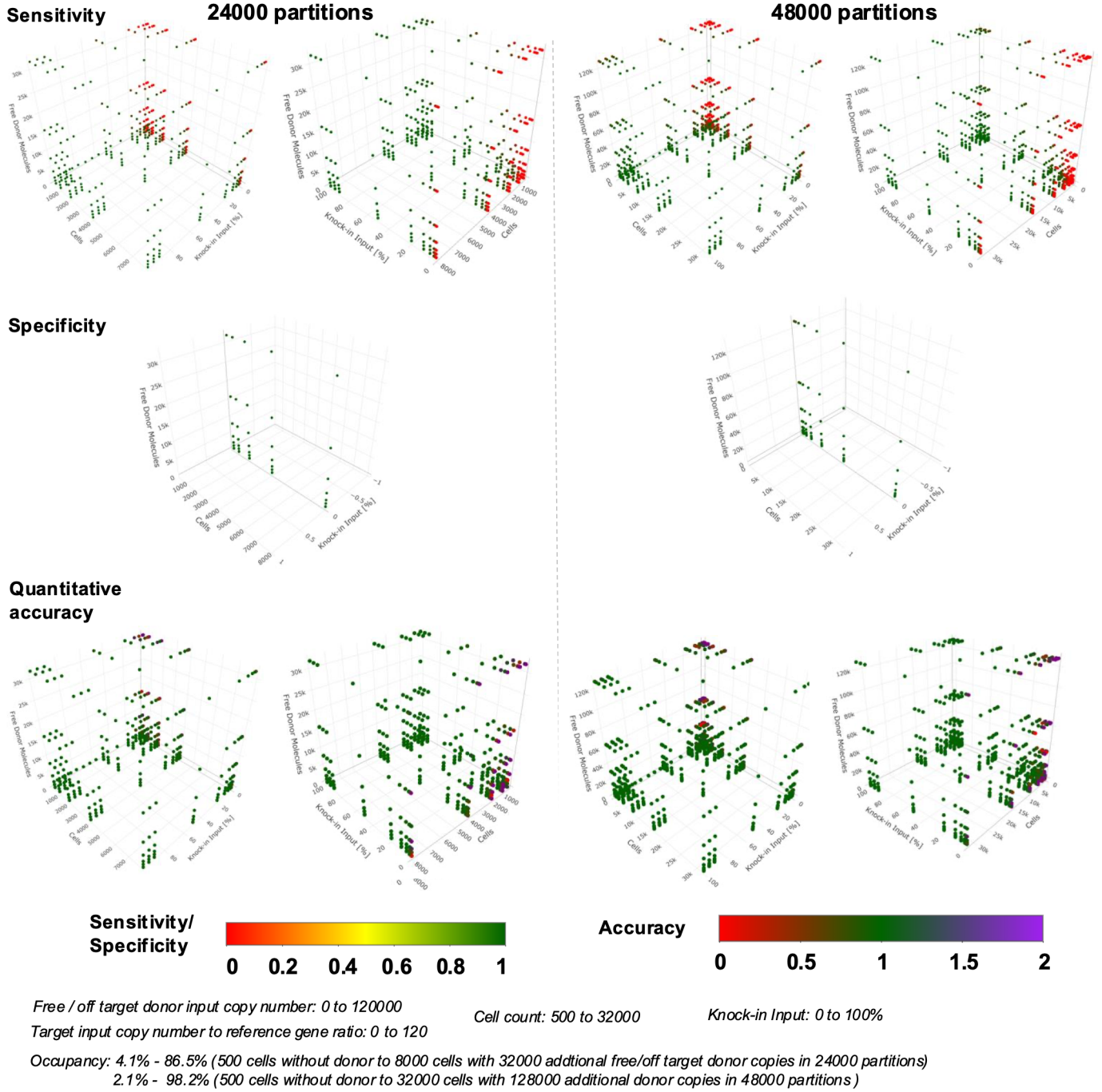

**Supplementary Fig. 13. Monte Carlo simulations reveal robustness of the homology-arm (HA) assay across wide ranges of donor noise, knock-in fractions, and partition counts.**

Three-dimensional simulation grids showing sensitivity, specificity, and quantitative accuracy of the HA assay at 24 000 and 48 000 partitions across varying knock-in inputs, cell counts, and free/off-target donor background. Sensitivity improves with increasing partition number, especially at low knock-in fractions, while specificity remains uniformly high across all tested conditions except at extreme saturation. Quantitative accuracy is maintained across broad ranges of donor noise because HA-based knock-in inference depends on relative channel arithmetic rather than absolute droplet counts, leading to stable performance up to very high occupancy. Accuracy is highest at moderate occupancy and modestly reduced only near saturation (>95%). Heatmaps depict sensitivity/specificity (red-green) and accuracy (red-purple) scales. **Data presentation:** simulated datasets; 100 independent replicates per parameter combination.

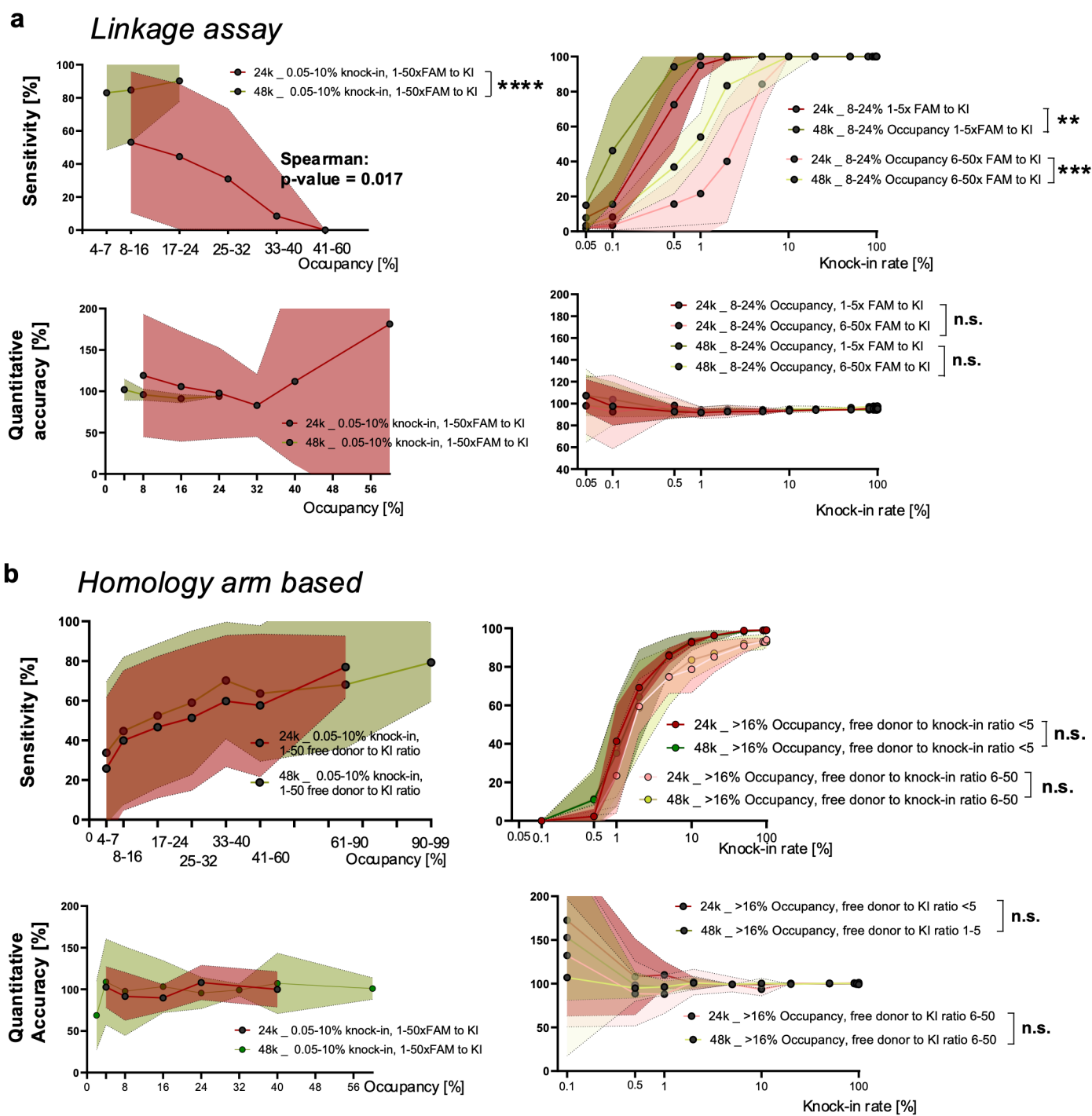

**Supplementary Fig. 14. Relationship between partition occupancy, knock-in abundance, and assay performance for linkage and homology-arm methods.**

**(a)** Performance characteristics of the linkage assay across occupancy levels and knock-in frequencies at 24 000 and 48 000 partitions. Sensitivity decreases significantly with high occupancy at low knock-in rates, with higher partition availability yielding superior performance; Spearman correlation testing confirms an inverse correlation between occupancy and sensitivity. In line quantitative accuracy declines at high occupancy due to increased WT-TG colocalization noise. Sensitivity curves plotted against knock-in rate show improved detection at $\geq 0.5$-1% KI, with 48 000 partitions outperforming 24 000 across all donor-to-reference ratios. Accuracy remains stable across knock-in frequencies except at very low KI, where stochastic variation dominates. **(b)** Equivalent analysis for the homology-arm assay. Here, sensitivity improves with occupancy due to the HA method's dependence on relative rather than absolute molecule counts. Quantitative accuracy is stable across occupancies. Both Sensitivity and quantitative accuracy are largely unaffected by free/off-target donor background within the tested range, with no significant differences between 24 000 and 48 000 partitions. All performance metrics shown in Supplementary Fig. 14 are derived from the simulation datasets presented in Supplementary Figs. 12 and 13. **Data presentation:** (a,b) mean ± SD from simulated datasets; shaded regions represent the full distribution across 100 replicates per condition. **Statistical tests:** (a) Spearman correlation for sensitivity-occupancy relationship; two-way ANOVA for knock-in rate comparisons; (b) two-way ANOVA for partition-count comparisons; n.s., not significant; \*$<0.05$; \*\*$<0.01$. Source data are provided as a Source Data file.

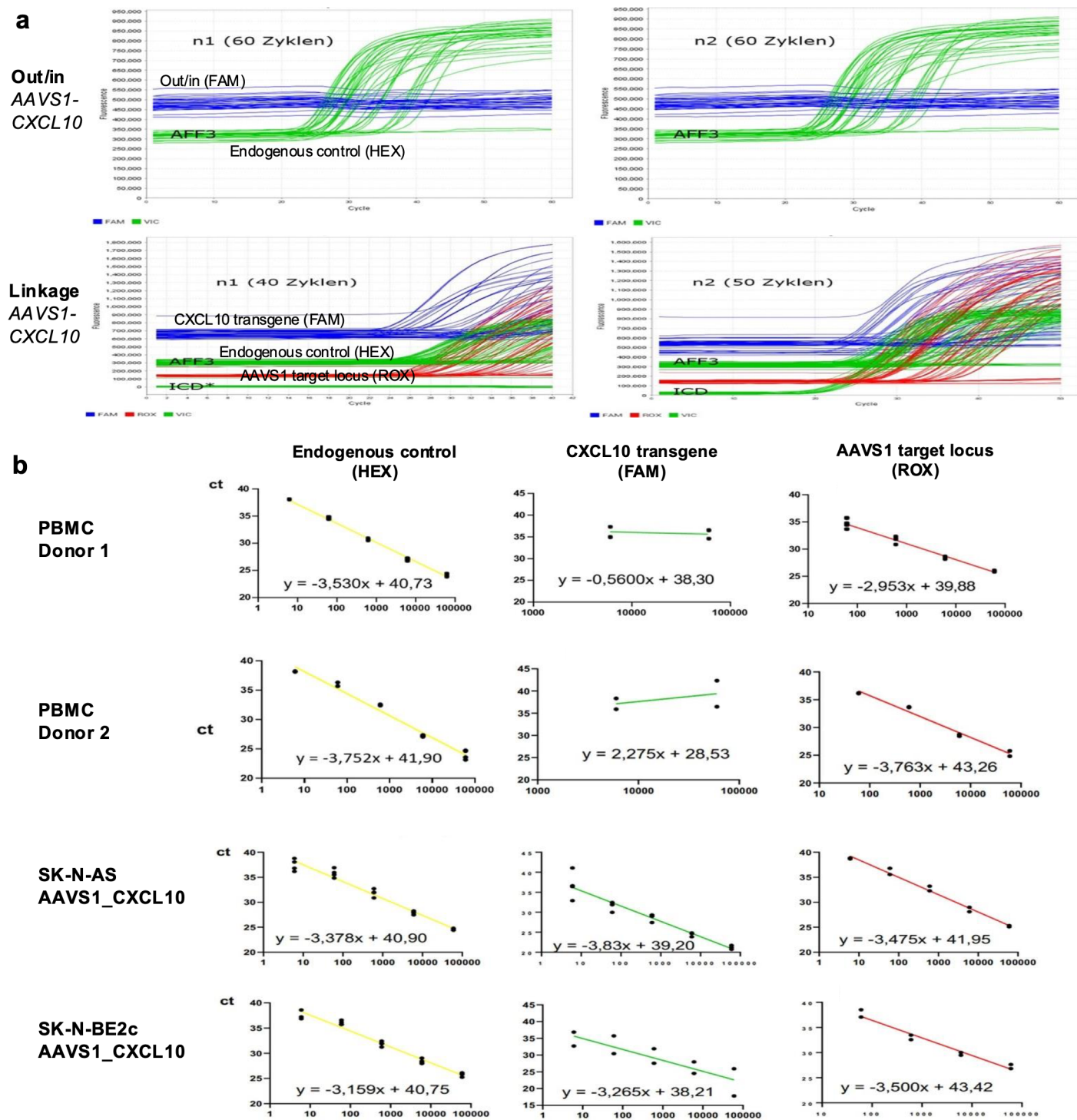

Supplementary Fig. 15. qPCR performance of Out/In-In/Out, linkage, and homology-arm assay components across primary cells and neuroblastoma lines.

(a) qPCR amplification curves for Out/In-In/Out and linkage assay components targeting AAVS1-CXCL10, showing amplification behavior of the FAM-labelled transgene channel, HEX-labelled endogenous control (AFF3), and, for the linkage method, the ROX-labelled AAVS1 target-locus assay. Out/In-In/Out assays do not show exponential amplification and a flat signal, whereas the individual short-amplicon components used in the linkage assay exhibit consistent exponential amplification with earlier Ct values. (b) Standard curves and linear regressions for HEX (endogenous control), FAM (CXCL10 transgene), and ROX (AAVS1 target locus) across PBMCs from two donors and edited SK-N-AS and SK-N-BE2c cells. Short-amplicon linkage components yield linear, well-behaved qPCR responses with slopes approximating expected efficiencies underscoring the robustness of linkage and HA assay components for potential qPCR use.

#### a Dilutions

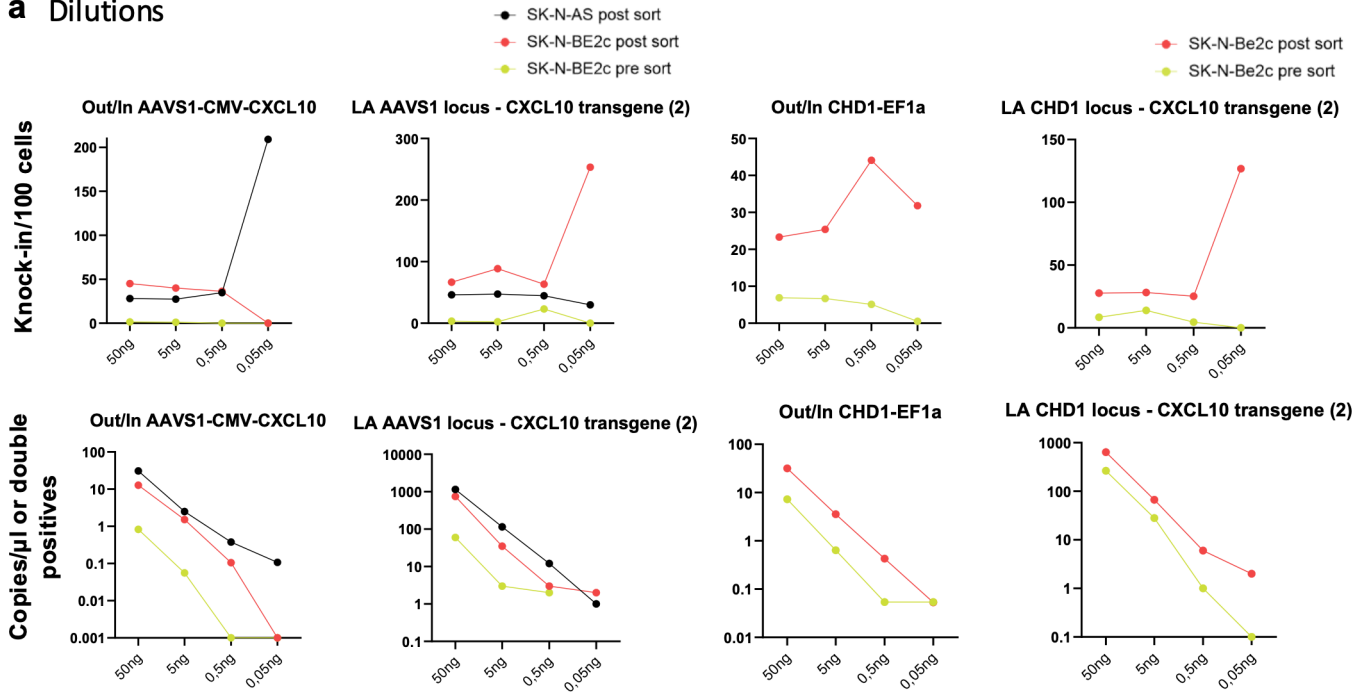

## b

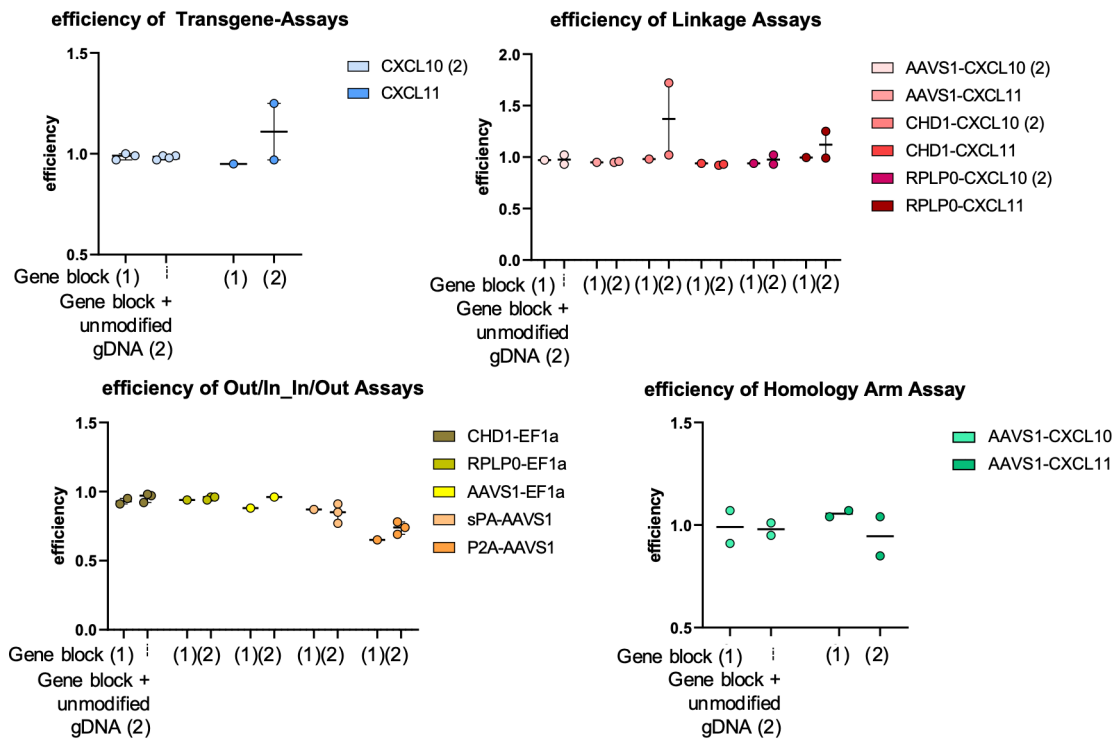

**Supplementary Fig. 16. Dilution-series performance and analytical efficiency of transgene, linkage, Out/In-In/Out, and homology-arm assays.**

**(a)** Dilution-series analysis (50 ng, 5 ng, 0.5 ng, 0.05 ng gDNA) for Out/In and linkage assays across SK-N-AS and SK-N-BE2c samples. Out/In assays show strong dilution-dependent instability, with knock-in estimates becoming highly variable or collapsing at lower inputs. Linkage assays display a more proportional decrease in transgene or double-positive counts across dilutions, maintaining monotonic signal decay in most samples, although both assay types show signal loss at the lowest DNA amounts. **(b)** Analytical efficiency of transgene, linkage, Out/In-In/Out, and homology-arm assays measured using gene-block DNA alone and gene-block DNA spiked into unmodified genomic DNA. Transgene and linkage assays show efficiencies near unity across donors and loci. Out/In-In/Out assays display greater variance and primer-probe combination-dependent efficiency loss, reflecting long-amplicon effects. Despite visible variance across donors, homology-arm assays retain mean efficiencies near unity under both gene-block and mixed gDNA conditions, consistent with expectations for short-amplicon assays. **Data presentation:** (a,b) mean ± SD; dots represent biological replicates.

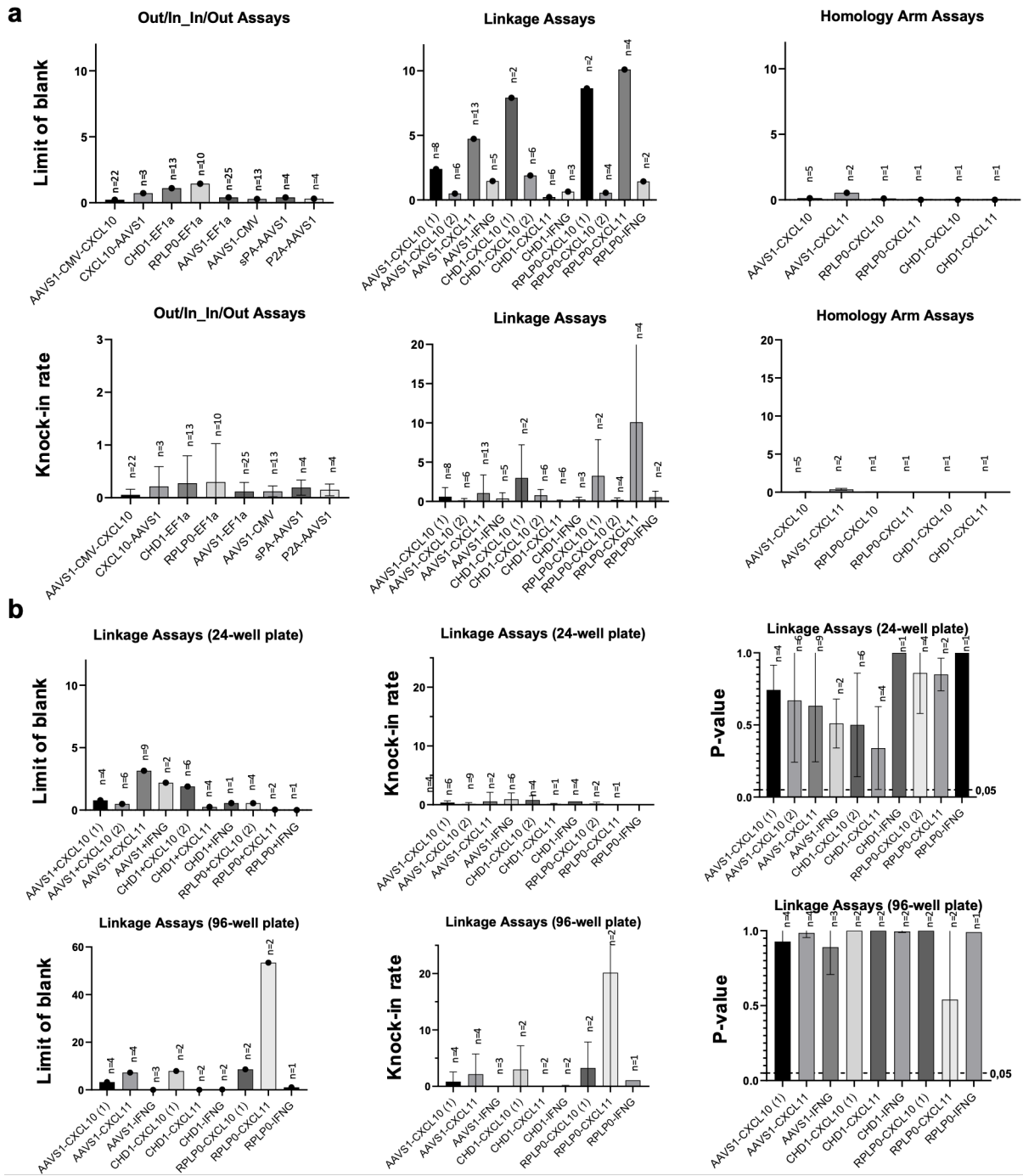

**Supplementary Fig. 17. Limit-of-blank characteristics and background knock-in estimates for Out/In-In/Out, linkage, and homology-arm assays across donor constructs and plate formats.**

**(a)** Limit-of-blank (LoB) values and corresponding knock-in background rates for Out/In-In/Out, linkage, and homology-arm assays across all AAVS1- and CHD1-targeting donor templates, extending the LoB analysis from Fig. 3e by additionally reporting assay-intrinsic background knock-in frequencies, which were not shown in the main figure; Linkage assays exhibit the highest variability, reflecting stochastic WT-TG colocalization, and homology-arm assays yield the lowest and most stable LoB. **(b)** Effect of partition number on linkage assay LoB, background knock-in, and p-value-based specificity in 24-well versus 96-well formats, providing an expanded view beyond Fig. 3f, which reports only 24-well specificity and does not include 96-well data; higher partition counts reduce LoB and improve negative-control discrimination, whereas lower partition counts increase stochastic co-positive events, inflate LoB, but shifts p-values away from the significance threshold. **Data presentation:** mean ± SD; dots represent replicate measurements, with n values shown above bars. **Statistical analysis:** LoB calculated as the 95th percentile of negative-control replicates according to CLSI EP17-A2; background knock-in derived from Poisson-inverted copy numbers; p-values computed using binomial tests comparing observed and expected double-positive counts under the null hypothesis of random colocalization; Source data are provided as a Source Data file.

**Supplementary Fig. 18. Intra- and inter-assay precision comparison between Out/In-In/Out and linkage assays across donor constructs and loci.**

**(a)** Intra-assay coefficient of variance (CoV) for Out/In-In/Out and linkage assays across AAVS1-CMV-CXCL10, AAVS1-EF1α, CHD1-EF1α, and RPLP0-EF1α donor-locus combinations, showing that linkage assays generally exhibit lower variability within runs, although differences between assay types are not statistically significant. **(b)** Inter-assay CoV across independent runs for the same donor-locus combinations, demonstrating reproducibility patterns similar to (a), with linkage assays tending toward lower between-run dispersion than Out/In-In/Out assays, again without statistically significant differences. **Data presentation:** bar graphs with overlaid individual data points and whiskers representing SD; n values correspond to independent biological or technical replicates. **Statistical analysis:** CoV defined as $100 \times (\text{SD}/\text{mean})$ for replicate measurements; comparisons between assay types performed using two-tailed Two-Way-Anova, with non-significant differences indicated (n.s.); Source data are provided as a Source Data file.

Supplementary Fig. 19. Flow cytometric analysis of transgene expression

(a) Gating strategy for detecting reporter-positive cells using PE-conjugated CD34 antibody (Qbend) for the Q8 epitope on unmodified and CRISPR edited neuroblastoma cell lines. (b) Gating strategy for detecting CD3 knock-out and anti-CD19 CAR positive T cells.

### Multiplexing

#### a Multiplexing MMs

|  | Mastermix 1 | Mastermix 2 | Mastermix 3 | Mastermix 4 |
| --- | --- | --- | --- | --- |
| Transgene | CXCL10 (1) | CXCL10 (1) | CXCL11 | CXCL11 |
| Reference | AFF3 | AFF3 | AFF3 | AFF3 |
| Out/In | AAVS1-EF1α | sPA-AAVS1 | AAVS1-EF1α | sPA-AAVS1 |
| Cut site | AAVS1 cut site | AAVS1 cut site | AAVS1 cut site | AAVS1 cut site |
| Target locus | AAVS1 locus | AAVS1 locus | AAVS1 locus | AAVS1 locus |

#### b cross talk matrix plate layout

|  | 1 | 2 | 3 |
| --- | --- | --- | --- |
| A | CXCL10 (2)-<br>ATTO550 | Endogenous<br>ctrl-HEX | reference<br>dye |
| B |  |  |  |
| C |  | AAVS1-EF1α-FAM |  |
| D | CXCL11-ATTO550 |  |  |
| E |  | AAVS1<br>locus-ROX |  |
| F |  |  |  |
| G | sPA-AAVS1-FAM | AAVS1 cut<br>site-Cy5 |  |
| H |  |  |  |

#### c Cross talk matrix single stains

Supplementary Fig. 20. Assay composition and cross-talk calibration for five-color multiplex dPCR.

(a) Overview of the four mastermix compositions used during multiplex assay development (Fig. 5). Each mastermix contains one HDR donor assay (CXCL10 or CXCL11), an endogenous reference assay, the AAVS1 target-locus assay, and the AAVS1 cut-site assay, with combinations selected to evaluate signal separation and dye compatibility across fluorophores. (b) Plate layout used to generate a five-color cross-talk matrix on the QIAcuity system. Each well contains a *single* assay labeled with a unique dye (ATTO550, HEX, FAM, ROX, Cy5), enabling quantification of fluorescence bleed-through into other channels and generation of a correction matrix later applied to multiplex reactions. (c) Representative single-stain amplitude plots for all included assays. These plots illustrate primary signal distributions and measurable bleed-through patterns across the unused channels, forming the basis for constructing the cross-talk correction matrix used in Fig. 5 and Supplementary Fig. 21.

#### Multiplexing raw data before and after cross talk matrix

**Supplementary Fig. 21. Effect of cross-talk correction on multiplexed five-color digital PCR amplitude plots.**

Raw two-dimensional fluorescence amplitude distributions for all pairwise channel combinations included in the AAVS1 five-color multiplex assay, shown **before** and **after** application of the QIAcuity cross-talk correction matrix. For each dye-assay combination (e.g., ATTO550-transgene, HEX-endogenous control, FAM-Out/In, ROX-AAVS1 locus, Cy5-cut site), multiplexed reactions exhibit varying degrees of spectral bleed-through into non-target channels, visible as diffuse low-amplitude secondary clouds in the uncorrected plots. Application of the cross-talk matrix markedly improves cluster separation by reducing bleed-through and restoring discrete positive/negative droplet populations. Shown are representative CXCL10 and CXCL11 multiplex reactions structured according to the mastermixes in Supplementary Fig. 20; arrows indicate the direction of correction (left = raw, right = corrected).

#### Multiplexing raw data before and after cross talk matrix (unmodified sample)

**Supplementary Fig. 22. Cross-talk correction improves multiplex cluster separation in unmodified samples.**

Two-dimensional fluorescence amplitude plots for all pairwise channel combinations in the five-color AAVS1 multiplex assay, shown before and after application of the cross-talk correction matrix for an unmodified genomic DNA samples. In the raw multiplex data, unmodified samples display characteristic diffuse low-amplitude bleed-through clouds originating from ATTO550 (CXCL10 transgene assay), HEX (endogenous control), FAM (Out/In assay), ROX (AAVS1 locus), and Cy5 (cut-site assay). Although no true positive populations are expected in unmodified samples for transgene assay or Out/In assay, spectral spillover produces structured background patterns that could interfere with threshold placement or droplet classification in multiplex mode. Application of the cross-talk matrix substantially reduces this bleed-through, resulting in cleaner negative clouds and improved separation between noise and the assay-specific positive droplets observed only in edited samples (see Supplementary Fig. 21). However, despite correction, the Out/In assay (FAM channel) still exhibits sporadic unexpected positive partitions, indicating residual non-specific activation that is intrinsic to the long-range Out/In assay in the multiplex setting rather than a cross-talk artefact. Arrows indicate the direction of correction (left = raw, right = cross-talk-corrected).

a

SPRI (Solid-phase reversible immobilization) bead depletion of free HDR donor template using a low bead:DNA ratio

b

Supplementary Fig. 23. SPRI bead-based depletion of free HDR donor template.

(a) Schematic of the SPRI bead purification strategy used to reduce excess free HDR donor template (HDRT) while retaining genomic DNA. A low bead:DNA ratio was applied to preferentially bind high-molecular-weight genomic DNA, allowing unbound free donor template to remain in the supernatant and be removed during magnetic separation. Genomic DNA was subsequently recovered from the bead fraction for downstream analysis. (b) Ratio of HDR donor template copies to endogenous TRAC locus copies following SPRI purification across a range of bead:DNA ratios in SK-N-AS genomic DNA spiked with HDR donor template at approximately 50 or 150 copies per genome equivalent. HDRT:TRAC ratios indicate preferential depletion of free HDR donor template relative to genomic DNA for SPRI bead:DNA ratios between 0.3-0.45.

#### Supplementary Tables

**Supplementary Table 1. Collection of all Assays**

| Name | Probe sequence | FWD sequence | REV sequence | Amplicon length | Reporter | Quencher |
| --- | --- | --- | --- | --- | --- | --- |
| **Cut Site Assays** |  |  |  |  |  |  |
| AAVS1_wt | CGGTTAATGTGGCTCTGGTT | CTTCAGGACAGCATGTTTGC | ACAGGAGGTGGGGGTTAGAC | 69 | FAM | BHQ-1 |
| AAVS1_cut site | TGTCACCAATCCTGTCCCTAGTG | GCTCTGGTTCTGGGTACTTT | AGGAGGAGGCCTAAGGATG | 100 | Cy5 | BHQ-2 |
| AAVS1_cut site LNA | CC A+AT +C+C+T +GTC +CC | GCTCTGGTTCTGGGTACTTT | AGGAGGAGGCCTAAGGATG | 100 | Cy5 | 3IABkFQ |
| TRAC KO | TCTGCCTATTCACCGATT | CTCTCTTTTGAAGCAGGCC | AGTGTCTGTTTTGACACGATCTGTA CTC | 551 | HEX | 3IABkFQ |
| **Transgene Assays** |  |  |  |  |  |  |
| CXCL10 transgene (1) | CGAGATCATCCCTGCTTCTC | CCTGATCTTCCTGACCCTGA | GTGGCGATGATTTCCACTCT | 165 | FAM | BHQ-1 |
|  | CGAGATCATCCCTGCTTCTC | “ | “ | 165 | ATTO550 | BHQ-2 |
| CXCL10 transgene (2) | CTGCTTCTCAGTTCTGCCCTAGAGTG G | GGAAAAGCTCGAGATCATCC | TCTTTTTCATTGTGGCGATGATT | 71 | FAM | BHQ-1 |
| CXCL11 transgene | ACGCTGTCTTTGCATAGGCCC | CTACAGTTGTTCAAGGCTTC | GGTACATTATGGAGGCTTTC | 111 | FAM | BHQ-1 |
|  | ACGCTGTCTTTGCATAGGCCC | “ | “ | 111 | ATTO550 | BHQ-2 |
| IFNG transgene | TGCAATCCCAGATCGTGAGCTTCT | AAGGAAGAAAGCGATCGAAA | ATAGATTGGTCGTCCTTGAA | 93 | FAM | BHQ-1 |
| P2A-CD19CAR | CTCTCTTTTGAAGCAGGCC | CCTATTCACCGCCACCAAC | GGGCCCAGGATTTTCTTCCACAT | 67 | Cy5 | 3IAbRQSp |
| **Out/In assays** |  |  |  |  |  |  |
| AAVS1-CMV-CXCL10 | CAGGCAGCAGATCAGGATGGCG | GGCATCTCTCCTCCCTCA | GGCATCTCTCCTCCCTCA | 946 | FAM | BHQ-1 |
| CHD1-EF1α | CAGTTTACCCCGCGCCACCT | AGAGACAAGGTCTCACTATG | GCACTTATATACGGTTCTCC | 724 | FAM | BHQ-1 |
| RPLP0-EF1α | CTATTGGCTGCGCCATCGCCCGT | TATCCAATGGTTGCCTGTAT | GCACTTATATACGGTTCTCC | 638 | FAM | BHQ-1 |
| AAVS1-EF1α | TCCCTTGCGTCCCGCCTCCCCTT | GCCGTCTTCACTCGCTG | GCACTTATATACGGTTCTCC | 710 | FAM | BHQ-1 |
| AAVS1-CMV | TGGAGTTCCGCGTTACATAA | CCTTCTTGTAGGCCTGCATC | GGGCCATTTACCGTAAG | 633 | FAM | BHQ-1 |
|  | TCCCTTGCGTCCCGCCTCCCCTT | GCCGTCTTCACTCGCTG | GGGCCATTTACCGTAAG | 633 | FAM | BHQ-1 |
| TRAC KI | CTCTCTTTTGAAGCAGGCC | CTTTGCTGGGCCTTTTTCCC | TACACCTTCTTTTAGGACCCGGG | 528 | Cy5 | 3IAbRQSp |
| **In/Out assays** |  |  |  |  |  |  |
| CXCL10-AAVS1 | CTCTTCACCTTTCTAGTCCCC | CCCTAGAGTGGAAATCATCG | CACCATTCTCACAAAGGGAG | 1211 | FAM | BHQ-1 |
| sPA-AAVS1 | CTCTTCACCTTTCTAGTCCCC | AATAAAAGATCTTTATTTTCATT AGATCTGTGTGTGG | GGCCTTATCTCACAGGTAAA | 526 | FAM | BHQ-1 |
|  | CTCTTCACCTTTCTAGTCCCC | “ | “ | 526 | ROX | BHQ-2 |
| P2A-AAVS1 | CTCTTCACCTTTCTAGTCCCC | CGTGGAAGAAAATCCAGGACC TATG | GGCCTTATCTCACAGGTAAA | 965 | FAM | BHQ-1 |
|  | CTCTTCACCTTTCTAGTCCCC | “ | “ | 965 | ROX | BHQ-2 |
| **Linkage target locus assays** |  |  |  |  |  |  |
| AAVS1 3'wt-link | CTCTTCACCTTTCTAGTCCCC | AAGTTGGTCCTGAGTTCTAA | GGCCTTATCTCACAGGTAAA | 95 | ROX | BHQ-2 |
| CHD1 5'wt-link | TGGCCTCCCAGTCATGCTGG | CTCAAGTTATCTTCCTGCCT | AAGAAAAATGTGGCTGAGTG | 82 | ROX | BHQ-2 |
| RPLP0 5'wt-link | GCCAACGGAAAGCCGCCTT | CTTCGCGACCCTACTTA | CAAAGCTGAACGAATACAGG | 89 | ROX | BHQ-2 |
| TRAC 5'wt | CCAGCAATATAACTCTGG | CTTTGCTGGGCCTTTTTCCC | ACCTGAAATGCAGGGCTACT | 123 | FAM | BHQ-1 |
| **Homology Arm assays** |  |  |  |  |  |  |
| AAVS1 5' HA | CCTGGACACCCCGTTCTCCTGTGG | CTCTCCATCCTCTTGCTTTCTT | GGAGTGAGAGGTGACCCGA | 69 | FAM | BHQ-1 |
| RPLP0 5'HA | CTCCATGTTCCCAAAGGCCC | GGTTCGTGTGCTAGTTAGATG | AGGCGGAACAGAATAGGA | 101 | ROX | BHQ-2 |
| CHD1 3'HA | AAAGCAGAACCACCCAATT | GAGGCAAATCTGTTATTTAGAT AGG | GTGTAGTCTAAGCAGTTTAGCG | 130 | ROX | BHQ-2 |
| TRAC HA | CACCAAAGCTGCCCTTACCT | CGCCTTCAACAACAGCATTA | CCATTCCTGAAGCAAGGAAAC | 101 | ATTO550 | BHQ-2 |
| **Reference Assays** |  |  |  |  |  |  |
| AFF3 | AACAACTCTTTCTGTCCCCCT | CACCTAGCATGTGTGGCATT | GCAGATCCAGGTCGTTGAAG | 70 | HEX | BHQ-1 |
| NAGK | TGTTGCCCGAGATTGACCCGGT | TGGGCAGACACATCGTAGCA | CACCTTCACTCCCACCTCAAC | 66 | HEX | BHQ-1 |
| SarsCov2 | ACCCCGCATTACGTTTGGTGGACC | GACCCCAAAATCAGCGAAAT | TCTGGTTACTGCCAGTTGAATCTG | 72 | ATTO550 | BHQ-2 |
| RPP30 | TCTGACCTGAAGGCTCTGCGCG | GATTTGGACCTGCGAGCG | GCGGCTGTCTCCACAAGT | 62 | ATTO590 | 3IAbRQSp |
| TRAC Internal control | CGGTCTCAATATAACGACC | CTCTCTTTTGAAGCAGGCC | AGTGTCTGTTTTGACACGATCTGTA CTC | 551 | FAM | BHQ-1 |

HDRT: Homology Directed Repair Template, FWD: forward, REV: reverse, freq.: Frequency, dPCR: digital PCR, BHQ1: Black Hole Quencher 1, BHQ2: Black Hole Quencher 2, 3IAbRQSp: 3'Iowa Black RQ-Sp, 3IABkFQ: 3'Iowa Black FQ-Sp

**Supplementary Table 2. Formulas for calculating performance measures**

|  | Formula | Legend |
| --- | --- | --- |
| Rain | $\text{Rain (\%)} = 1 - \left( \frac{\text{cloud with rain}}{\text{cloud without rain}} \right) \times 100$ | *Cloud with rain*: clearly positive partitions/droplets + partitions/droplets between positive and negative clusters<br>*Cloud without rain*: positive partitions/droplets clearly separated from negative clusters |
| Quantitative accuracy | $\text{Accuracy (\%)} = \frac{\text{measured result}}{\text{expected result}}$ | expected result = assigned copy number of the reference material, determined by flow cytometry-based estimation |
| Limit of blank (LoB) | LoB = mean of all negative controls + 1,65 × standard deviation | According to the CLSI guideline EP17-A2 for establishing limits of detection and limits of blank at a 95% confidence level. |
| Coefficient of variance | $\text{CoV (\%)} = \frac{\text{standard deviation of results}}{\text{mean result of repeatedly measured sample}} \times 100$ | Intra Assay = same sample measured with the same assay on the same plate<br>Inter Assay = same sample measured with the same assay on different plates |
| Diagnostic sensitivity | $\text{sensitivity (\%)} = \frac{\text{true positives}}{(\text{true positives} + \text{false negatives})}$ | |
| Diagnostic specificity | $\text{specificity (\%)} = \frac{\text{true negatives}}{(\text{true negatives} + \text{false positives})}$ | |
| Efficiency of Transgene, Out/In_In/Out and Linkage Assays | $\text{Efficiency} = \frac{\text{Knock in of sample and Gene block}}{\text{Knock in of Gene block only} + \text{Knock in of sample only}}$ | Knock-in values represent measured copy numbers determined by dPCR |
| Efficiency of Homology Arm Assays | $\text{Efficiency} = \frac{H}{(T + r(\text{wt}) \times E)}$ | H = Homology Arm copy number<br>T = transgene copy number<br>E = endogenous control copy number<br>r(wt) = calibration factor for cell line CNV -><br>$r(\text{wt}) = \frac{H(\text{wt} - \text{only})}{E(\text{wt} - \text{only})}$ |
| Accuracy (dilution) | $\text{Accuracy} = \frac{\text{measured result}}{\text{measured result in undiluted sample}} \times 100$ | Accuracy was calculated as the ratio of the measured result in diluted samples relative to the undiluted sample. |
| Limit of Detection (LoD) | LoD = LoB + 1.65 × SD<sub>low concentration sample</sub> | LoD was defined as the lowest target concentration at which ≥95% of reactions were positive, according to CLSI EP17-A2. |

**Supplementary Table 3. Sequences of transgene components and gene block**

|  |
| --- |
| **AAVS1 gRNA** |
| GGGGCCACTAGGGACAGGAT |
| **CH1 gRNA** |
| GAATAGTCCGTGGTCTTCCA |
| **RPLP0 gRNA** |
| GCCGTGATGCCCAGGGAAGA |
| **TRAC gRNA** |
| GAGAATCAAAATCGGTGAAT |
| **AAVS1 5'homology arm (399bp)** |
| ACCGTTTTTCTGGACAACCCCAAAGTACCCCGTCTCCCTGGCTTTAGCCACCTCTCCATCCTCTTGCTTTCTTTGCCTGGACACCCCGTTCTCCTGTGGATTCGGGTCACCTCTCACTCCTTTCATTTGGGCAGCTCCCCTACCCCCCTTACCTCTCTAGTCTGTGCTAGCTCTTCCAGCCCCCTGTCATGGCATCTTCCAGGGGTCCGAGAGCTCAGCTAGTCTTCTTCCTCCAACCCGGGCCCCTATGTCCACTTCAGGACAGCATGTTTGCTGCCTCCAGGGATCCTGTGTCCCCGAGCTGGGACCACCTTATATTCCCAGGGCCGGTTAATGTGGCTCTGGTTCTGGGTACTTTTATCTGTCCCCTCCACCCCACAGTGGGGCCACTAGGGACAG |
| **AAVS1 3'homology arm (400bp)** |
| GATTGGTGACAGAAAAGCCCCATCCTTAGGCCTCCTCCTTCCTAGTCTCCTGATATTGGGTCTAACCCCCACCTCCTGTTAGGCAGATTCCTTATCTGGTGACACACCCCCATTTCCT GGAGCCATCTCTCTCCTTGCCAGAACCTCTAAGGTTTGCTTACGATGGAGCCAGAGAGGATCCTGGGAGGGAGAGCTTGGCAGGGGGTGGGAGGGAAGGGGGGGATGCGTGA CCTGCCCGGTTCTCAGTGGCCACCCTGCGCTACCCTCTCCCAGAACCTGAGCTGCTCTGACGCGGCCGTCTGGTGCGTTTCACTGATCCTGGTGCTGCAGCTTCCTTACACTTCCC AAGAGGAGAAGCAGTTTGGAAAAACAAAATCAGAATAAGTTGGTCCTGAGTTCT |
| **RPLP0 5'homology arm (400bp)** |
| ACAGGAGCGCTATCCGCGGTTTCTGATTGGCTACTTTGTTCGCATTATAAAAGGCACGCGCGGGCGCGAGGCCCTTCTCTCGCCAGGCGTCCTCGTGGAAGGTTCGTGTGCTAGT TAGATGGGCGCCAGGGGTCGCCGGCGGGAAGCATGGAGGGGTCTTTGGGGGCCTTTGGGAACATGGAGTCCTATTCTGTTCCGCCTGGGGCCTCGGTGGCGGCTTGCACGCCC CGAGATGACGGCCGCTGCCCTAGGCAGGGCCGGCGGGCGATTGCGCGTGTCCTGCTCCTCTTAGGCCCGGGACCGCGGGATGGGTGTCGGCGTGACCAGGCCTGAGCTCCCT GTCTCTCCTCAGTGACATCGTCTTTAAACCCTGCGTGGCAATCCCTGACGCACCGCCGTG |
| **RPLP0 3'homology arm (400bp)** |
| GGCGACCTGGAAGTCCAACACTCTTCTTAAGATCATCGTAAGTGCAAGGTGGGTGCGCTCTGCTCTTCATGTTGCCCCAGCGCAAATAGGGACAGTCAGCTGCTATGTGCTGAGG GTCTACTCACACCGGCTACTGAATTAGGCCATTTTTGGGAAAATACTGTTTAGTTAAACAATTTCTGAGATAGGTCCTTCTGTTGCAGATAAACGGGCTCAGGCAAGTTAAGTGG GTCTCTAAGATGACAGCATTCTGATCCAGGTCTGTCTGGCTTCTAAAAGAGCGCGCTTTATACTTTTATTTTTAAACGGAGTCTCGCTCTGTTGCCAGGCTGGAGTGCAGTGGCT TGATCTTGGCTTACTGCAACCTCCGCTTCCAGGTTCAAGCGATTCTCTCG |
| **CHD1 5'homology arm (400bp)** |
| TTATTGATCAATGTTTCTTCACAGTTCTGATCTCCTCAGGAAAAATGCCTAGAGGTTATGTTGCTGCGTCAGTAGAATAACTTTTGGAGGCTTTTGATATAATGTCTGATTACCTTTT AAATCAAAATATCCACCAGAGGTATATATTTTTCTCTGTTTGCTCTTTAATTTTGTTTATGGAATGTTTTTGATACATTTAATAACATTTTTAAGATTCCGACCATCTTTTATAATGGTT TTTGCTCTTAAGATACAAGCTGTTTTTACACTAAGGTTGGGTAAATATTTTCCAATTCCAGATTAGTTTCAATGGAAGTGAAGGGAGGCGCAGTAGAAGTAGGAGATACTCTGGAT CTGATAGTGATTCCATCTCAGAAGGGAAAAGGCCAAAGAACCGTGG |
| **CHD1 3'homology arm (400bp)** |
| AAGACCACGGACTATTCCTCGGGAGAATATTAAAGGATTTAGTGATGCAGAAATTAGGCGGTAAGGTGTTATAAGACAATCGCTTCAACTTGGTTTTAGTAGAGTGTGAAACTAA AATGTAAAAACGCCGTAATTTTCAAAAATTGTGATGAAGTTTTAGAAGAGGCAAATCTGTTATTTGAGTAGGAATTGGGTGGTTCTGCTTTTGGGTATTTGTTATAACTGAACAAA ACAAGTTTGGTGTCTAATCTTTGCATGTTATTTATTTATCGCTAAACTGCTTAGACTACACAATTTTTTTTATTAATTTGACTCAATAGTGATCCTTTCGAATAATAAAAGTAGTTTTG ATGCATTTGATTTTTAAAAATTCTCACGGTGTTTATGACTCCCAAAGTTA |
| TRAC 5' homology arm (400bp) |
| CCTATTAAATAAAAGAATAAGCAGTATTATTAAGTAGCCCTGCATTTCAGGTTTCCTTGAGTGGCAGGCCAGGCCTGGCCGTGAACGTTCACTGAAATCATGGCCTCTTGGCCAAG ATTGATAGCTTGTGCCTGTCCCTGAGTCCCAGTCCATCACGAGCAGCTGGTTTCTAAGATGCTATTTCCCGTATAAAGCATGAGACCGTGACTTGCCAGCCCCACAGAGCCCCGCC CTTGTCCATCACTGGCATCTGGACTCCAGCCTGGGTTGGGGCAAAGAGGGAAATGAGATCATGTCCTAACCCTGATCCTCTTGTCCCACAGATATCCAGAACCCTGACCCTGCCGT GTACCAGCTGAGAGACTCTAAATCCAGTGACAAGTCTGTCTGCCTATTCACC |
| TRAC 3' homology arm (400bp) |
| ATTTTGATTCTCAAACAAATGTGTCACAAAGTAAGGATTCTGATGTGTATATCACAGACAAAACTGTGCTAGACATGAGGTCTATGGACTTCAAGAGCAACAGTGCTGTGGCCTG GAGCAACAAATCTGACTTTGCATGTGCAAACGCCTTCAACAACAGCATTATTCCAGAAGACACCTTCTTCCCCAGCCCAGGTAAGGGCAGCTTTGGTGCCTTCGCAGGCTGTTTCC TTGCTTCAGGAATGGCCAGGTTCTGCCCAGAGCTCTGGTCAATGATGTCTAAAACTCCTCTGATTGGTGGTCTCGGCCTTATCCATTGCCACCAAAACCCTCTTTTTACTAAGAAAC AGTGAGCCTTGTTCTGGCAGTCCAGAGAATGACACGGGAAAAAAGCAGATG |
| **Ef1α(s) promoter 239bp** |
| GCTCCGGTGCCCGTCAGTGGGCAGAGCGCACATCGCCCACAGTCCCCGAGAAGTTGGGGGGAGGGGTCGGCAATTGAACCGGTGCCTAGAGAAGGTGGCGCGGGGTAAACTG GGAAAGTGATGTCGTGTACTGGCTCCGCCTTTTTCCCGAGGGTGGGGGAGAACCGTATATAAGTGCAGTAGTCGCCGTGAACGTTCTTTTTCGCAACGGGTTTGCCGCCAGAAC ACAGGCCGCCACC |
| **CXCL10 (294bp)** |
| ATGAACCAGACCGCCATCCTGATCTGCTGCCTGATCTTCCTGACCCTGAGCGGCATCCAGGGAGTCCCCCTGTCCAGAACAGTGCGGTGCACCTGTATCAGCATCTCTAATCAACC TGTGAACCCAAGAAGCCTGGAAAAGCTCGAGATCATCCCTGCTTCTCAGTTCTGCCCTAGAGTGGAAATCATCGCCACAATGAAAAAGAAGGGCGAGAAGAGATGTCTGAACCC CGAGAGCAAGGCCATTAAGAACCTGCTGAAGGCCGTGTCCAAAGAGATGAGCAAGCGGAGCCCT |
| **CXCL11 (282bp)** |
| ATGAGTGTGAAGGGCATGGCTATAGCCTTGGCTGTGATATTGTGTGCTACAGTTGTTCAAGGCTTCCCCATGTTCAAAAGAGGACGCTGTCTTTGCATAGGCCCTGGGGTAAAAG CAGTGAAAGTGGCAGATATTGAGAAAGCCTCCATAATGTACCCAAGTAACAACTGTGACAAAATAGAAGTGATTATTACCCTGAAAGAAAATAAAGGACAACGATGCCTAAATCC CAAATCGAAGCAAGCAAGGCTTATAATCAAAAAAGTTGAAAGAAAGAATTTT |
| **IFNG (498bp)** |
| ATGAAGTACACCTCTTACATTCTGGCATTCCAATTGTGCATTGTGCTGGGAAGCCTCGGGTGCTATTGTCAAGATCCCTACGTGAAAGAGGCTGAGAATCTGAAGAAGTACTTCAA CGCCGGGCACAGTGACGTCGCTGACAACGGTACACTGTTCTTGGGAATCTTAAAGAACTGGAAGGAAGAAAGCGATCGAAAGATCATGCAATCCCAGATCGTGAGCTTCTATTTT AAGCTGTTCAAGAACTTCAAGGACGACCAATCTATACAGAAATCAGTCGAGACAATCAAAGAGGATATGAACGTGAAGTTCTTTAACTCTAATAAGAAGAAGAGGGACGATTTTG AGAAACTTACAAACTACAGCGTCACGGATCTCAACGTACAGCGGAAGGCGATCCACGAGCTTATTCAGGTCATGGCCGAGTTGTCCCCGGCTGCGAAGACTGGCAAACGGAAGC GTTCACAAATGCTTTTCAGAGGGAGACGAGCGTCACAA |
| **GFP (714bp)** |
| GTGAGCAAGGGCGAGGAGCTGTTCACCGGGGTGGTGCCCATCCTGGTCGAGCTGGACGGCGACGTAAACGGCCACAAGTTCAGCGTGTCCGGCGAGGGCGAGGGCGATGCC ACCTACGGCAAGCTGACCCTGAAGTTCATCTGCACCACCGGCAAGCTGCCCGTGCCCTGGCCCACCCTCGTGACCACCCTGACCTACGGCGTGCAGTGCTTCAGCCGCTACCCCG ACCACATGAAGCAGCACGACTTCTTCAAGTCCGCCATGCCCGAAGGCTACGTCCAGGAGCGCACCATCTTCTTCAAGGACGACGGCAACTACAAGACCCGCGCCGAGGTGAAGT TCGAGGGCGACACCCTGGTGAACCGCATCGAGCTGAAGGGCATCGACTTCAAGGAGGACGGCAACATCCTGGGGCACAAGCTGGAGTACAACTACAACAGCCACAACGTCTAT ATCATGGCCGACAAGCAGAAGAACGGCATCAAGGTGAACTTCAAGATCCGCCACAACATCGAGGACGGCAGCGTGCAGCTCGCCGACCACTACCAGCAGAACACCCCCATCGG CGACGGCCCCGTGCTGCTGCCCGACAACCACTACCTGAGCACCCAGTCCGCCCTGAGCAAAGACCCCAACGAGAAGCGCGATCACATGGTCCTGCTGGAGTTCGTGACCGCCGC CGGGATCACTCTCGGCATGGACGAGCTGTACAAG |
| **P2A (66bp)** |
| GGCAGTGGAGCAACCAACTTCAGCCTGCTGAAGCAGGCAGGCGACGTGGAAGAAAATCCAGGACCT |
| **Q8 reporter tag (417bp)** |
| ATGGGATTGGTTCGCCGAGGAGCGCGAGCAGGTCCCCGGATGCCCCGCGGATGGACTGCCCTGTGCCTGCTTTCCCTCCTTCCAAGCGGCTTCATGGCAGAATTGCCAACTCAA<br>GGTACTTTCTCAAATGTTTCCACGAACGTTAGTCCAGCTAAACCCACTACAACACCTGCACCTAGACCCCCGACGCCAGCCCCTACTATAGCTTCTCAACCACTGAGTCTTCGGCCA<br>GAGGCATGCAGGCCCGCAGCAGGTGGGGCAGTGCACACAAGGGGTCTCGATTTTGCTTGTGACATTTATATTTGGGCTCCTTTGGCAGGGACCTGTGGGGTACTTCTGCTGAGT<br>CTGGTGATCACGTTGTATTGTAACCACAGGAATAGGCGGCGGGTTTGTAAATGTCCACGACCGGTTGTCTGA |
| **sPA (49bp)** |
| AATAAAAGATCTTTATTTTCATTAGATCTGTGTGTTGGTTTTTTGTGTG |
| **CMV (508bp)** |
| CGTTACATAACTTACGGTAAATGGCCCGCCTGGCTGACCGCCCAACGACCCCCGCCCATTGACGTCAATAATGACGTATGTTCCCATAGTAACGCCAATAGGGACTTTCCATTGAC<br>GTCAATGGGTGGAGTATTTACGGTAAACTGCCCACTTGGCAGTACATCAAGTGTATCATATGCCAAGTACGCCCCCTATTGACGTCAATGACGGTAAATGGCCCGCCTGGCATTA<br>TGCCCAGTACATGACCTTATGGGACTTTCCTACTTGGCAGTACATCTACGTATTAGTCATCGCTATTACCATGGTGATGCGGTTTTGGCAGTACATCAATGGGCGTGGATAGCGGT<br>TTGACTCACGGGGATTTCCAAGTCTCCACCCCATTGACGTCAATGGGAGTTTGTTTTGGCACCAAAATCAACGGGACTTTCCAAAATGTCGTAACAACTCCGCCCCATTGACGCAA<br>ATGGGCGGTAGGCGTGTACGGTGGGAGGTCTATATAAGCAGAGCT |
| **P2A - CD19 targeting CAR - IgG4 linker - IgG1 Hinge - CD28-CDZeta - bGH poly A signal** |
| GCCACCAACTTCTCTCTTTTGAAGCAGGCCGGAGATGTGGAAGAAAATCCTGGGCCCATGGCTCTTCCTGTGACTGCCCTTCTGCTGCCCCTGGCTCTCCTGCTCCATGCTGCCCG<br>GCCAGACATCCAGATGACACAGACAACCAGCAGCCTCAGCGCCAGCCTGGGGGACAGAGTGACCATTAGCTGCCGGGCCTCTCAGGACATCAGCAAATACCTGAACTGGTACCA<br>GCAGAAACCAGATGGCACTGTCAAGCTGCTGATTTACCACACCTCCAGGCTCCACAGCGGCGTGCCCAGTCGCTTCAGCGGCAGTGGGAGCGGGACAGATTATTCCCTCACAAT<br>CTCCAACCTGGAGCAGGAAGATATTGCCACATACTTCTGCCAGCAAGGCAACACCCTGCCATACACATTTGGAGGCGGCACCAAATTGGAGATCACCGGCGGTGGTGGATCTGG<br>AGGAGGGGGCAGCGGAGGTGGCGGCTCTGAGGTGAAACTGCAGGAGAGTGGCCCTGGCCTGGTGGCTCCCAGCCAGAGCCTTTCTGTCACCTGCACCGTGTCTGGGGTGTCC<br>CTGCCTGACTATGGAGTCTCCTGGATCCGGCAGCCTCCAAGAAAAGGACTGGAATGGCTGGGCGTCATCTGGGGAAGTGAGACCACCTACTATAATTCAGCCCTCAAGTCCCGG<br>CTCACCATCATTAAGGACAACTCCAAATCCCAGGTGTTCCTGAAGATGAATTCTCTCCAGACTGATGACACAGCCATCTACTACTGTGCCAAGCATTATTATTACGGCGGGTCCTAT<br>GCCATGGACTACTGGGGCCAGGGGACCAGTGTCACTGTTTCTTCTGAAAGCAAGTATGGCCCCCCATGCCCTCCCTGCCCCGGACAGCCGCGGGAGCCCCAGGTCTACACCCTCC<br>CACCCTCCAGAGATGAGCTGACCAAGAACCAGGTTTCACTGACATGCCTGGTGAAGGGCTTCTACCCCTCTGACATTGCTGTGGAGTGGGAGAGCAATGGGCAGCCAGAGAACA<br>ACTACAAGACCACACCTCCTGTCCTGGACAGCGACGGCTCCTTCTTCCTCTACAGTAAGCTCACTGTGGACAAGAGCCGCTGGCAGCAGGGCAATGTCTTCTCCTGCAGCGTGAT<br>GCACGAGGCCCTGCACAATGCCTATACCCAGAAGTCACTCTCTCTGAGCCCTGGAAAGAAGGATCCCAAGTTTTGGGTCCTCGTGGTGGTCGGGGGTGTGCTGGCCTGCTACAG<br>CTTGCTGGTCACAGTGGCCTTCATCATCTTCTGGGTGCGCTCCAAGAGGAGCCGGCTGCTTCACAGTGATTACATGAACATGACCCCCAGGAGGCCAGGACCCACCAGGAAGCA<br>CTACCAGCCCTACGCTCCCCCGCGGGACTTTGCTGCTTACCGCAGCAGGGTCAAATTTTCTAGATCTGCAGATGCGCCGGCCTATCAACAAGGCCAGAACCAGCTCTATAACGAG<br>CTCAATCTAGGACGAAGAGAGGAGTACGATGTTTTGGACAAGAGACGTGGCCGGGACCCTGAGATGGGGGGAAAGCCGAGAAGGAAGAACCCTCAGGAAGGCCTGTACAATG<br>AACTGCAGAAAGATAAGATGGCGGAGGCCTACAGTGAGATTGGGATGAAAGGCGAGCGCCGGAGGGGCAAGGGGCACGATGGCCTTTACCAGGGTCTCAGTACAGCCACCAA<br>GGACACCTACGACGCCCTTCACATGCAGGCCCTGCCCCCTCGCTAATAAGAATTCTAACTAGAGCTCGCTGATCAGCCTCGACTGTGCCTTCTAGTTGCCAGCCATCTGTTGTTTGC<br>CCCTCCCCCGTGCCTTCCTTGACCCTGGAAGGTGCCACTCCCACTGTCCTTTCCTAATAAAATGAGGAAATTGCATCGCATTGTCTGAGTAGGTGTCATTCTATTCTGGGGGGTGG<br>GGTGGGGCAGGACAGCAAGGGGGAGGATTGGGAAGAGAATAGCAGGCATGCTGGGGAGCGGCCGCGTTAA |
| **Geneblock (1892bp)** |
| CTGATAAGCGCTATCCGCGGTTTCTCTGATAATGGACCCCAAAATCAGCGAAATGCACCCCGCATTACGTTTGGTGGACCCTCAGATTCAACTGGCAGTAACCAGAATGTCTAGA<br>GAGAGGCGACGTGGAAGAAAATCCAGGACCTATGGGATTGGTTCGCCGAGGAGCGCGAGCAGGTCCCCGGATGCCCCGCGGATGGACTGCCCTGTGCCTGCTTTCCCTCCTTC<br>CAAGCGGCTTCATGGCAGAATTGCCAACTCAAGGTACTTTCTCAAATGTTTCCACGAACGTTAGTAGACCCGCGCCGAGGTGAAGTTCGAGGGCGACACCCTGGTGAACCGCATC<br>GAGCTGAAGGGCATCGACTTCAAGGAGGACGGCAACATCCTGGGGCACAAGCTGGAGTACAACTACAACAGCCACAACGTCTATATCATGGCCGACAAGCAGAAGAACGGCAT<br>CAAGGTGAACTTCAAGATCCGCCACAACATCGAGGACGGCAGCGTGCAGCTCGCCGACCACTACCAGCAGAACACCCCCATCGGCGACGGCCCCGTGTTAGCCACCTCTCCATC<br>CTCTTGCTTTCTTTGCCTGGACACCCCGTTCTCCTGTGGATTCGGGTCACCTCTCACTCCAATAAAAGATCTTTATTTTCATTAGATCTGTGTGTTGGTTTTTTGTGTGTGTGTGCTAC<br>AGTTGTTCAAGGCTTCCCCATGTTCAAAAGAGGACGCTGTCTTTGCATAGGCCCTGGGGTAAAAGCAGTGAAAGTGGCAGATATTGAGAAAGCCTCCATAATGTACCCAAGTACT<br>TCAGGACAGCATGTTTGCTGCGACCACCTTATATTCCCAGGGCCGGTTAATGTGGCTCTGGTTCTGGGTACTTTTATCTGTCCCCTCCACCCCACAGTGGGGCCACTAGGGACAGG<br>ATTGGTGACAGAAAAGCCCCATCCTTAGGCCTCCTCCTTCCTAGTCTCCTGATATTGGGTCTAACCCCCACCTCCTGAAGTTGGTCCTGAGTTCTAACTTTGGCTCTTCACCTTTCTA<br>GTCCCCAATTTATATTGTTCCTCCGTGCGTCAGTTTTACCTGTGAGATAAGGCCAGTAGCCCCCATGCCGTCTTCACTCGCTGGGTTCCCTTTTCCCGACGGATGTCTCCCTTGCGTC<br>CCGCCTCCCCTTCTTGTAGCCTTCCTTCGCGACCCTACTTAAAGGCGGCTTTCCGTTGGCTGGAGTGGCAGCGATATTATCCAATGGTTGCCTGTATTCGTTCAGCTTTGTCTGACG<br>GGCGATGGCGCAGCCAATAGACAGGATTTGGTAGAGACAAGGTCTCACTATGTTGCCTAAGCTGGTCTTGAACTCCTGGGCTCAAGTTATCTTCCTGCCTTGGCCTCCCAGTCAT<br>GCTGGGATTACAGGCATGAGCTGCTGCACTCAGCCACATTTTTCTTTTATGACTGCTGCCTGATCTTCCTGACCCTGAGCGGCATCCAGGGAGTCCCCCTGTCCAGAACAGTGCGG<br>TGCACCTGTATCAGCATCTCTAATCAACCTGTGAACCCAAGAAGCCTGGAAAAGCTCGAGATCATCCCTGCTTCTCAGTTCTGCCCTAGAGTGGAAATCATCGCCACAATGAAAAA<br>GAAGGGCGAGAAGAGGAACTGGAAGGAAGAAAGCGATCGAAAGATCATGCAATCCCAGATCGTGAGCTTCTATTTTAAGCTGTTCAAGAACTTCAAGGACGACCAATCTATACA<br>GAAACATAACTTACGGTAAATGGCCCGCCTGGTAGAGAAGGTGGCGCGGGGTAAACTGGGAAAGTGATGTCGTGTACTGGCTCCGCCTTTTTCCCGAGGGTGGGGGAGAACCG<br>TATATAAGTGCAGTAGTCAGGTTCAAGCGATTCTCCTGCCTGATA |
| **Included:** SarsCov2 Nucleocapsid detection assay, In-Out: P2A-AAVS1, AAVS1 5’HA, CXCL11 transgene, AAVS1 5’ target locus, AAVS1 cut site, AAVS1 3’ target locus, Out/In AAVS1-EF1α(s), Out/In RPLP0-EF1α(s), Out/In CHD1-EF1α(s), RPLP0 target locus, CHD1 target locus, CXCL10 transgene, IFNG transgene. |
| **SarsCov2 NucleoCapsid sequence (72bp)** |
| GACCCCAAAATCAGCGAAATGCACCCCGCATTACGTTTGGTGGACCCTCAGATTCAACTGGCAGTAACCAGA |

Supplementary Table 4. Equipment and Consumables used

| Item | Catalogue n° | Provider |
| --- | --- | --- |
| **Equipment** |  |  |
| 4D-Nucleofector® core unit |  | Lonza Bioscience |
| 4D-Nucleofector® X Unit |  | Lonza Bioscience |
| Agilent 2100 Bioanalyzer |  | Agilent Technologies |
| C1000 Touch Thermal cycler |  | Bio-Rad |
| Centrifuge 5415R |  | Eppendorf GmbH |
| Centrifuge 5424 |  | Eppendorf GmbH |
| Centrifuge 5427R |  | Eppendorf GmbH |
| Centrifuge 5810 R |  | Eppendorf GmbH |
| Covaris S220 Focused-ultrasonicator |  | Covaris |
| Duomax 1030, platform shaker |  | Heidolph instruments |
| Electronic Balance ABS 80-4 |  | Kern & Sohn GmbH |
| Erlenmeyer flasks |  | Thermo Scientific |
| FlowCytometer LSR-Fortessa X-20 |  | BD Biosciences |
| Fortessa Aria Cell sorter |  | BD Bioscience |
| Fortessa X-20 |  | BD Bioscience |
| Freezer (-20°C) LCv4010 |  | Carl Roth |
| Freezer (-80°C), HeraFreeze T series HFU400TV63 |  | Thermo Scientific |
| Ice machine Manitowoc SOTTO |  | Manitowoc |
| Incubator HERAcell 240i CO2 |  | Thermo Scientific |
| Laminar Airflow Bench HERA safe 2020 KSP18 |  | Thermo Scientific |
| Magnetic Shaker RH basic |  | IKA Laboratoy Equipment |
| Magnetic Rack (for 1.5-2mL tubes) |  | Qiagen |
| Manual system microscope Olympus BX43 |  | Olympus |
| Microscope Axio Vert.A1 |  | Zeiss |
| Millipore Barnstead MicroPure |  | Thermo Scientific |
| NanoDrop 2000 |  | Thermo Scientific |
| Neubauer counting chamber |  | Brand |
| NextSeq 500/550 Sequencing System |  | Illumina |
| PCR workstation Pro |  | VWR Peqlab |
| pH-meter Five Easy Le409 |  | Mettler Toledo |
| Pipette filler pipetusR |  | Hirschmann Labortechnik |
| Pipette multipipetteR stream |  | Eppendorf GmbH |
| Pipettes (2.5 - 1000 µl) |  | Eppendorf GmbH |
| PX1 PCR Plate Sealer |  | Bio-Rad |
| QuadroMACS |  | Miltenyi |
| Qubit fluorometer |  | Thermo Scientific |
| QIAcuity Digital PCR System (6-plex) |  | Qiagen |
| QX200 & QX600 Droplet Digital PCR System |  | Bio-Rad |
| Shandon Excelsior ES |  | Thermo Scientific |
| Spectrophotometer EPOCH |  | BioTek Instruments |
| StepOnePlus Real-Time PCR system |  | Thermo Scientific |
| Suction pump AC02 |  | Carl Roth |
| Table centrifuge mini star silverline |  | VWR |
| ThermoMixer C |  | Eppendorf GmbH |
| Vortexer Reax top |  | Heidolph instruments |
| Vortexer VWRR Galaxy Mini Star |  | VWR International bvba |
| Waterbath GFL 1086 |  | GFL Technology |
| **General consumables** |  |  |
| 8-Well PCR tube strips plus 8 domed caps | strips: 72.985.002, caps:<br>65.989.002 | Sarstedt |
| 96-Well Plate Advanced TC (flat bottom) | 655983 | Greiner Bio-one |
| Biosphere® Filter Tip 10 µl | 70.1130.210 | Sarstedt |
| Biosphere® Filter Tip 1000 µl | 70.762.211 | Sarstedt |
| Biosphere® Filter Tip 100 µl | 70.760.212 | Sarstedt |
| Cell culture multiwell plate, 6 well, PS, clear, sterile | 657160 | Greiner Bio-one |
| Cell culture plates CELLSTAR®, sterile, white-96-well plates | KL43.1 | Roth |
| CELLSTAR®, TC, lid with condensation rings, sterile | 655180 | Greiner Bio-one |
| CryoPure Tube 1.6 ml yellow | 72.380.004 | Sarstedt |
| Disposable needles Sterican® long bevel facet, 0.30x12 mm | 4656300 | B.Braun |
| Disposable Syringe, Luer, 1 ml | CH030001L | Charina |
| FACS tubes | 352052 | BD |
| Falcon® 10 ml Serological Pipet | 357551 | Corning |
| Falcon® 12-well Clear Flat Bottom TC-treated Multiwell Cell Culture Plate | 353043 | Corning |
| Falcon® 12-well Clear Flat Bottom TC-treated Multiwell Cell Culture Plate | 353043 | Corning |
| Falcon® 15 ml High Clarity PP Centrifuge Tube | 352096 | Corning |
| Falcon® 2 ml Serological Pipet | 357507 | Corning |
| Falcon® 24-well Clear Flat Bottom TC-treated Multiwell Cell Culture Plate | 353047 | Corning |
| Falcon® 24-well Clear Flat Bottom TC-treated Multiwell Cell Culture Plate | 353047 | Corning |
| Falcon® 25 ml Serological Pipet | 357525 | Corning |
| Falcon® 35 mm TC-treated Easy-Grip Style Cell Culture Dish | 353001 | Corning |
| Falcon® 40 µm Cell Strainer | 352340 | Corning |
| Falcon® 48-well Clear Flat Bottom TC-treated Multiwell Cell Culture Plate | 353078 | Corning |
| Falcon® 5 ml Round Bottom Polystyrene Test Tube | 352052 | Corning |
| Falcon® 5 ml Round Bottom Polystyrene Test Tube, with Cell Strainer Snap Cap | 352235 | Corning |
| Falcon® 5 ml Serological Pipet | 357543 | Corning |
| Falcon® 50 ml High Clarity PP Centrifuge Tube | 352070 | Corning |
| FrameStar Fast Plate 96-well semi skirted | 4ti-1200 | 4titude |
| Injekt Solo-2-piece single-use syringe, 10 ml, Luer Lock | 201235 | B.Braun |
| MACS LS Columns | 130-042-401 | Miltenyi Biotec |
| MACS MS Columns | 130-042-201 | Miltenyi Biotec |
| Pasteurpipette glass, 145mm | 500635 | Brand |
| Pasteurpipette glass, 230mm | 500636 | Brand |
| Reagent Reservoirs, 25 ml | EKT8.1 | Carl Roth |
| Rotilab®-syringe filters, CA, sterile, 0.45µm | KC71.1 | Carl Roth |
| Spitzen Filter Surphob 100 µl (10x96) | VT0230 | Biozym |
| Spitzen Filter Surphob 10 µl lang (10x96) | VT0200 | Biozym |
| Spitzen Filter Surphob 1,250 µl (10x96) | VT0270 | Biozym |
| Stericup | CT92.1 | Carl Roth |
| Sterile, Stainless Steel, Premium Disposable Scalpel, 11PA | 03025 | Razormed |
| Vasco® Nitril blue glove S-L | 9205518 | B.Braun |
| **Chemicals and reagents (general)** |  |  |
| 0.05% Trypsin-EDTA (1x) | 25300-096 | Gibco |
| Albumin from bovine serum Fraktion V | 8076.3 | Carl Roth |
| Anti-human CD28 antibody | 302934 | Biolegend |
| Anti-PE microbeads | 130-105-639 | Invitrogen |
| CD3 antibody anti-human, pure-functional grade clone OKT3 | 130-093-387 | Miltenyi |
| CD8 microbeads, human - lyophilized | 130-097-057 | Miltenyi |
| CS&T Research Beads | 655050 | Becton Dickinson |
| Cut Smart Buffer | NEB#B6004 | New England Biolabs |
| D-Luciferin | 122799 | Perkin Elmer Inc. |
| Dimethyl sulfoxide (DMSO) | A994.1 | Carl Roth |
| DISPASE II | D4693-1G | Sigma Aldrich |
| Dnase I | A3778.0100 | AppliChem |
| Dynabeads Human T-Activator CD3/CD28 | 11131D | ThermoFisher |
| Dynabeads Mouse T-Activator CD3/CD28 | 114530 | ThermoFisher |
| Ethanol Rotipuran 99.8% p.a. | 2065.2 | Carl Roth |
| Fetal Bovine Serum (FBS) SUPERIOR | S0615-500ML | Merck |
| Ficoll Paque Plus | 17144002 | Cytiva |
| Flow-Set Pro Fluorospheres | A62492 | Beckman Coulter |
| FlowCheck Beads | A63493 | Beckman Coulter |
| FlowClean Cleaning Agent | A64669 | Beckman Coulter |
| Glycine Buffer ≥ 99% p.a. | 3908.3 | Carl Roth |
| H<sub>2</sub>O<sub>2</sub> (2N) | 339741 | Sigma Aldrich |
| Human IL-15, premium grade | 130-095-765 | Miltenyi |
| Human IL-7, premium grade | 130-095-362 | Miltenyi |
| Human Fc Receptor Blocking Solution | 422301 | Biolegend |
| IsoFlow Sheath Fluid | 8546859 | Beckman Coulter |
| KAPA HiFi HotStart Ready Mix | 7958927001 | Roche |
| L-Glutamine | 25020-081 | Life Technologies |
| LE Agarose | 240004 | Biozym |
| Live/Dead Fixable Near IR (780) Viability Kit, for 633 nm excitation | L34975 | ThermoFisher Scientific |
| Monensin | M5273-500MG | Sigma Aldrich |
| Nuclease-free water | 1097705 | Invitrogen |
| Penicillin-Streptomycin (10,000 U/mL) | 18140122 | Gibco |
| PhosSTOP | 4906837001 | Sigma Aldrich |
| Potassium chloride (KCl) ≥ 99.5% p.a. | 6781.1 | Carl Roth |
| Potassium dihydrogen phosphate (KH<sub>2</sub>PO<sub>4</sub>) ≥ 99% p.a. | 3904.1 | Carl Roth |
| Potassium hydrogen carbonate (KHCO<sub>3</sub>) ≥ 99% | X887.1 | Carl Roth |
| Precision Plus Protein Standards | 161-0374 | Bio-Rad |
| Q5 Polymerase | M0491L | New England Biolabs |
| Retronectin | T1008 | Takara |
| SDS Pellets ≥ 99.9% | CN30.3 | Carl Roth |
| Sodium butyrate | B5887-1g | Sigma Aldrich |
| TEMED ≥ 98.5% | 2367.1 | Carl Roth |
| Tris Bufferan® ≥ 99.9% p.a. | 4855.5 | Carl Roth |
| Trypan Blue Solution 0.4% | 15250061 | Gibco |
| Tween®20 | 9127.2 | Carl Roth |
| Vibrant™ Dio Cell-Labeling Solution | V22886 | ThermoFisher |
| β-Mercaptoethanol 99% p.a. | 4227 | Carl Roth |
| **Cell culture media** |  |  |
| RPMI Medium 1640 (1x) | 21875-034 | Life Technologies |
| DMEM (1x) Dulbecco's Modified Eagle Medium | 41966052 | Life Technologies |
| Opti-MEM | 31985062 | Life Technologies |
| Freezing medium | 10% DMSO, 90% fetal calf serum (FCS) |  |
| Tumor cell medium with DMEM | DMEM (1x), 10% FCS |  |
| T cell (human) medium | RPMI Medium 1640 (1x), 10% FCS, 2 mM L-Glutamin |  |
| **HiC workflow specific** |  |  |
| EpiTecT HiC Kit | 1076826 | Qiagen |
| QIAseq Library Quant Kit | 180501 | Qiagen |
| Agilent High Sensitivity DNA Kit | 5067-4626 | Agilent Technologies |
| NextSeq 500/550 v2.5 Sequencing Kit | 20024907 | Illumina |
| Formaldehyde solution (36-37%) | 252549 | Sigma-Aldrich |
| Tris-HCl, pH 7.5 (quenching buffer) | - | Carl Roth |
| Glycine (quenching reagent) | 3908.3 | Carl Roth |
| **HDRT-generation** |  |  |
| Q5 Hot Start High-Fidelity 2X Master Mix | M0494S | NEB |
| NEBuilder HiFi DNA Assembly Master Mix | E2621L | NEB |
| XL10-Gold Ultracompetent Cells | 200315 | Agilent |
| Kapa HiFi Hot Start ready mix | KK2602/7958935001 | Roche |
| AMPure XP, 60 mL | A63881 | Beckman Coulter |
| DynaMag2 | 12321D | Thermo Fisher Scientific |
| DpnI | R0176S | New England Biolabs |
| Effectene Transfection Reagent | 301425 | Qiagen |
| **CRISPR knock-in** |  |  |
| Alt-R® S.p. Cas9 Nuclease V3, 500 µg | 1081059 | IDT |
| Poly-L-glutaminsäure Natriumsalz | P4761-100MG | Sigma-Aldrich |
| SF Cell Line 4D-Nucleofector™ X Kit L | V4XC-2024 | Lonza |
| P3 Primary Cell 4D-Nucleofector™ X Kit L | V4XP-3024 | Lonza |
| **dPCR** |  |  |
| Pipet-Lite Multi Pipette L8-200XLS+ | 17013805 | Mettler-Toledo |
| Rainin Pipette Tips TR LTS 200 µL F 960A/10 | 17014963 | Mettler-Toledo |
| Eppendorf twin.tec® PCR Plate 96, semi-skirted, clean | 30128575 | Eppendorf |
| DNA LoBind® Tubes | 30108051 | Eppendorf |
| DG8 Cartridges for QX200 Droplet Generator | 186-4008 | Bio-Rad |
| DG8 Gaskets | 186-3009 | Bio-Rad |
| Droplet Generation Oil for Probes | 186-3005 | Bio-Rad |
| Droplet Reader Oil | 186-3004 | Bio-Rad |
| Pierceable Foil Heat Seals | 181-4040 | Bio-Rad |
| ddPCR Supermix for Probes | 186-3024 | Bio-Rad |
| ddPCR Multiplex Supermix | 120-01926 | Bio-Rad |
| QIAamp DNA Mini Kit (50) | 51304 | Qiagen |
| QIAcuity Probe PCR Kit (5 ml) | 250102 | Qiagen |
| QIAcuity Nanoplate 26k 24-well (10) | 250001 | Qiagen |
| QIAcuity Nanoplate 26k 8-well (10) | 250031 | Qiagen |
| QIAcuity Nanoplate 8.5k 24-well (10) | 250011 | Qiagen |
| QIAcuity Nanoplate 8.5k 96-well (10) | 250021 | Qiagen |
| Nanoplate Seals (11) | 250099 | Qiagen |
| Nanoplate Tray | 250098 | Qiagen |
| XbaI | R0145S | New England Biolabs |

**Supplementary Table 5. Antibodies used for flow cytometry and IF Staining**

| Antibodies | Clone | Dilution | Catalogue no. | Provider |
| --- | --- | --- | --- | --- |
| Anti-FC, Ax647 | polyclonal | 1:50 | 109-605-098 | Jackson Immuno Research |
| Invitrogen, CD34 Monoclonal Antibody (QBEND/10), PE | 537860 | 1:10 | #MA1-10205 | ThermoFisher Scientific |
| Live/Dead Fixable UV | not applicable | 1:100 | L34962 | Thermo Fisher Scientific |
| CD4, FITC | RPA-T4 | 1:400 | 357406 | Biolegend |
| CD4, BV421 | RPA-T4 | 1:50 | 317434 | Biolegend |
| CD8, PerCP | SK1 | 1:400 | 344707 | Biolegend |
| CD8, BV605 | SK1 | 1:400 | 301036 | Biolegend |
| CD3, PB | UCHT1 | 1:50 | 300442 | Biolegend |
| CD3, Ax700 | HIT3a | 1:400 | 300324 | Biolegend |
| CD3, FITC | HIT3a | 1:400 | 300306 | Biolegend |
| CD3, PerCP-Cy5.5 | HIT3a | 1:400 | 300328 | Biolegend |
| EGFRt (Cetuximab), PE | C225 | 1:100 | 537860 | Invitrogen |

**Supplementary Table 6. Software used**

| Software | Use | Provider |
| --- | --- | --- |
| DNA Folding Form (Mfold) | DNA accessibility analysis for primer design | UNAFold.org |
| EpiTect HiC Analysis Portal | Primary Hi-C QC & processing | Qiagen |
| FlowJo_v.10.6.2 | Analysis of flow cytometry data | FlowJo |
| GraphPad PRISM_v8 to v10 | Data analysis and graphical presentation | GraphPad |
| HiNT | CNV and translocation inference from Hi-C | Wang et al. |
| ICE Analysis | Gene knock-out quantification from Sanger data | Synthego |
| Illumina BaseSpace / Local Run Manager | Primary Sequencing data handling | Illumina |
| PrimerQuest Tool | Primer design and adjustment | IDT |
| Primer3Plus | Primer design and adjustment | Free software, Copyright (c) 2006, 2007 by Andreas Untergasser and Harm Nijveen |
| ProSort_v1.6 | FACS Sorter | Bio-Rad |
| QuantaSoft Analysis Pro_v2.2 | ddPCR Analysis | Bio-Rad |
| QX Manager Software_v 2.1 | ddPCR Analysis | Bio-Rad |
| QuiaCuity Software Suite_v1.7 | dPCR Analysis | Qiagen |
| SnapGene | Development of cloning strategies, new Plasmids | Dotmatics |
| StepOne Software_v2.3 | qRT-PCR set-up and analysis | Applied Biosystems |
